# Identification of a putative RocS homolog through phenotypic profiling of uncharacterized essential genes in *Streptococcus mutans*

**DOI:** 10.64898/2026.07.15.738700

**Authors:** Courtney E. Dover, Kipa Tamrakar, Bikash Dwivedi, Evan R. Roberts, Sangam Chudal, Shawn King, Emilio Soriano Chavez, Valérie de Crécy-Lagard, Robert C. Shields

## Abstract

Genome-wide viability catalogs produced by transposon sequencing (Tn-seq) and CRISPR interference (CRISPRi) have successfully mapped the essential genome of *Streptococcus mutans*. In this study, we combined predictive bioinformatics, conditional CRISPRi transcriptional silencing, transmission electron microscopy, transcriptomics, and genetic suppressor screens to investigate nine poorly characterized essential genes in *S. mutans*. From this screen, phenotypic and genetic analyses identified SMU_393 as a functional homolog of the pneumococcal chromosome segregation factor, RocS. Depletion of SMU_393 resulted in abnormal cell widening, hypersensitivity to DNA damage, and a significant subpopulation of anucleate cells. These phenotypes were bypassed by a spontaneous surface-exposed missense mutation (*dnaA*^Q197E^) within the AAA+ ATPase domain of the replication initiator. Together, this study refines annotations within the *S. mutans* essential genome and provides genetic insights into streptococcal chromosome segregation and cell cycle control.

## Introduction

*Streptococcus mutans* is a primary etiological agent of dental caries, a ubiquitous biofilm-mediated disease driven by the pathogen’s ability to metabolize dietary carbohydrates into lactic acid (1, 2). Despite widespread preventative measures, dental caries remains one of the most prevalent chronic diseases globally, placing a massive economic and healthcare burden on populations (3). The pathogen’s residence within the complex, protective matrix of dental plaque renders it highly resilient against environmental fluctuations and traditional antimicrobial approaches. Consequently, identifying the fundamental genetic requirements for *S. mutans* survival is a major priority not only for understanding its core physiology but for discovering novel targets for next-generation therapeutics. Recently, the application of genome-wide screening technologies, such as transposon sequencing (Tn-seq) and CRISPR interference (CRISPRi), has successfully mapped the essential genome of *S. mutans* (4, 5). These high-throughput viability catalogs have identified a core set of approximately 200 genes that are indispensable for bacterial growth and survival.

While these viability catalogs are important, they highlight a secondary bottleneck for the field: translating a list of essential genes into a mechanistic biological understanding. This challenge is compounded by the reliance on the original *S. mutans* UA159 reference genome (6). Sequenced over two decades ago, its functional annotation relies heavily on automated computational pipelines that depend on primary sequence homology (7, 8). These models frequently fail to recognize highly divergent proteins and have not kept pace with recent structural and functional discoveries in related Gram-positive organisms (9). Consequently, approximately 5% of the core essential genome is annotated as hypothetical, or uncharacterized. Defining the functions of these uncharacterized genes is necessary to complete our understanding of basic streptococcal physiology and to evaluate their potential as therapeutic targets.

Characterizing essential genes is inherently difficult because their requirement for viability precludes standard gene disruption in wild-type backgrounds (10). Inducible CRISPR interference (CRISPRi) overcomes this barrier by providing tunable transcriptional repression of essential targets (11–13). This conditional approach enables the phenotypic, ultrastructural, and transcriptomic consequences of target depletion to be evaluated in viable cultures. Furthermore, pairing transcriptional depletion with genetic suppressor selection allows the identification of compensatory mutations and genetic bypass mechanisms that permit viability in the absence of the targeted gene. Together, these complementary strategies provide a framework to assign possible functions to unannotated loci.

In this study, we investigated nine uncharacterized essential genes in *S. mutans* by combining updated bioinformatic analyses, conditional CRISPRi silencing, transmission electron microscopy (TEM), transcriptomics, and genetic suppressor screens. After these experiments we focused on SMU_393, identifying it as a homolog of the pneumococcal chromosome segregation factor RocS whose depletion phenotypes, including cell widening, DNA damage sensitivity, and anucleate cell formation, are suppressed by a point mutation in the replication initiator DnaA. Together, these findings update annotations within the *S. mutans* genome and provide genetic insights into streptococcal chromosome segregation and cell cycle regulation.

## Results and Discussion

### Revising annotations within the core essential genome

The reference genome for *S. mutans* UA159 has served as a foundational cornerstone for oral microbiology for over two decades (6). However, its annotation relies heavily on automated computational pipelines that have not kept pace with recent structural and functional discoveries across microbiology. Consequently, the UA159 genome contains numerous outdated, incomplete, or artifactual functional assignments. This limitation becomes particularly acute when attempting to map the core biological pathways required for pathogen survival. Our recent genome-wide Tn-seq and CRISPRi screens defined a core essential genome of approximately 200 genes required for *S. mutans* viability (4, 5). While most of these genes map to well-characterized features, nine essential genes, representing roughly 5% of the essential genome, are annotated as hypothetical, uncharacterized, or possessing domains of unknown function. To update these annotations, we initiated a systematic re-annotation of these targets. By integrating BLASTp analysis, AlphaFold2 structural predictions, and extensive database mining (summarized in methods), we successfully assigned tentative functions to seven of these genes (Table S1). Comparative analysis revealed that most of these genes are broadly conserved across the *Streptococcus* genus and other Gram-positive bacteria (Figure 1) and we broadly categorized these conserved genes into distinct functional modules:

**Figure 1.**
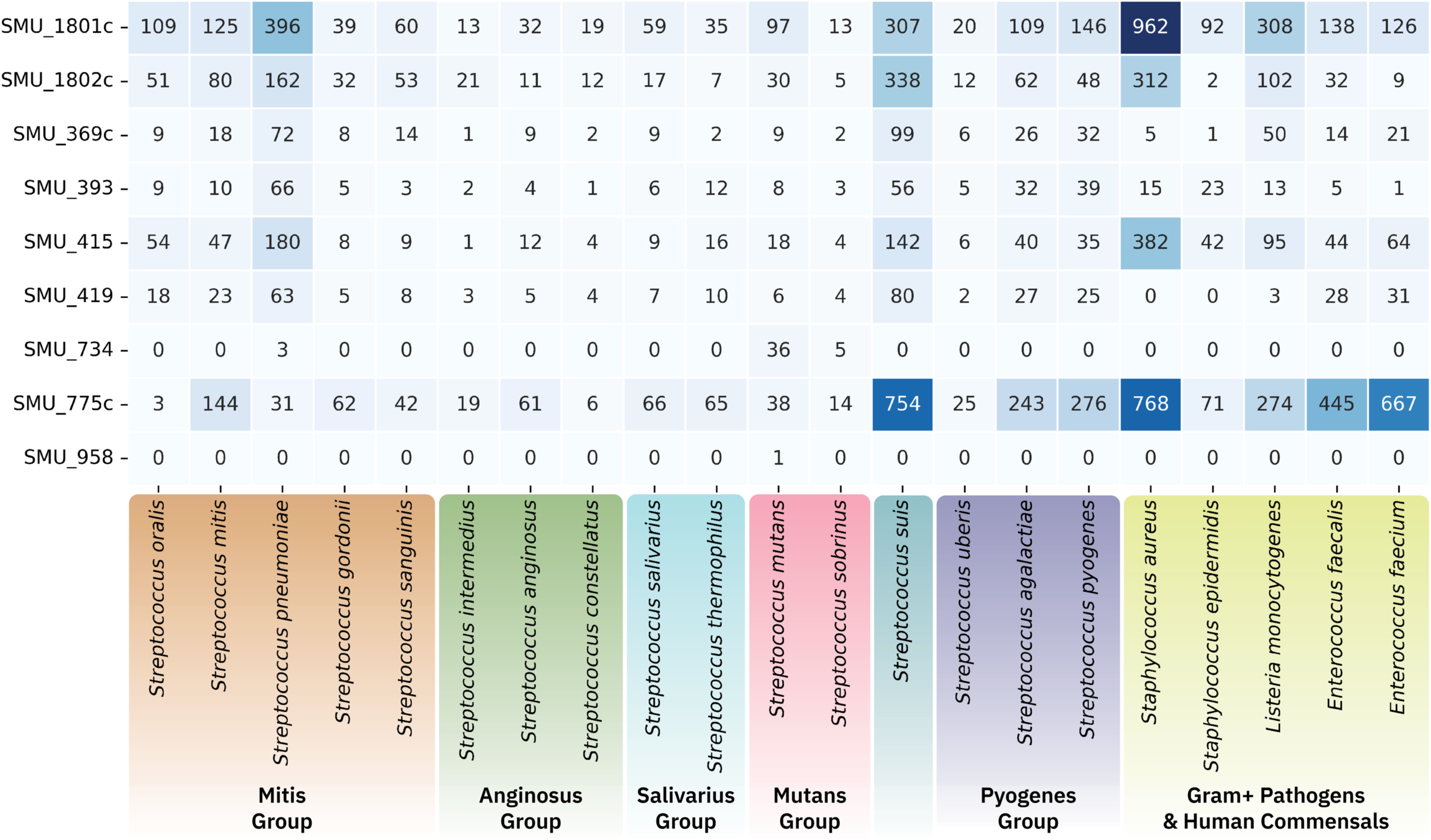
Distribution of candidate essential protein homologs across representative *Streptococcus* species and Firmicutes. Heatmap displaying the total count of retrieved homologs from the NCBI non-redundant (nr) protein database for each *S. mutans* query sequence (rows) across selected bacterial species (columns). Homology searches were performed using BLASTp with filtering criteria of E-value < 10^-5^ and ≥ 90% query coverage. Taxa are grouped by phylogenetic relationship (Mitis, Anginosus, Salivarius, Mutans, Suis, Pyogenes, and other Firmicutes). Color intensity and numerical values indicate the absolute number of matching homolog records in the NCBI database per species, distinguishing broadly conserved loci from those with genus- or species-restricted distributions (with total counts reflecting database genome representation).

#### Cell cycle and chromosome segregation

Two highly conserved genes, SMU_393 and SMU_415, have putative functions in cell cycle coordination. SMU_393 has a putative helix-turn-helix DNA binding domain followed by a coiled coil domain. SMU_393 is a homolog of *S. pneumoniae* RocS (regulator of chromosome segregation), a membrane-bound protein known to interact with DNA, the chromosome partitioning protein ParB, and the capsule regulator CpsD (14–16). Like RocS, SMU_393 possesses a highly conserved C-terminal membrane-targeting region (KKGFFARLFGG in *S. mutans*), a motif originally characterized in the MinD segregation protein of rod-shaped bacteria (17). Furthermore, recent studies highlight that this protein undergoes phosphorylation at T41 (14), a post-translational modification we have also recently observed (18), hinting at dynamic regulatory activity. SMU_415, while containing a predicted choline kinase domain, is a homolog to the essential pneumococcal protein CcrZ (cell cycle regulator interacting with FtsZ). Work in both *S. pneumoniae* and *B. subtilis* indicates that CcrZ coordinates cell division with DNA replication initiation, confers resistance to DNA damage, and phosphorylates an as-yet- unknown substrate rather than choline (19, 20).

#### Transcription, translation, and ribosome biogenesis

A subset of our conserved targets map directly to information processing. SMU_419, possessing a DUF448 domain, is a homolog of the nucleic acid-binding protein YlxR (recently characterized as the RNase P modulator, RnpM) (21, 22). Residing within the essential *nusA*-*infB* operon, SMU_419 (YlxR/RnpM) may function as a conserved RNA-binding protein that modulates RNase P activity, providing a regulatory link between tRNA processing and the core machinery of transcription and translation. SMU_369c is a small (231 bp) gene residing immediately upstream of the RNase J1 gene (*rnjA*). It contains an RpoY (RNA polymerase epsilon subunit) domain motif and has been identified as an RNAP subunit in *B. subtilis*, though its exact physiological necessity remains enigmatic as *B. subtilis rpoY* mutants lack an obvious phenotype (23).

We also identified a highly conserved operon housing SMU_1801c and SMU_1802c. SMU_1801c contains a YqeH ribosome biogenesis GTPase domain; in *B. subtilis*, YqeH is essential for growth, and its depletion causes marked deficits in 70S ribosome assembly (24). Exactly how YqeH exerts its effects on the ribosome is poorly understood, and its physiological role beyond *B. subtilis* has not been studied. Its operon partner, SMU_1802c, is a predicted phosphohydrolase of the haloacid dehalogenase (HAD) IIIA-type family. Recent findings in *B. subtilis*, with a homolog of SMU_1802c, have shown that this gene encodes a phosphatidylglycerophosphate phosphatase (PGP; gene known as *pgpP*) that is responsible for converting PGP into phosphatidylglycerol, a vital lipid component of the bacterial membrane (25). Notably, *pgpP* (also known as *yqeG*) is essential in *B. subtilis*, *E. faecalis*, *L. monocytogenes*, and several streptococci (26).

#### Cell envelope maintenance and niche-specific adaptations

Our analysis mapped SMU_775c to cell envelope integrity. This widely conserved gene contains both lipoteichoic acid (LTA) synthase and phosphoglycerol transferase domains, serving as the likely *S. mutans* equivalent to the essential *S. aureus* type I LTA synthase (LtaS) (27, 28). Conversely, the gene SMU_734 appears highly restricted, found solely within the Mutans group (*S. mutans* and *S. sobrinus*) (Figure 1). Containing a DUF3935 domain, SMU_734 is the least understood of our targets but resides in a putative operon alongside a transcriptional regulator (SMU_730), an ABC transporter (SMU_731), and an inner membrane protein (SMU_732).

Finally, our bioinformatic analysis highlighted issues with the legacy genome annotations by resolving SMU_958, a locus restricted to *S. mutans* UA159, as an annotation artifact. The predicted open reading frame directly overlaps with the essential L10 (RplJ) and L12 (RplL) ribosomal proteins, confirming its essentiality is tied entirely to canonical translation machinery rather than a novel hypothetical function.

### Phenotypic profiling of essential gene knockdowns under stress conditions

To map the physiological roles of these genes, we deployed our established inducible CRISPRi system (4) to generate targeted knockdowns that circumvent the immediate lethality of complete gene deletion. Unfortunately, despite several attempts, we were never successful in obtaining sgRNAs targeting SMU_734 and SMU_775c but we were successful for all other genes. To resolve the specific cellular pathways these genes support, we subjected the knockdown strains to an extensive agar plate stress test. By spot-plating serial dilutions of each strain onto agar supplemented with 0.5% xylose (to induce sgRNA expression) alongside an array of targeted physiological stressors, we determined their distinct growth defects (Figure 2). This robust phenotypic profiling allowed us to stratify these uncharacterized genes into distinct functional vulnerabilities.

**Figure 2.**
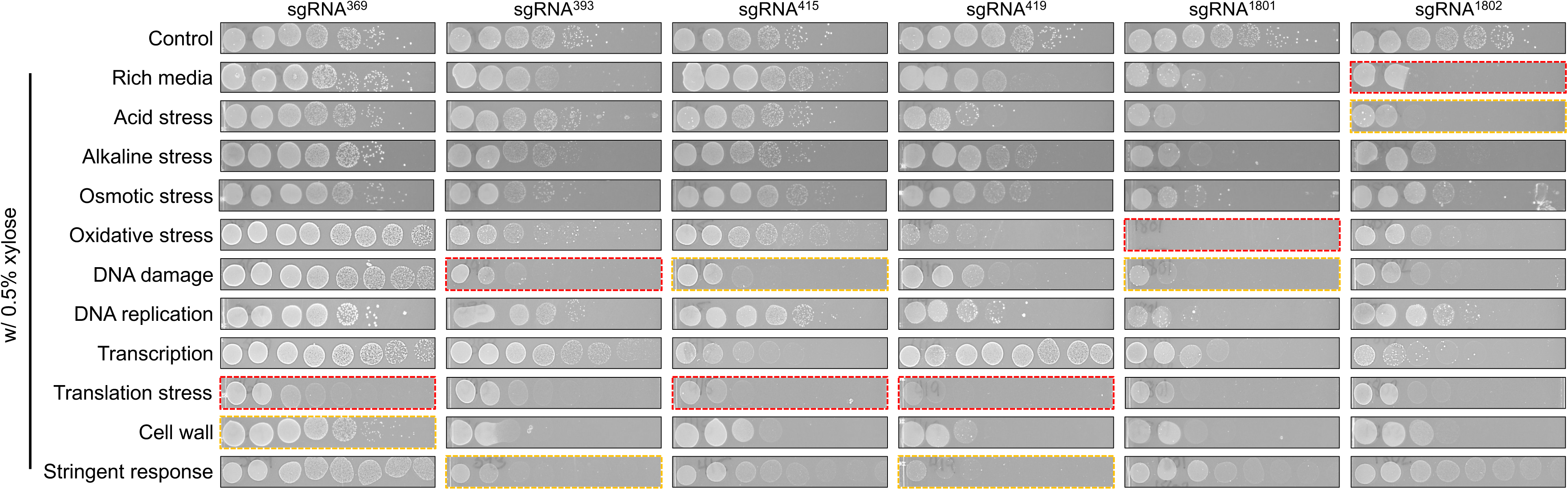
Stress phenotyping reveals distinct cellular vulnerabilities upon essential gene depletion. Viability and stress tolerance of *S. mutans* CRISPRi knockdown strains were evaluated via agar spotting assays. Ten-fold serial dilutions of the indicated strains were spotted onto solid rich medium in the absence (control) or presence (0.5% xylose) of the dCas9 inducer. Plates were supplemented with the following stressors: acid (pH 5.5), alkaline (pH 8.5), osmotic (2.5% NaCl), oxidative (1 mM H_2_O_2_), DNA damage (10 ng/mL mitomycin C), DNA replication (1.5 µg/mL ciprofloxacin), transcription (2 µg/mL rifampicin), translation (2 µg/mL chloramphenicol), cell wall (2 µg/mL ampicillin), and the stringent response (125 ng/mL mupirocin). Red and orange boxes show conditions where strains showed the highest and second highest sensitivity to those conditions respectively.

Strains targeting SMU_369c, SMU_415, and SMU_419 resulted in reduced growth on plates supplemented with the translation inhibitor chloramphenicol (Figure 2). For SMU_419, sensitivity to translation inhibition is consistent with predicted roles in translation/RNA processing. For SMU_415 (a predicted CcrZ homolog), reduced tolerance to translation stress was unexpected given its predicted role in cell division and replication coordination. However, transcriptomic analysis revealed that CRISPRi targeting of SMU_415 causes polar downregulation of the downstream gene *trmB* (SMU_416), which encodes a tRNA (guanine-N(7)-)-methyltransferase (Table S5), suggesting that polar effects on tRNA modification may contribute to this phenotype. Depletion of SMU_369c caused only mild growth reduction under translation stress (Figure 2), indicating that its initial classification as essential in Tn-seq screens likely reflects polar effects on downstream *rnjA* rather than an absolute requirement for SMU_369c itself.

Repression of SMU_393 and SMU_415 resulted in increased sensitivity to the DNA cross-linking agent mitomycin C (MMC). In *S. pneumoniae*, RocS is involved in chromosome segregation and nucleoid protection against DNA damage, consistent with the MMC sensitivity observed upon SMU_393 depletion. Similarly, the DNA damage sensitivity observed during SMU_415 knockdown mirrors phenotypes reported for *ccrZ* mutants in *B. subtilis* (20).

Both the SMU_393 and SMU_419 knockdown strains also displayed reduced viability when challenged with mupirocin, an inhibitor of isoleucyl-tRNA synthetase used to induce the stringent response (29, 30). In other Firmicutes, the stringent response coordinates replication arrest with translation shutdown. The increased sensitivity of SMU_393 and SMU_419 knockdowns to mupirocin is consistent with proposed roles for RocS and RnpM/YlxR in maintaining chromosome organization and modulating RNA processing during metabolic stress. In *B. subtilis*, RnpM functions as a high-affinity modulator of RNase P (21); whether SMU_419 similarly coordinates tRNA maturation with metabolic cues during stringent response induction in *S. mutans* warrants direct biochemical investigation.

Among the tested targets, knockdown of SMU_1802c (*pgpP*) produced the most pronounced growth defect on standard BHI agar upon xylose induction, showing a multi-log reduction in plating efficiency even in the absence of exogenous stressors (Figure 2). In addition, the SMU_1802c knockdown exhibited heightened sensitivity to low pH (pH 5.5) and cell wall-targeting antibiotics (ampicillin). Because PgpP catalyzes the dephosphorylation of phosphatidylglycerol phosphate to phosphatidylglycerol, the major anionic phospholipid in the Firmicute cell membrane (25), depletion of this enzyme possibly disrupts membrane lipid composition and integrity, rendering cells more vulnerable to envelope and acid stress.

To examine whether gene depletion altered cell envelope or division morphology, we imaged CRISPRi knockdown strains using transmission electron microscopy (TEM) and measured cell length and width distributions (Figure 3). Because *S. mutans* is an ovococcus that relies on the precise, spatiotemporal coordination of peripheral elongation and mid-cell septation, disruptions to essential pathways can manifest as morphological aberrations (31). Wild-type *S. mutans* exhibited characteristic ovoid morphology (Figure 3A). Depletion of SMU_369c, SMU_415, SMU_419, or SMU_1801c did not result in obvious alterations in cell shape or dimensions (Figure 3A). In contrast, depletion of SMU_393 led to a noticeable increase in cell width across a small subset of imaged cells (Figure 3A and 3B). In *S. pneumoniae*, loss of RocS disrupts the coordination between chromosome segregation and division site selection, yielding division abnormalities and anucleate cells (15). The morphological heterogeneity observed during SMU_393 depletion is consistent with a similar uncoupling of cell cycle coordination in *S. mutans*.

**Figure 3.**
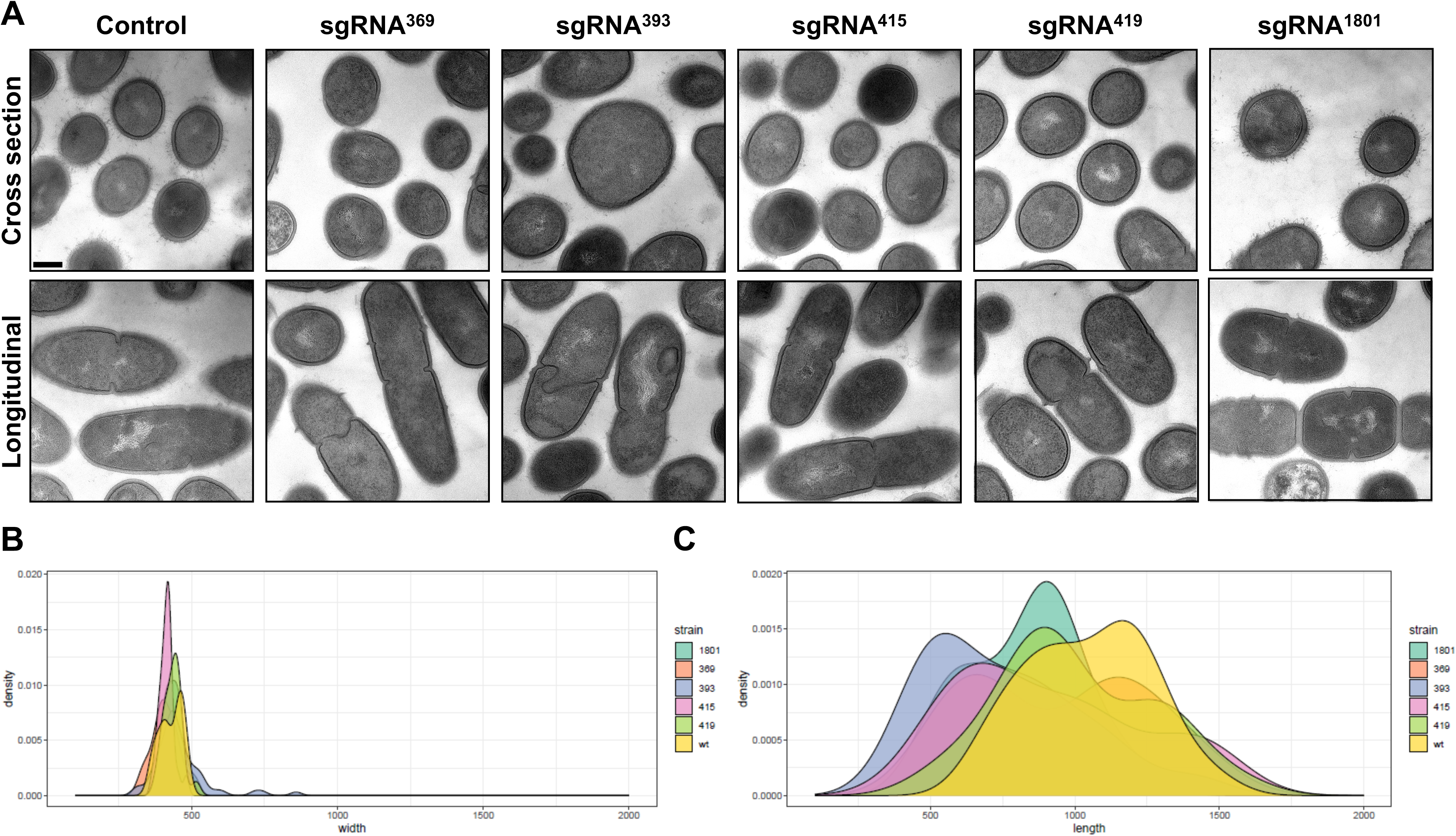
Transmission electron microscopy of *S. mutans* following CRISPRi-mediated knockdown of essential genes. (A) Transmission electron microscopy (TEM) was performed on stationary-phase cells to resolve the structural impact of essential gene depletion. Control cells (UA159) grown in the presence of 0.5% xylose exhibit characteristic ovoid streptococcal morphology with well-defined, symmetrical septa. Black scale bar is 200 nm. Image analysis, using ImageJ, was implemented to measure cell width (B) and cell length (C) from TEM micrographs. The median number of cells measured was 24.

### Genetic suppressor screens identify bypass mechanisms for putative essential genes

While CRISPRi allows phenotypic evaluation of gene depletion, targeted gene deletion provides a direct test of whether a gene is indispensable. We attempted to replace each of the nine candidate essential genes with a non-polar kanamycin resistance cassette (*aphA3*) via homologous recombination in both wild-type UA159 and a high-density transposon-mutagenized library (Figure 4A). Five genes, SMU_415, SMU_419, SMU_775c, SMU_1801c, and SMU_1802c, were refractory to deletion in both genetic backgrounds, indicating an absolute requirement for viability under the conditions tested (Figure 4A).

**Figure 4.**
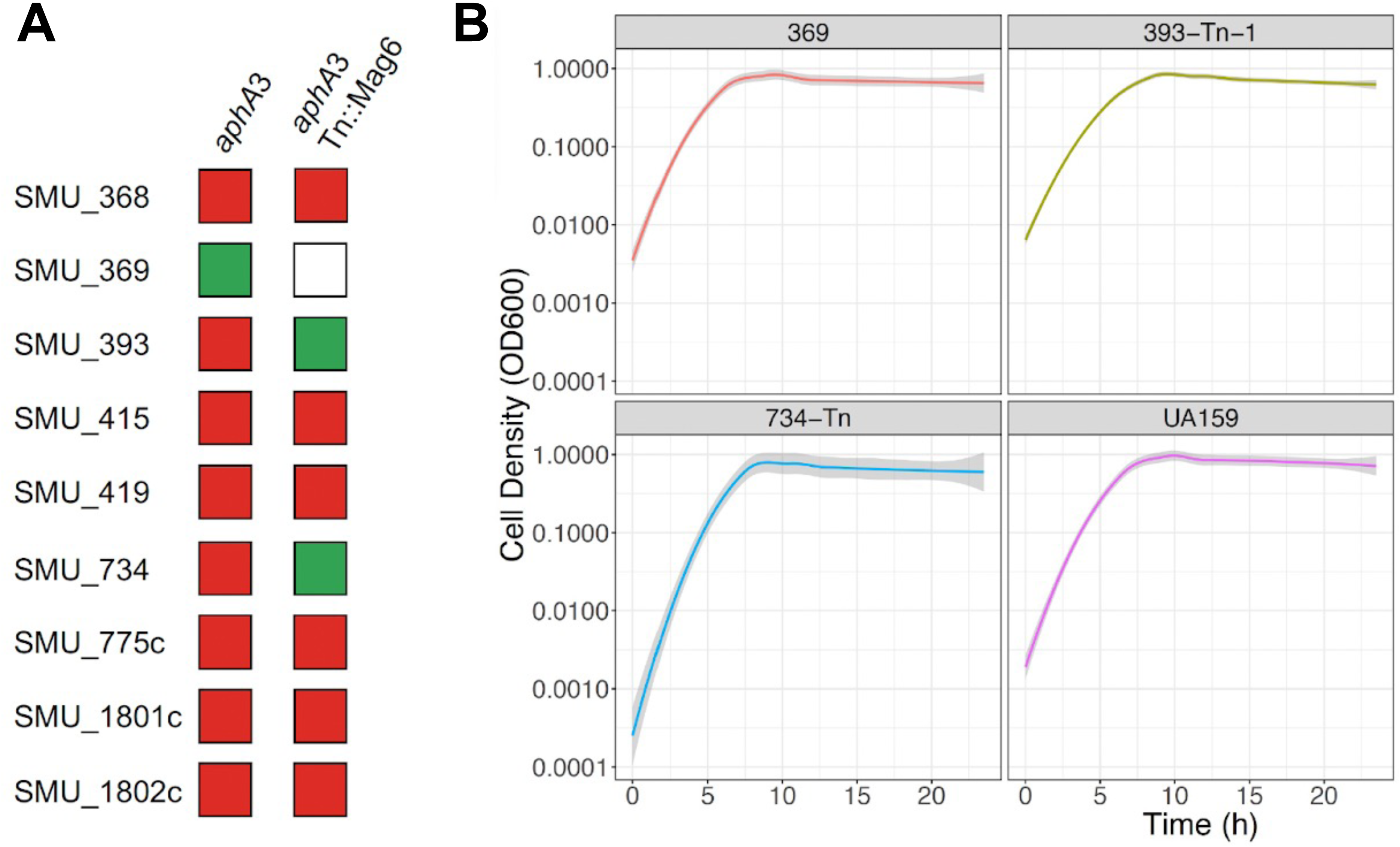
Genetic suppressor screens and growth kinetics identifying bypass mutations in *S. mutans*. (A) A matrix mapping the recovery of viable deletion mutants across nine target loci using a non-polar kanamycin resistance cassette (*aphA3*). Green squares indicate successful recovery of viable deletion mutants, red squares indicate that the deletion was intractable (lethal), and white squares indicate conditions not evaluated. Direct mutagenesis was performed in either a clean wild-type background (*aphA3*) or inside a high-density transposon insertion library (*aphA3* Tn::Mag6) to uncover genetic escape pathways and context-dependent suppressors. (B) Growth curves tracking cell density (OD_600_ on a logarithmic scale) over 24 hours in rich media. Shaded ribbons along each curve denote the variance across three independent biological replicates.

In contrast, deletions were successfully obtained for three loci under specific genomic conditions. Once viable deletion strains were established, both the ΔSMU_393 and ΔSMU_734 strains exhibited growth kinetics in rich medium comparable to wild-type UA159 (Figure 4B). Deletion of SMU_393 could not be recovered in wild-type UA159 but was obtained from the transposon library background. Whole-genome sequencing of this initial isolate identified a transposon insertion in SMU_2070 (427 bp into the coding sequence) and a single-nucleotide polymorphism in *dnaA* resulting in a glutamine-to-glutamate substitution at position 197 (Q197E). SMU_2070 contains a YaaA-like peroxide stress domain (E-value of 2.2 x 10^-86^) implicated in oxidative stress and DNA repair in *Escherichia coli* (32). Position Q197 of DnaA resides within the surface-exposed AAA+ ATPase domain (Domain III; residues 111–329) (Figure S1), which participates in oligomerization and ATP hydrolysis (33, 34). As detailed below, subsequent reconstruction of the *dnaA* Q197E point mutation in clean backgrounds demonstrated that this mutation suppresses defects caused by SMU_393 depletion.

Viable ΔSMU_734 mutants were also recovered from the transposon library background. Sequencing of the ΔSMU_734 isolate revealed a transposon insertion within the SMU_136c-*mleS* intergenic region and a spontaneous ∼75 kb genomic deletion spanning from *bacD* to SMU_1408c, corresponding to the loss of the Tn*Smu2* genomic island and an adjacent ∼20 kb region (Figure S2). Tn*Smu2* encodes a nonribosomal peptide synthetase/polyketide synthase (NRPS-PKS) cluster involved in pigment production and oxidative tolerance (35). Tn*Smu2* is known to undergo spontaneous excision during envelope or metabolic stress, such as in *cidB* holin mutants (36). The loss of Tn*Smu2* in the ΔSMU_734 isolate suggests that SMU_734 may function in managing metabolic demands associated with this island, though the precise mechanism of suppression remains to be defined. Although targeted reconstruction of this large-scale deletion will be required to formally verify genetic causality, these genomic alterations identify a putative bypass route and provide an initial framework for future dissection of SMU_734 function.

Finally, we obtained a clean deletion of SMU_369c in the wild-type UA159 background without secondary mutations, demonstrating that SMU_369c is dispensable for viability. Its prior classification as essential in Tn-seq screens likely reflects transcriptional polarity on the downstream essential ribonuclease J1 gene (*rnjA*/SMU_368c). In support of this, we were unable to delete *rnjA* in either wild-type or library backgrounds (Figure 4A), and CRISPRi-mediated knockdown of *rnjA* caused severe growth defects across all tested conditions (Figure S3). While Chen *et al*. (37) previously reported a viable Δ*rnjA* strain in S*. mutans*, that mutant displayed severe growth defects, consistent with the strong phenotypic consequences of *rnjA* repression observed here.

### Defining the transcriptomic signature of essential gene depletion

To determine how *S. mutans* globally responds to the failure of varied essential pathways, we performed RNA-seq profiling across our CRISPRi knockdown strains (targeting SMU_368c, 369c, 415, 419, 1801c, 1802c, and 393; see Table S2-S8). To further investigate the potential polar effects of SMU_369c knockdown on the downstream essential gene *rnjA* (SMU_368c), we included a direct CRISPRi-targeted depletion of SMU_368c in our transcriptomic analysis. As an internal control for CRISPRi induction, *cas9* (SMU_1405c) was universally upregulated across all xylose-induced samples (log_2_FC = +6.37 to +8.00). Targeted repression of the intended transcripts was confirmed in the following strains: SMU_368c (log_2_FC = -4.77), SMU_393 (log_2_FC = -2.94), SMU_415 (log_2_FC = -2.43), SMU_419 (log_2_FC = -1.88),

SMU_1801c (log_2_FC = -3.68), and SMU_1802c (log_2_FC = -4.12). Using a significance cutoff of FDR < 0.05 and log2FC > 1, the total number of differentially expressed genes (DEGs) varied considerably among knockdowns. Depletion of the essential ribonuclease J1 (SMU_368c) caused the largest transcriptional disruption (604 DEGs), followed by SMU_1801c (295 DEGs) and SMU_419 (221 DEGs), whereas depletion of SMU_393 resulted in a more targeted profile (64 DEGs).

Hierarchical clustering of the knockdown strains based on genes with marked differential expression (log_2_FC > 2; Figure 5) revealed shared transcriptional patterns across multiple conditions. Depletion of SMU_368c, SMU_393, SMU_415, SMU_419, SMU_1801c, and SMU_1802c consistently resulted in reduced transcript abundance for the *citZ*-*citB*-*idh* locus (citrate synthase, aconitase, and isocitrate dehydrogenase). To confirm that this downregulation was not an off-target artifact of dCas9 expression or xylose exposure, we evaluated two independent controls. First, in our previous RNA-seq profiling of the *S. mutans* CRISPRi platform (4), xylose induction of dCas9 paired with a non-targeting sgRNA (targeting *gfp*) had no detectable effect on the locus: *citZ* (log_2_FC = -0.08, FDR = 0.94), *citB* (log_2_FC = +0.07, FDR = 0.91), and *idh* (log_2_FC = +0.001, FDR > 0.99). Second, within the present study, the SMU_369c knockdown strain serves as an internal biological control; despite robust d*cas9* induction (log_2_FC = +6.37; Table S3), the non-essential status of SMU_369c and its lack of growth impairment left the *citZ*-*citB*-*idh* locus unaffected. While growth rate deceleration likely contributes to reduced expression of core TCA cycle genes, this shared downregulation reflects a reproducible physiological state associated with essential gene impairment in *S. mutans*.

**Figure 5.**
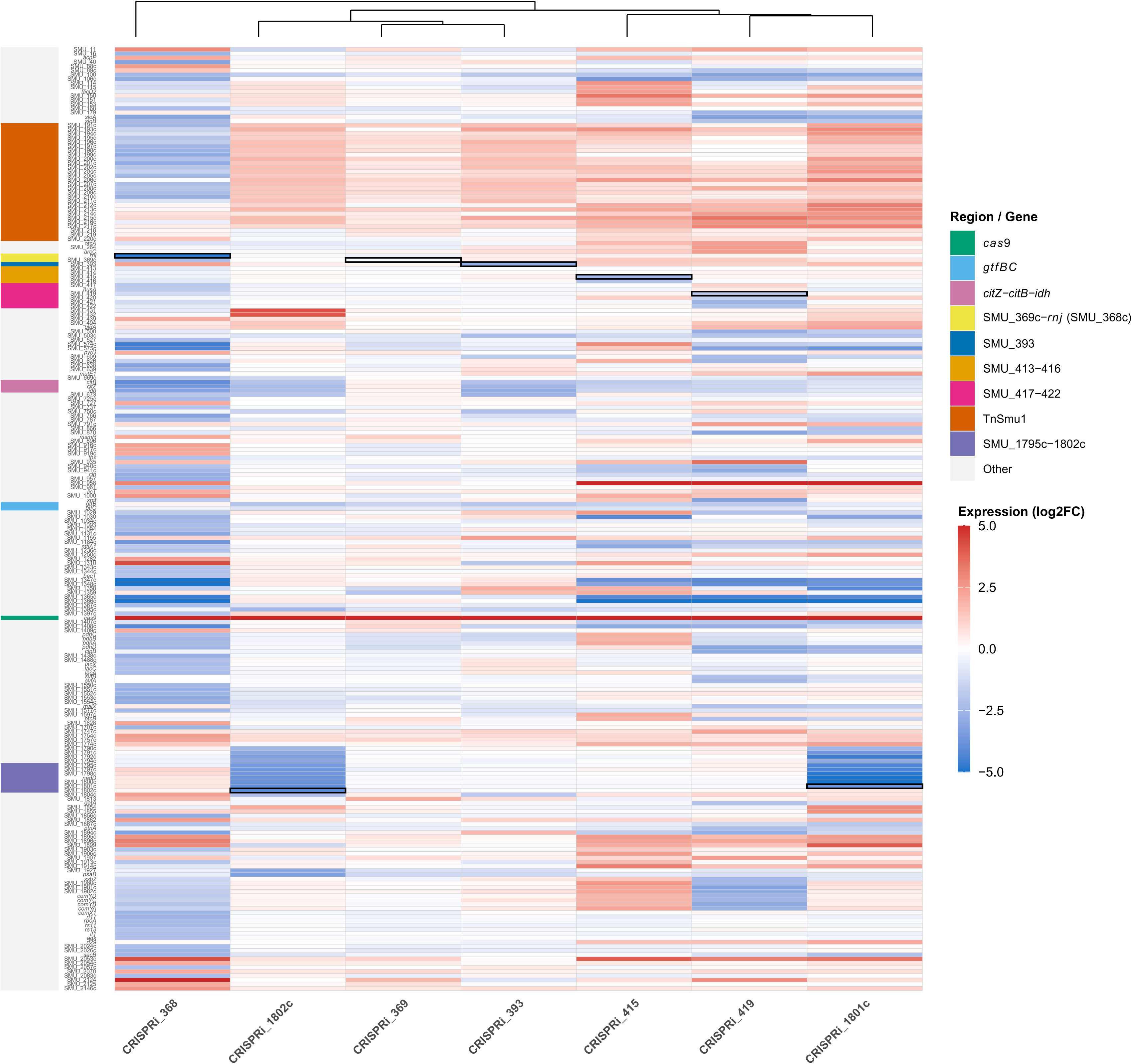
Comparative transcriptomic profiling of uncharacterized essential gene knockdowns in *S. mutans*. RNA-seq was performed on seven CRISPRi-targeted strains following xylose-induced depletion of uncharacterized essential genes. The color gradient indicates transcriptional upregulation (red) or downregulation (blue) on a scale from -5.0 to +5.0. Color-coded sidebars on the left denote specific genes, operons, and genomic regions of interest, including *cas9* (SMU_1405c), *gtfBC*, the *citZ*-*citB*-*idh* metabolic locus, the Tn*Smu1* mobile genetic element, and targeted operonic regions (SMU_369c-*rnj*, SMU_393, SMU_413–416, SMU_417–422, and SMU_1795c–1802c). Target gene expression, in the respective CRISPRi strain, is marked with a thick line box. Strains were clustered using Euclidean distance to identify strains sharing co- regulatory signatures.

In addition to *citZ*-*citB*-*idh* downregulation, several knockdowns exhibited increased transcript levels across the Tn*Smu1* genomic island (SMU_191c to SMU_220). In five knockdown strains (SMU_393, SMU_415, SMU_419, SMU_1801c, and SMU_1802c), up to 24 of the 28 annotated genes in this region were upregulated (log_2_FC = +1.5 to +3.5). To rule out non-specific stress responses driven by the silencing machinery itself, we examined island expression under control conditions. Consistent with the metabolic locus, analysis of our prior non-targeting CRISPRi dataset (4) confirmed that dCas9 induction alone does not activate the island (FDR > 0.17 across all 28 genes), indicating that Tn*Smu1* upregulation is a response to the stress of essential gene failure rather than a platform artifact. TnS*mu1* is an integrative and conjugative element (ICE) known to be induced by DNA damage and biofilm growth conditions (38, 39). In contrast, targeted knockdown of RNase J1 (SMU_368c) led to significant downregulation of 15 genes within Tn*Smu1*. Because RNase J1 is a primary enzyme involved in global RNA processing and decay (37), its depletion likely alters transcript stability across the mobilome differently than general growth stress.

Pathway-specific expression differences were also observed among the knockdowns, particularly regarding glucosyltransferases involved in extracellular matrix production. While depletion of translation-associated factors (SMU_368c and SMU_1801c) did not significantly reduce *gtfBC* expression, knockdown of genes potentially involved in cell cycle and DNA replication (SMU_393 and SMU_415), as well as SMU_419, resulted in downregulation of *gtfB* (log_2_FC = -1.55 to -2.04).

The RNA-seq datasets also provided direct confirmation of CRISPRi-mediated transcriptional polarity within operons. Because dCas9 blocks transcriptional elongation, repression of upstream genes frequently reduces expression of downstream operon members. For example, knockdown of SMU_1802c caused strong polar downregulation of downstream genes within the same transcriptional unit, including *nadD* (SMU_1799; log_2_FC = -3.77), an essential nicotinate mononucleotide adenylyltransferase. A similar polar effect on *nadD* was detected upon knockdown of the upstream partner SMU_1801c. Despite these shared polar effects on adjacent genes, global DEG profiling demonstrated that SMU_1801c and SMU_1802c knockdowns cluster into distinct clades (Figure 5). For instance, SMU_1802c depletion uniquely induced expression of SMU_431 and SMU_432 (log_2_FC > +4.0), which encode an ABC transporter coupled to the SMU_433/434 LytTR Regulatory System (LRS) (40). These data indicate that while polar effects occur within operonic loci, target-specific transcriptional signatures can still be distinguished.

### Characterization of SMU_393 as a functional homolog of the chromosome segregation factor RocS

Among the evaluated targets, SMU_393 displayed phenotypic and genetic characteristics consistent with cell cycle and chromosome organization functions. Although annotated as a hypothetical protein in the UA159 genome (6), structural modeling using AlphaFold2 revealed high similarity to *Streptococcus pneumoniae* RocS (TM-score = 0.86; Figure S4) (15, 16). SMU_393 also retains the conserved C-terminal membrane-targeting sequence (MTS; KKGFFARLFGG) required for membrane association in pneumococcal RocS (Figure S4) (16).

To further understand the global cellular response to SMU_393 depletion, we performed quantitative proteomic (MS/MS) profiling (Table S9). MS/MS analysis confirmed the CRISPRi-mediated depletion of the SMU_393 protein (log_2_FC = -3.98; FDR < 0.001). Proteomic analysis revealed a significant downregulation of the error-prone DNA polymerase IV, DinB (log_2_FC = -1.31; FDR < 0.001). In *E. coli*, DinB participates in translesion DNA synthesis to bypass replication-blocking lesions (41). In addition, several cell envelope-associated proteins showed increased abundance during SMU_393 depletion, including the water-soluble glucan synthesizer GtfD (log_2_FC = +1.98; P < 0.001), the glucan-binding protein GbpD (log_2_FC = +1.50; P < 0.001), the surface protein WapE (log_2_FC = +1.37; P < 0.001), and a putative 5′-nucleotidase (SMU_1213c; log_2_FC = +1.64; P < 0.001). Conversely, insoluble glucan synthases GtfB and GtfC remained stable or slightly increased at the protein level (log_2_FC = +0.42 and +0.61, respectively; P < 0.01), despite the modest reduction in *gtfB* transcript detected by RNA-seq. Similar to the transcriptomic findings in slow-growing strains, the TCA cycle enzymes aconitate hydratase (AcnA; log_2_FC = -1.26; P < 0.001) and isocitrate dehydrogenase (Icd; log_2_FC = -1.24; P < 0.001) were depleted at the protein level.

To determine whether the *dnaA* Q197E mutation directly bypasses SMU_393 depletion independently of the secondary SMU_2070 transposon insertion, we introduced the *dnaA* Q197E point mutation into clean wild-type and CRISPRi backgrounds using markerless site-directed mutagenesis. We then evaluated chromosome segregation and DNA damage sensitivity. Fluorescence microscopy of DAPI-stained cells showed that CRISPRi depletion of SMU_393 resulted in an accumulation of anucleate cells (∼11.4% of total cells), indicative of a chromosome segregation defect (Figure 6A and B). Introduction of the *dnaA* Q197E allele into the SMU_393 knockdown background restored chromosome inheritance, reducing the proportion of anucleate cells to wild-type baseline levels (Figure 6A and B).

**Figure 6.**
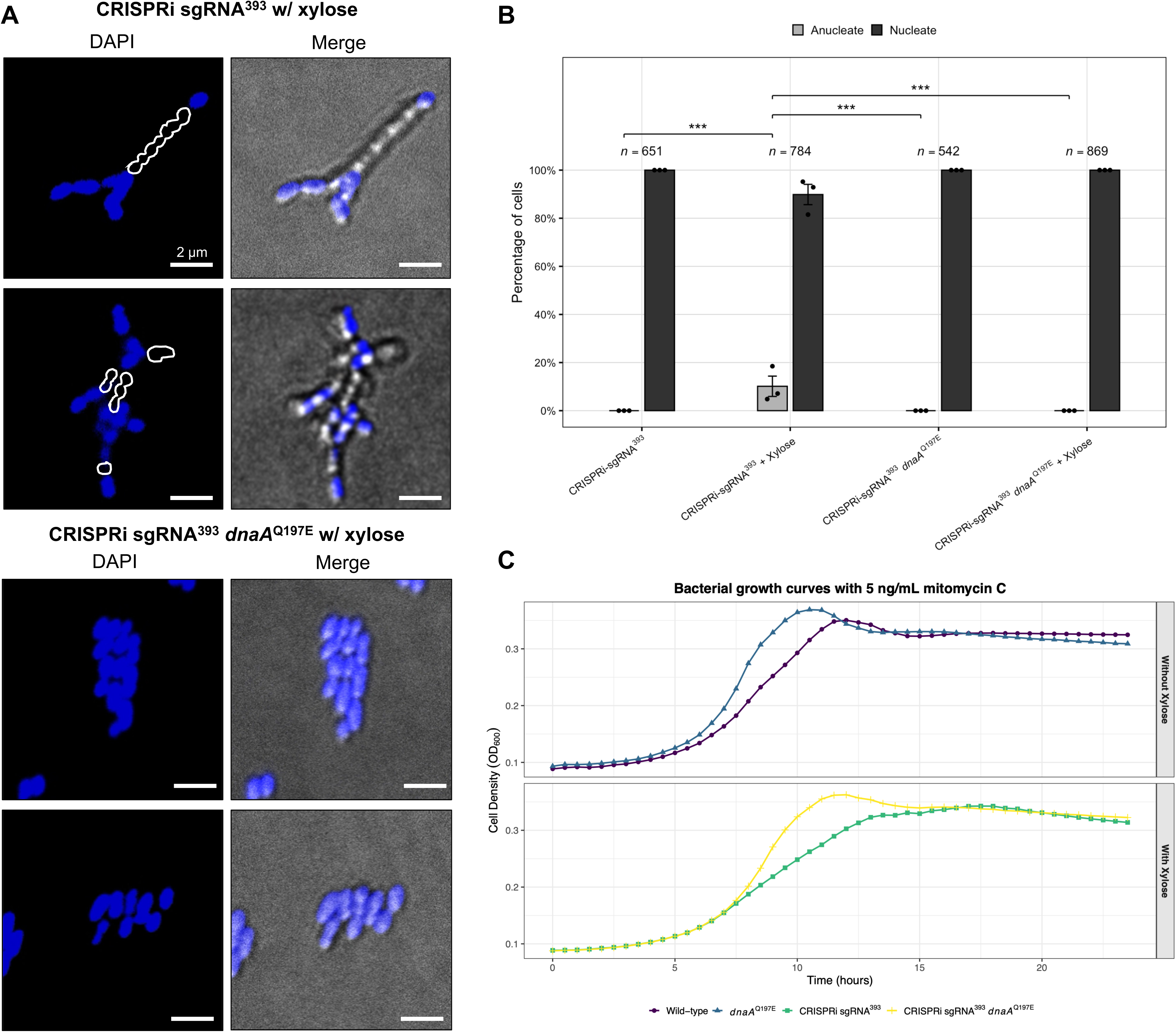
The *dnaA*^Q197E^ mutation suppresses chromosome segregation defects and mitomycin C (MMC) hypersensitivity induced by SMU_393 depletion. (A) Exponential-phase *S. mutans* cells stained with DAPI to visualize chromosome distribution. CRISPRi-mediated knockdown of SMU_393 yields a distinct subpopulation of anucleate cells that fail to inherit chromosomal DNA (white border around cells lacking a chromosome). This physical segregation defect is absent in the same background when the suppressive *dnaA* Q197E mutation is present. White scale bar is 2 µm. (B) Image analysis counting the percentage of anucleate cells across CRISPRi SMU_393 mutant backgrounds. Data are representative of counted fields from independent biological replicates. (C) Growth curves monitoring cell density over 24 hours in the presence of 5 ng/mL mitomycin C (MMC).

Furthermore, expressing *dnaA* Q197E in the wild-type background conferred increased baseline tolerance to MMC, and its introduction into the CRISPRi SMU_393 background mitigated the growth lag and arrest caused by MMC challenge during SMU_393 repression (Figure 6C).

Together, the structural homology, phenotypic consequences of depletion (MMC sensitivity, cell widening, and anucleate cell formation), proteomic shifts, and genetic suppression by *dnaA* Q197E support the classification of SMU_393 as a functional homolog of RocS in S. mutans (Figure 7). In *S. pneumoniae*, RocS links the chromosome partitioning machinery to the membrane to coordinate nucleoid segregation with division (14, 15). Our data indicate that SMU_393 performs a similar coordinating role in *S. mutans*, where loss of this factor uncouples DNA inheritance from division and alters cell envelope homeostasis. The ability of a point mutation in the AAA+ domain of DnaA to suppress these partitioning and stress-sensitivity phenotypes suggests that altering replication initiation dynamics can compensate for the absence of RocS-mediated coordination.

**Figure 7.**
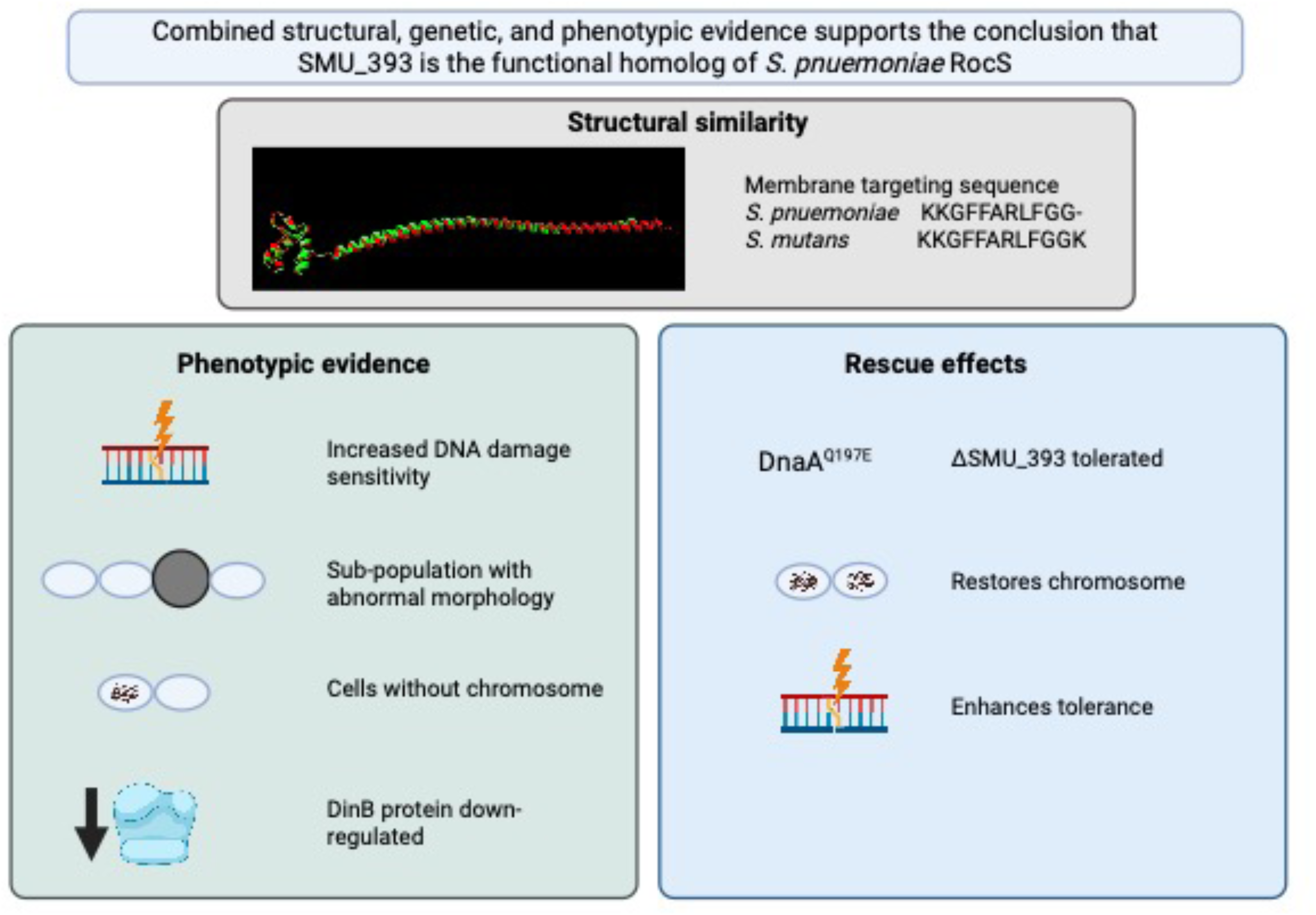
Structural, phenotypic, and genetic evidence establishing SMU_393 as the *Streptococcus mutans* RocS functional equivalent. High-resolution structural overlay modeling of *S. mutans* SMU_393 and *S. pneumoniae* RocS. The adjacent sequence alignment highlights the strict conservation of the C-terminal amphipathic alpha-helix membrane-targeting sequence (MTS) between the two species, which is required for membrane association. Physiological and molecular defects caused by depletion of SMU_393 include significantly increased sensitivity to DNA damage (mitomycin C), a sub-population exhibiting abnormal morphology (increased cell width), the accumulation of anucleate cells lacking chromosomal DNA, and the proteomic downregulation of the error-prone lesion-bypass DNA polymerase IV (DinB). Importantly, some of these phenotypes can be suppressed by a mutation in the DnaA replication protein. The *dnaA*^Q197E^ mutation renders the otherwise lethal ΔSMU_393 deletion tolerated. The suppressor completely restores regular chromosome inheritance to wild-type baselines and enhances baseline tolerance against DNA cross-linking agents.

## Conclusions

In this study, we combined bioinformatic analyses, conditional CRISPRi silencing, comparative transcriptomics and proteomics, and genetic suppressor selection to re-evaluate hypothetical essential genes in *Streptococcus mutans*. Our findings assign putative biological functions to seven previously uncharacterized genes involved in translation, membrane biogenesis, and cell division, while identifying annotation artifacts and operonic polar effects that confound essentiality screens. Furthermore, we established SMU_393 as a functional homolog of the pneumococcal chromosome segregation factor RocS and demonstrated that its depletion phenotypes can be suppressed by a point mutation within the replication initiator DnaA. Together, these findings update annotations within the *S. mutans* genome and provide genetic insights into streptococcal chromosome partitioning and cell cycle coordination.

## Materials and Methods

### Bacterial strains and culture conditions

Strains of *Streptococcus mutans* were cultured from single colonies in Brain Heart Infusion (BHI) broth (Difco). Unless otherwise noted, *S. mutans* was routinely cultured at 37°C in a 5% CO_2_ microaerophilic atmosphere. *Escherichia coli* strains were routinely cultured in LB broth (Lennox formula; 10 g/L tryptone, 5 g/L yeast extract and 5 g/L NaCl) at 37°C with aeration. Antibiotics were added to growth media at the following concentrations: erythromycin (5 µg/mL for *S. mutans*, 300 µg/mL for *E. coli*), kanamycin (1.0 mg/mL for *S. mutans*, 50 μg/mL for *E. coli*), and spectinomycin (1.0 mg/mL for *S. mutans*). CRISPRi experiments were conducted in a chemically defined medium (CDM) as described by Chang et al. (42) supplemented with 0.5% maltose. A list of strains and plasmids (Table S10) can be found in the supplementary material.

### Homolog analysis

Using the FASTA sequences of each uncharacterized protein, a BLASTP was run against the non-redundant (nr) protein database of the National Center for Biotechnology Information (NCBI) (43). The search was performed using the command line application provided by NCBI, with search limited to the taxa of interest. Results were filtered to include those with an E-value < 10^-5^ and with at least a 90% coverage of the uncharacterized query sequences. The number of results was counted for each target protein and taxa to identify the potential number of homologs as given by NCBI records, as shown in Figure 1.

To update legacy annotations for essential hypothetical proteins in *S. mutans* UA159 identified through Tn-seq, we implemented a multi-layer comparative annotation workflow integrating sequence homology, protein domain analysis, orthology inference, genomic context, structural prediction, curated biological databases, and manual literature review. Protein sequences were retrieved from the GenBank *S. mutans* UA159 reference genome (GenBank accession AE014133.2) (6) and cross-referenced against multiple annotation resources, including RefSeq (44), UniProtKB (45), KEGG (46), BioCyc (47), and BV-BRC (48). Conserved protein domains were identified using InterPro (49) and the NCBI Conserved Domain Database (CDD) (50), while orthology relationships were evaluated using KEGG Orthology (KO) (51), PANTHER (52), and eggNOG version 7 (53). Gene neighborhood, operon organization, phylogenetic distribution, and predicted functional associations were examined using STRING (54), SeqHub (55), and the Enzyme Function Initiative Genome Neighborhood Tool (EFI-GNT) (56, 57) to strengthen functional assignments and distinguish functionally divergent paralogous protein families. Candidate homologs were prioritized based on reciprocal sequence similarity, alignment coverage, conservation of catalytic and other functionally important residues, taxonomic distribution, and preservation of genomic context. Functional assignments were evaluated using complementary evidence from UniProtKB, BioCyc, KEGG, and BV-BRC, and experimentally validated functions were confirmed through manual review of the primary literature identified using PubMed and PaperBLAST (58). Where appropriate, AlphaFold (59) structural models and experimentally determined protein structures were compared using UCSF ChimeraX version 1.10.1 (60) to evaluate conservation of overall protein architecture, active-site geometry, and characteristic functional motifs. Structural evidence was used as supporting evidence for functional assignments and was interpreted alongside sequence conservation, genomic context, and experimentally characterized orthologs rather than as a standalone predictor of protein function.

Gene Ontology (GO) molecular function and biological process terms were assigned using experimentally validated annotations from the corresponding orthologs. Protein names, functional descriptions, and GO annotations were manually reviewed for consistency across databases and reconciled with the primary literature. PubMed identifiers supporting each functional assignment were recorded. The final annotations reported in Supplementary Table S1 include the *S. mutans* locus tag, UniProt accession, original GenBank annotation, experimentally validated homolog, homolog accession number, source organism, sequence identity and alignment coverage, supporting PubMed reference, proposed biological role, and associated GO molecular function and biological process terms.

### Phenotypic analysis

To resolve the physiological roles of the targeted essential genes, CRISPRi knockdown strains were subjected to phenotyping via agar plate spot-dilution assays. Strains were initially cultured overnight to stationary phase in BHI broth. Following growth, 10-fold serial dilutions of each culture were prepared in PBS and spotted (5 µL) onto solid rich medium (BHI agar). To induce dCas9-mediated transcriptional repression, the agar was supplemented with 0.5% (w/v) xylose. To identify specific functional vulnerabilities, the medium was further supplemented with an array of targeted physiological stressors: acid (pH 5.5), alkaline (pH 8.5), osmotic (2.5% NaCl), oxidative (1 mM H_2_O_2_), DNA damage (10 ng/mL mitomycin C), DNA replication (1.5 µg/mL ciprofloxacin), transcription (2 µg/mL rifampicin), translation (2 µg/mL chloramphenicol), cell wall (2 µg/mL ampicillin), and the stringent response (125 ng/mL mupirocin). Plates were incubated at 37°C in a 5% CO_2_ microaerophilic atmosphere for 24 to 48 hours. Growth defects were evaluated by comparing xylose-induced spots against non-induced controls.

### Transmission electron microscopy

Bacterial cultures were grown overnight to stationary phase with appropriate concentrations of xylose (w/v) in BHI broth to induce CRISPRi repression. The culture samples were centrifuged at 4700 RPM for 5 min to produce a cell pellet, which was washed twice using PBS for 10 min, with 2 min of centrifugation at 12000 RPM between each wash. Following removal of media and PBS buffer, 1 mL of glutaraldehyde was added to fix the sample. Samples remained in fixative at least overnight, then fixative was removed and replaced with 1 mL of PBS (to ensure safe transportation) before being sent to UAMS Digital Microscopy Core for TEM analysis.

### Fluorescence microscopy

Strains were cultured overnight in BHI and sub-cultured at 1:20 ratio in CDM, with and without xylose for 3 hours (exponential growth phase). After three hours, the bacterial cells were pelleted, re-suspended in sterile 0.85% NaCl solution and washed. The washed pellets were then again re-suspended in the 0.85% NaCl and 10 µg/mL DAPI followed by incubation for 10 min. The suspension was washed again to remove extra DAPI and again re-suspended in 0.85% NaCl. For the microscopy, a 1% agarose pad was prepared using SecureSeal frames (Grace Bio-Labs). 1 µl of the sample was spotted onto the agarose pad, sealed with a coverslip and observed using a Nikon Eclipse Ti2 inverted confocal microscope. The images were taken using a 100x objective and zoomed to 3x for visual clarity. Image analysis and cell counting was performed using Nikon Imaging Software (NIS elements).

### sgRNA design and plasmid construction

Target-specific 20-nt sgRNA sequences, against SMU_734, SMU_775c, and SMU_1802c, were identified using CRISPy-web (61) based on the *S. mutans* UA159 reference genome (GenBank NC_004350.2). To mitigate potential off-target effects, sequences were filtered for predicted matches within the 13-bp region distal to the PAM. Only sgRNAs with two or more mismatches to any predicted off-target site were retained, as we previously determined that such divergence significantly diminishes off- target activity (4). Selection was prioritized for sequences in close proximity to the translation initiation site of each target gene. Furthermore, the secondary structure of each potential guide, including the 20-nt targeting sequence and the 42-nt dCas9 binding scaffold, was modeled via the ViennaRNA RNAfold program (62). Only sequences that maintained the requisite hairpin architecture of the dCas9 binding handle were processed for cloning; notably, these rigorous criteria precluded the identification of viable sgRNAs for SMU_734. Construct assembly was performed using the Q5 Site-Directed Mutagenesis Kit (New England Biolabs). Primers were designed using NEBaseChanger (v.2.7.2) to substitute the default *gfp*-targeting sequence with each unique 20-nt guide. Following PCR amplification of the pPM_sgRNA backbone, products were treated with a Kinase-Ligase-DpnI (KLD) enzyme mix for 5 min and transformed into chemically competent *E. coli* 10-beta. Kanamycin-resistant transformants were verified via Sanger sequencing (pRCS1seqR primer; Table S11) and archived as glycerol stocks at -80°C. Despite repeated attempts, including designing alternative sgRNA primers, we were not able to obtain an sgRNA targeting SMU_775c.

### Validation of gene essentiality and identification of genetic suppressors

To verify the essentiality of the targeted loci and identify potential bypass mechanisms, we attempted to replace each gene with a non-polar kanamycin resistance marker via homologous recombination. Genetic challenges were performed in two backgrounds: wild-type *S. mutans* UA159 and a high-density transposon mutant library. The inclusion of the Tn-mutagenized background was designed to provide a diverse genetic landscape capable of providing “escape” routes, either through secondary mutations or specific Tn-disruptions, that facilitate survival in the absence of an essential gene. Mutants were engineered using a PCR ligation mutagenesis strategy (63). Briefly, 500– 600 bp flanking regions were amplified using primers A/B (upstream) and C/D (downstream), with approximately 50 bp of overlap with the target coding sequence. Primers B and C contained BamHI sites for the ligation of flanking fragments to a non-polar kanamycin cassette harvested from plasmid pALH124. Following transformation and selection on BHI-kanamycin agar, the absence of transformants in the wild-type background was used as a primary indicator of essentiality. Rare “escape” colonies recovered from either the wild-type or Tn-library backgrounds were subjected to whole-genome sequencing to identify suppressive variants. Genomic DNA was isolated using the MasterPure Gram-Positive DNA purification kit (Epicentre) and sequenced via SeqCenter (Illumina; 200 Mbp package). Variant calling and the identification of chromosome rearrangements or transposon insertion sites were performed using the Breseq software package (v.0.35.0) against the *S. mutans* UA159 reference genome (AE014133.2).

### RNA isolation and RNA sequencing

Bacterial cultures were grown to mid-log phase (OD_600_ ∼0.4-0.6) with appropriate concentrations of xylose (w/v) in CDM to induce CRISPRi repression. The culture samples were centrifuged to produce a cell pellet, and RNAprotect was added to stabilize the RNA. Samples were then incubated for 5 min at room temperature, the RNAprotect was removed, and samples were frozen at -80°C. Next, frozen samples were placed on ice to prevent RNA degradation, and all surfaces and instruments were sprayed with RNase Zap to prevent RNA degradation by environmental RNases. Samples were transferred to tubes containing glass beads, acidic phenol was added to the samples to lyse cells, then cells were resuspended in 280 μL of 50/10 Tris EDTA buffer with 0.4% sodium dodecyl sulfate (SDS) to prevent digestion of nucleic acids. A bead beater was used for 30 sec at homogenizing speed, then samples were placed on ice for 3 min. This process was repeated three times, then samples were centrifuged at 14,000 RPM for 10 min at 4°C. Subsequent extraction steps used a Qiagen RNeasy extraction kit following the manufacturer’s instructions. RNA concentration was checked using a DeNovix QFX Fluorometer and Qubit RNA Broad Range (BR) assay kit. In total, three biological replicates were produced for each CRISPRi strain. Next, RNA was sent to SeqCenter who generated RNA-seq libraries using Illumina stranded RNA library preparation with RiboZero Plus rRNA depletion. Sequencing was conducted on an Illumina platform providing up to 12 million paired end reads (2×51 bp) per sample. After sequencing, SeqCenter conducted a basic RNA-seq analysis pipeline that provided raw transcript level quantification. Afterwards, gene expression changes between samples were quantified with Degust (http://degust.erc.monash.edu/) using the edgeR methodology (64).

### Tandem mass tag mass spectrometry

Bacterial cultures were grown to mid-log phase (OD_600_ ∼0.4-0.6) with appropriate concentrations of xylose (w/v) in CDM to induce CRISPRi repression. The culture samples were centrifuged to produce a cell pellet, and the pellets were sent to the IDeA Proteomics Center at the University of Arkansas for Medical Sciences for mass spectrometry. There, small peptides were produced from the samples using chloroform/methanol extraction with trypsin digestion. High-pH peptide fractionation was performed to separate peptides, then mass spectrometry was performed on an Orbitrap Eclipse (TMT MS3; 60 min gradient per fraction). Quality control, normalization, and differential expression analysis were also completed by the IDeA Proteomics Center.

### Site-directed mutagenesis

A splice-overlap extension (SOE) PCR mediated homologous recombination method was used to introduce a point mutation into *dnaA* (SMU_01) gene that substituted glutamine (CAA) at the 197th amino acid to glutamate (GAA) (*dnaA*^Q197E^) (65). PCR amplifications were performed using NEB Q5 HF Master Mix according to the manufacturer’s instructions. In the first PCR step, two overlapping fragments were generated using primer pairs dnaA_Q197E_A/ dnaA_Q197E_B and dnaA_Q197E_C/ dnaA_Q197E_D (Table S11). The two products were subsequently fused by a second overlap-extension PCR using dnaA_Q197E_A and dnaA_Q197E_D to generate the full-length recombinant fragment containing the Q197E mutation. This recombinant PCR fragment was purified, then transformed into *S. mutans* strains using competence-stimulating peptide and the pLacG suicide integration plasmid carrying an erythromycin resistance marker (66). Colonies were screened by colony PCR using primers dnaA_Q197E_A and dnaA_Q197E_D, followed by confirmation of presence of the Q197E mutation with Sanger sequencing. Then, the mutation confirmed colonies were serially streaked twice on tryptone-vitamin (TV) agar containing 0.5% lactose as the sole carbohydrate to obtain markerless *dnaA*^Q197E^ mutant, which has lost the suicide integration plasmid due to spontaneous excision. Q197E mutation was again validated in the markerless mutant with colony PCR and Sanger Sequencing.

## Data availability

The RNA-seq transcriptomic datasets have been deposited in the NCBI Gene Expression Omnibus (GEO) database and are accessible under the GEO series accession number GSE338288. The quantitative proteomic datasets and metadata have been deposited in Figshare at https://doi.org/10.6084/m9.figshare.32948528.

## Acknowledgements

This work was supported by the Arkansas INBRE program [NIGMS P20 GM103429] (RCS). This work was also supported by the National Institute of Dental and Craniofacial Research (NIDCR) grants DE033403 and DE036253 (RCS).

