## Supplemental_Materials for "Identification of a putative RocS homolog through phenotypic profiling of uncharacterized essential genes in *Streptococcus mutans*"

**This file includes**:

Figures S1 - S4

Tables S1-S11

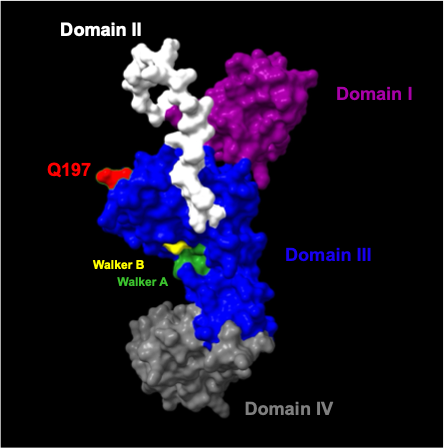

**Figure S1 Structural and functional architecture of the *Streptococcus mutans* DnaA protein.** The protein is composed of four distinct functional domains: Domain I (protein-protein interactions), Domain II (flexible linker), Domain III (AAA+ ATPase), and Domain IV (dsDNA binding). The critical ATP-binding Walker A (GGPGLGKT; residues 152–159) and ATP-hydrolyzing Walker B (DLLLIDDI; residues 210–217) motifs are highlighted within Domain III. In the AlphaFold-predicted structure (AF-Q8DWN9-F1), the Q197 residue (red) is situated on the outside of the DnaA protein within Domain III.

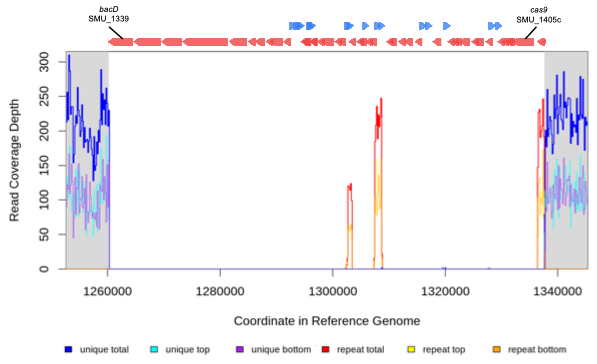

**Figure S2 Characterization of the TnSmu2 genomic island deletion.** Illumina read depth is plotted against the coordinates of the *Streptococcus mutans* UA159 reference genome (x-axis). The y-axis represents the read coverage depth, with the baseline expected at approximately 200x. A significant loss of unique read coverage is observed between coordinates ~1,260,000 and ~1,340,000, indicating a large-scale genomic deletion of the Tn*Smu2* island in the sequenced isolate. Red and blue arrows at the top of the panel represent the orientation and position of predicted open reading frames (ORFs) within the reference genome. The lack of reads directly correlates with the region encompassing the Tn*Smu2* element and a ~20 kb 3′ flanking region.

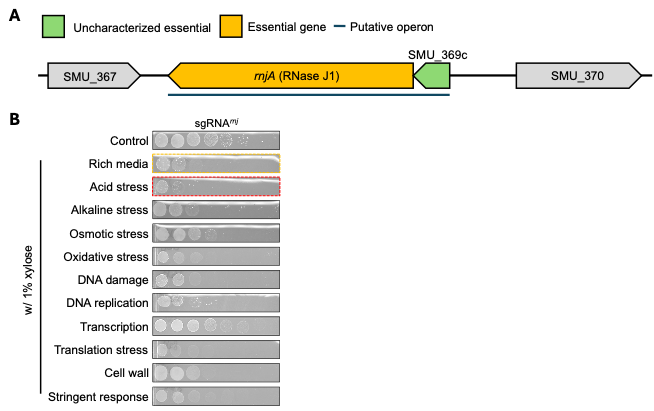

**Figure S3 Genomic organization and validation of essentiality for SMU_368/*rnjA*.** (A) Map of the SMU_369c and *rnjA* (SMU_368) genetic locus. Arrows indicate the genomic orientation and relative positions of the target genes in *S. mutans* UA159. SMU_368 encodes the essential ribonuclease J1 (*rnjA*), which is transcriptionally linked to the upstream gene SMU_369c. (B) Phenotypic validation of SMU_368/*rnjA* gene essentiality. Ten-fold serial dilutions of the indicated strains were spotted onto solid rich medium in the absence (control) or presence (0.5% xylose) of the dCas9 inducer. Plates were supplemented with the following stressors: acid (pH 5.5), alkaline (pH 8.5), osmotic (2.5% NaCl), oxidative (1 mM H2O2), DNA damage (10 ng/mL mitomycin C), DNA replication (1.5 µg/mL ciprofloxacin), transcription (2 µg/mL rifampicin), translation (2 µg/mL chloramphenicol), cell wall (2 µg/mL ampicillin), and the stringent response (125 ng/mL mupirocin). Red and orange boxes show conditions where *rnjA* repression showed the highest and second highest sensitivity to those conditions respectively.

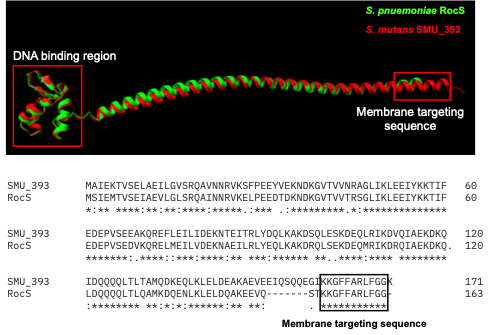

**Figure S4 Structural characterization of *S. mutans* RocS (SMU_393).** Predicted structure of SMU_393 and RocS generated via AlphaFold2 (AF-Q8DWN9-F1 and AF-Q8DQ15-F1) and visualized in ChimeraX. The model reveals a prominent C-terminal amphipathic alpha-helix, characteristic of membrane-anchoring domains in chromosome segregation proteins. Sequence alignment (bottom) highlights the conservation of this MTS between *S. mutans* and *S. pneumoniae*.

**Table S1 Bioinformatic characterization, evolutionary conservation, and predicted functional assignments of the nine uncharacterized essential gene targets**

| **GENE** | **SMU_1801c** | **SMU_1802c** | **SMU_369c** | **SMU_393** | **SMU_415** | **SMU_419** | **SMU_734** | **SMU_775c** | **SMU_958** |
| --- | --- | --- | --- | --- | --- | --- | --- | --- | --- |
| **Uniprot ID** | Q8DSI4 | Q8DSI3 | Q8DVU6 | Q8DVS4 | Q8DVQ5 | Q8DVQ1 | Q8DUZ3 | Q8DUW6 | Q8DUH1 |
| **GenBank Annotation** | putative GTP-binding protein | conserved hypothetical protein | conserved hypothetical protein | conserved hypothetical protein | conserved hypothetical protein | conserved hypothetical protein | conserved hypothetical protein | conserved hypothetical protein | hypothetical protein |
| **Experimentally Validated Homolog** | GTP-binding protein (YqeH) | phosphatidylglycerol phosphate phosphatase (PgpP) | DNA-directed RNA polymerase subunit epsilon (RpoY) | Regulator of Chromosome Segregation (RocS) | Cell cycle regulator (CcrZ) | RNase P modulator (RnpM) | - | PG-dependent type I LTA synthase (LtaS) | - |
| **Homolog Uniprot ID** | P54453 | P54452 | O31718 | Q8DQ15 | A0A0H2ZQL5 | P32728 | - | Not Available; Q7A1I3 | - |
| **Species** | *B. subtilis* | *B. subtilis* | *B. subtilis* | *S. pneumoniae* | *S. pneumoniae* | *B. subtilis* | - | *S. mitis*; *S. aureus* | - |
| **Alignment** | 58% identity, 100% coverage | 38% identity, 91% coverage | 44% identity, 99% coverage | 76% identity, 99% coverage | 60% identity, 98% coverage | 48% identity, 94% coverage | - | 61% identity, 98% coverage; 41% identity, 90% coverage | - |
| **PMID** | PMID: 17895579 | PMID: 39869797 | PMID: 25092033 | PMID: 31182798 | PMID: 34373624 | PMID: 38050972 | - | PMID: 33627509; PMID: 17483484 | - |
| **Function** | YqeH is an essential GTPase involved in 30S ribosomal subunit assembly | PgpP catalyzes the terminal step of the phosphatidylglycerol lipid synthesis pathway | RpoY is a non-essential RNA polymerase subunit found in Firmicutes with a putative role in protection from bacteriophage infection | RocS is a chromosome segregation protein that interacts with ParB serving as a central coordinator of chromosome segregation and cell division | CcrZ is a cell cycle regulator that interacts with FtsZ and controls DNA replication by modulating the activity of DnaA | RnpM binds directly to the RNA component (RnpB) of RNase P, modulating its activity | Uncharacterized Protein | LtaS is a membrane-associated enzyme that synthesizes type I lipoteichoic acid (LTA) | Annotation Artifact |
| **GO (Molecular Function)** | GTP Binding [GO:0005525]; GTPase activity [GO:0003924] | Phosphatidylglycerophosphatase activity [GO:0008962] | DNA binding [GO:0003677]; DNA-directed RNA polymerase complex binding [GO:1990391] | DNA binding [GO:0003677] | Protein binding [GO:0005515] | RNA binding [GO:0003723] | - | Lipoteichoic acid synthase activity [GO:0070395] | - |
| **GO (Biological Process)** | Ribosome biogenesis [GO:0042254]; Ribosomal small subunit assembly [GO:0000028] | Phospholipid biosynthetic process [GO:0008654]; Phosphatidylglycerol biosynthetic process [GO:0006658] | DNA-templated transcription [GO:0006351] | Chromosome segregation [GO:0007059]; Cell division [GO:0051301] | DNA replication initiation [GO:0006270] | tRNA processing [GO:0008033] | - | Lipoteichoic acid biosynthetic process [GO:0070394] | - |

**Table S2 Differential gene expression (RNA-seq) analysis of *S. mutans* UA159 during CRISPRi-mediated knockdown of SMU_368c/rnjA**

| **Locus tag** | **Gene** | **Description** | **log2 fold change** | **FDR** |
| --- | --- | --- | --- | --- |
| SMU_2124 |  | hypothetical protein | 6.41 | 3.90E-08 |
| SMU_1405c | cas9 | conserved hypothetical protein | 6.37 | 1.41E-124 |
| SMU_2053c |  | hypothetical protein | 4.33 | 7.61E-09 |
| SMU_1310 |  | hypothetical protein | 4.31 | 2.42E-07 |
| SMU_1896c |  | hypothetical protein | 3.18 | 5.27E-25 |
| SMU_958 |  | hypothetical protein | 3.07 | 3.26E-05 |
| SMU_11 |  | conserved hypothetical protein | 2.92 | 1.64E-06 |
| SMU_1895c |  | hypothetical protein | 2.67 | 1.80E-16 |
| SMU_2146c |  | hypothetical protein | 2.65 | 1.80E-66 |
| SMU_1000 |  | hypothetical protein | 2.64 | 1.57E-03 |
| SMU_1862 |  | hypothetical protein | 2.49 | 1.98E-26 |
| SMU_1282 |  | putative transcriptional regulator | 2.49 | 3.40E-34 |
| SMU_88c |  | conserved hypothetical protein; possible mechanosensitive ion channel | 2.44 | 6.01E-64 |
| SMU_1754c |  | conserved hypothetical protein | 2.43 | 1.24E-26 |
| SMU_1804c |  | hypothetical protein | 2.39 | 1.03E-05 |
| SMU_393 |  | conserved hypothetical protein | 2.37 | 3.07E-30 |
| SMU_916c |  | conserved hypothetical protein | 2.29 | 7.44E-17 |
| SMU_1628 |  | conserved hypothetical protein | 2.27 | 3.09E-22 |
| SMU_1757c |  | conserved hypothetical protein | 2.20 | 8.11E-20 |
| SMU_919c |  | putative ATPase, confers aluminum resistance | 2.15 | 1.76E-12 |
| SMU_917c |  | putative 6-pyruvoyl tetrahydropterin synthase | 2.13 | 3.34E-13 |
| SMU_727 |  | putative transcriptional regulator | 2.13 | 3.23E-22 |
| SMU_27 | acpP | putative acyl carrier protein; AcpP; ACP | 2.13 | 3.76E-14 |
| SMU_876 | msmR | putative MSM operon regulatory protein | 2.12 | 8.03E-23 |
| SMU_2125 |  | conserved hypothetical protein | 2.11 | 7.12E-32 |
| SMU_439 |  | putative transcriptional regulator | 2.10 | 2.07E-25 |
| SMU_2070 |  | conserved hypothetical protein | 2.07 | 2.71E-29 |
| SMU_977 | licT | putative transcriptional antiterminator LicT (fragment) | 2.06 | 9.43E-52 |
| SMU_595 | pyrD | putative dihydroorotate dehydrogenase; dihydroorotate oxidase | 2.04 | 6.74E-35 |
| SMU_2054c |  | conserved hypothetical protein | 2.04 | 8.42E-32 |
| SMU_1409c |  | putative transcriptional regulator | 2.02 | 3.48E-27 |
| SMU_933 |  | putative amino acid ABC transporter, periplasmic amino acid-binding protein | 1.98 | 1.31E-05 |
| SMU_2052c |  | hypothetical protein | 1.97 | 1.52E-21 |
| SMU_2129c |  | conserved hypothetical protein | 1.97 | 2.28E-17 |
| SMU_934 |  | putative amino acid ABC transporter, permease protein | 1.96 | 8.73E-05 |
| SMU_1064c |  | putative transcriptional regulator (GntR family) | 1.95 | 1.33E-18 |
| SMU_932 |  | hypothetical protein | 1.94 | 6.04E-05 |
| SMU_1775c |  | hypothetical protein | 1.94 | 2.86E-17 |
| SMU_576 | lytR | putative response regulator LytR | 1.93 | 5.49E-24 |
| SMU_616 |  | hypothetical protein | 1.92 | 2.44E-07 |
| SMU_1023 | pycB | putative pyruvate carboxylase/oxaloacetate decarboxylase, alpha subunit | 1.92 | 3.24E-53 |
| SMU_1592 | pepQ | putative dipeptidase PepQ | 1.92 | 8.51E-20 |
| SMU_1753c |  | conserved hypothetical protein | 1.92 | 1.99E-17 |
| SMU_223c |  | hypothetical protein | 1.90 | 7.86E-14 |
| SMU_2123 |  | hypothetical protein | 1.88 | 9.98E-27 |
| SMU_239c |  | hypothetical protein | 1.88 | 2.08E-16 |
| SMU_281 |  | hypothetical protein | 1.88 | 6.72E-11 |
| SMU_86 |  | conserved hypothetical protein | 1.88 | 4.38E-23 |
| SMU_662 |  | conserved hypothetical protein; possible membrane protein | 1.88 | 8.86E-33 |
| SMU_999 |  | hypothetical protein | 1.87 | 1.99E-10 |
| SMU_68 |  | hypothetical protein | 1.86 | 6.84E-27 |
| SMU_1022 | citG2 | conserved hypothetical protein, CitG-like protein | 1.86 | 3.20E-36 |
| SMU_1021 | cilA | putative citrate lyase, alfa subunit | 1.84 | 2.04E-38 |
| SMU_1315c |  | putative ATP-binding protein | 1.84 | 1.09E-33 |
| SMU_2133c |  | putative membrane protein | 1.83 | 1.99E-10 |
| SMU_2148c |  | conserved hypothetical protein; possible cobalt permease | 1.83 | 7.21E-36 |
| SMU_382c |  | putative oxidoreductase | 1.81 | 2.20E-10 |
| SMU_1398 |  | putative transcriptional regulator | 1.81 | 3.60E-19 |
| SMU_722 |  | hypothetical protein | 1.81 | 5.87E-20 |
| SMU_73 |  | conserved hypothetical protein | 1.80 | 1.29E-22 |
| SMU_277 |  | hypothetical protein | 1.80 | 3.73E-21 |
| SMU_930c |  | putative transcriptional regulator | 1.79 | 6.72E-11 |
| SMU_1224 | pyrK | putative dihydroorotate dehydrogenase, electron transfer subunit | 1.77 | 5.78E-19 |
| SMU_383c |  | conserved hypothetical protein; putative reductase | 1.77 | 3.01E-10 |
| SMU_1402c |  | conserved hypothetical protein | 1.77 | 1.19E-09 |
| SMU_915c |  | conserved hypothetical protein | 1.75 | 9.31E-11 |
| SMU_2050c |  | putative methyltransferase | 1.75 | 5.57E-37 |
| SMU_1020 | cilB | putative citrate lyase CilB, citryl-CoA lyase, beta subunit | 1.75 | 2.26E-27 |
| SMU_998 |  | putative ABC transporter, periplasmic ferrichrome-binding protein | 1.73 | 6.47E-53 |
| SMU_497c |  | conserved hypothetical protein | 1.73 | 1.43E-21 |
| SMU_1211 |  | conserved hypothetical protein | 1.72 | 1.11E-26 |
| SMU_1755c |  | conserved hypothetical protein | 1.70 | 2.12E-13 |
| SMU_72 |  | conserved hypothetical protein | 1.70 | 1.35E-15 |
| SMU_983 | bglC | putative transcriptional regulator | 1.69 | 3.89E-25 |
| SMU_1789c |  | conserved hypothetical protein | 1.69 | 4.05E-25 |
| SMU_1766c |  | hypothetical protein | 1.69 | 5.90E-25 |
| SMU_1897 |  | putative ABC transporter, ATP-binding protein | 1.68 | 3.82E-20 |
| SMU_378 |  | hypothetical protein | 1.66 | 3.18E-05 |
| SMU_661 |  | putative transcriptional regulator | 1.65 | 3.09E-22 |
| SMU_1539 | glgB | putative 1,4-alpha-glucan branching enzyme | 1.65 | 1.28E-07 |
| SMU_1019 | cilG | putative citrate lyase, gamma-subunit | 1.64 | 1.17E-24 |
| SMU_911c |  | hypothetical protein | 1.64 | 3.08E-21 |
| SMU_112c |  | putative transcriptional regulator | 1.64 | 7.00E-19 |
| SMU_1254 |  | conserved hypothetical protein | 1.63 | 4.40E-21 |
| SMU_976 | potD | putative ABC transporter, periplasmic spermidine/putrescine-binding protein | 1.63 | 2.37E-20 |
| SMU_935 |  | putative amino acid ABC transporter, permease protein | 1.62 | 2.52E-02 |
| SMU_1621c |  | conserved hypothetical protein | 1.62 | 4.13E-11 |
| SMU_253 | dacA | putative D-alanyl-D-alanine carboxypeptidase; penicillin-binding protein | 1.62 | 1.55E-24 |
| SMU_1487 |  | conserved hypothetical protein | 1.62 | 6.12E-14 |
| SMU_2066c |  | putative transmembrane protein | 1.62 | 4.15E-40 |
| SMU_283 |  | hypothetical protein | 1.61 | 9.81E-17 |
| SMU_936 |  | putative amino acid ABC transporter, ATP-binding protein | 1.60 | 1.71E-03 |
| SMU_270 | sgaT | putative PTS system, membrane component; possible ribulose-monophosphate PTS pathway enzyme IIC | 1.60 | 1.29E-12 |
| SMU_1219c |  | conserved hypothetical protein | 1.58 | 3.83E-23 |
| SMU_728 |  | putative oxidoreductase | 1.56 | 5.00E-18 |
| SMU_1316c |  | hypothetical protein | 1.56 | 3.47E-14 |
| SMU_87 |  | conserved hypothetical protein | 1.55 | 2.68E-13 |
| SMU_787 |  | putative transcriptional regulator | 1.55 | 5.05E-32 |
| SMU_440 |  | hypothetical protein | 1.54 | 1.31E-22 |
| SMU_573 |  | conserved hypothetical protein | 1.54 | 2.56E-27 |
| SMU_660 |  | putative histidine kinase SpaK | 1.54 | 4.59E-25 |
| SMU_1052 |  | conserved hypothetical protein | 1.53 | 4.33E-26 |
| SMU_05 |  | conserved hypothetical protein | 1.53 | 4.30E-16 |
| SMU_24 |  | putative amino acid aminotransferase | 1.53 | 5.30E-17 |
| SMU_914c |  | conserved hypothetical protein | 1.53 | 2.19E-08 |
| SMU_2049c |  | conserved hypothetical protein | 1.52 | 1.89E-23 |
| SMU_340 |  | 50S ribosomal protein L34 | 1.51 | 8.59E-20 |
| SMU_2065 |  | putative UDP-glucose 4-epimerase | 1.51 | 2.20E-30 |
| SMU_974 | potB | putative spermidine/putrescine ABC transporter, permease protein | 1.50 | 9.44E-30 |
| SMU_659 |  | putative response regulator SpaR | 1.50 | 3.15E-19 |
| SMU_1660c |  | conserved hypothetical protein | 1.50 | 4.66E-16 |
| SMU_577 | lytS | putative histidine kinase LytS | 1.49 | 1.44E-11 |
| SMU_1223 | pyrDB | putative dihydroorotate dehydrogenase B | 1.49 | 6.16E-12 |
| SMU_1317c |  | hypothetical protein | 1.49 | 2.12E-13 |
| SMU_89c |  | putative nitrite transporter | 1.48 | 3.68E-10 |
| SMU_997 |  | putative inorganic ion ABC transporter, ATP-binding protein; possible ferrichrome transport system | 1.47 | 4.13E-31 |
| SMU_2120c |  | putative 3-methyladenine DNA glycosylase | 1.47 | 1.33E-17 |
| SMU_982 | bglB2 | putative BglB fragment | 1.46 | 4.77E-05 |
| SMU_2069 |  | putative integral membrane protein | 1.46 | 7.01E-15 |
| SMU_1950 |  | putative pseudouridylate synthase | 1.45 | 5.11E-22 |
| SMU_2040 | treR | putative transcriptional regulator; repressor of the trehalose operon | 1.45 | 9.56E-23 |
| SMU_25 | recO | putative DNA repair protein RecO | 1.44 | 4.07E-12 |
| SMU_1372c |  | hypothetical protein | 1.44 | 4.97E-09 |
| SMU_1415c |  | putative phosphatases involved in N-acetyl-glucosamine catabolism | 1.44 | 1.26E-32 |
| SMU_220c |  | hypothetical protein | 1.43 | 1.07E-14 |
| SMU_1891c |  | hypothetical protein | 1.42 | 8.70E-04 |
| SMU_267c |  | putative glutamate-cysteine ligase | 1.41 | 8.94E-15 |
| SMU_255 | oppA | putative oligopeptide ABC transporter, substrate-binding protein OppA | 1.41 | 7.06E-19 |
| SMU_2149c |  | putative ABC transporter, ATP-binding protein; possible cobalt transport system | 1.40 | 2.56E-27 |
| SMU_1538 | glgC | putative glucose-1-phosphate adenylyltransferase; ADP-glucose pyrophosphorylase | 1.40 | 2.32E-05 |
| SMU_975 | potC | putative spermidine/putrescine ABC transporter, permease protein | 1.39 | 6.64E-27 |
| SMU_1065c |  | putative transcriptional regulator (GntR family) | 1.39 | 2.64E-15 |
| SMU_1213c |  | putative 5'-nucleotidase precursor | 1.39 | 1.16E-19 |
| SMU_768c |  | conserved hypothetical protein | 1.38 | 2.18E-11 |
| SMU_2150c |  | putative ABC transporter; ATP-binding protein; possible cobalt transport system | 1.38 | 2.20E-24 |
| SMU_637c |  | hypothetical protein | 1.37 | 8.83E-16 |
| SMU_496 | cysK | putative cysteine synthetase A; O-acetylserine lyase | 1.35 | 3.86E-06 |
| SMU_1251 |  | conserved hypothetical protein | 1.34 | 7.71E-15 |
| SMU_1319c |  | conserved hypothetical protein | 1.34 | 2.03E-23 |
| SMU_258 | oppD | putative oligopeptide ABC transporter, ATP-binding protein OppD | 1.34 | 2.91E-24 |
| SMU_1774c |  | conserved hypothetical protein | 1.34 | 3.58E-08 |
| SMU_1016 | bcc | putative acetyl-CoA carboxylase, biotin carboxyl carrier subunit | 1.34 | 6.29E-19 |
| SMU_259 | oppF | putative oligopeptide ABC transporter, ATP-binding protein OppF | 1.33 | 2.25E-26 |
| SMU_85 | thiD | putative phosphomethylpyrimidine kinase | 1.33 | 2.19E-16 |
| SMU_1991 | pbp1b | putative membrane carboxypeptidase, penicillin-binding protein 1b | 1.32 | 1.76E-31 |
| SMU_1661c |  | putative signal peptidase II | 1.31 | 7.54E-16 |
| SMU_948 |  | conserved hypothetical protein | 1.31 | 1.47E-08 |
| SMU_84 | truA | putative tRNA pseudouridine synthase A | 1.31 | 3.86E-15 |
| SMU_721 |  | conserved hypothetical protein | 1.30 | 8.38E-21 |
| SMU_1037c |  | putative histidine kinase | 1.30 | 1.38E-22 |
| SMU_799c |  | conserved hypothetical protein | 1.29 | 2.84E-06 |
| SMU_878 | msmE | multiple sugar-binding ABC transporter, sugar-binding protein precursor MsmE | 1.29 | 2.49E-10 |
| SMU_1941 | atmB | putative membrane lipoprotein | 1.29 | 3.22E-12 |
| SMU_1445c |  | putative ABC transporter, ATP-binding protein | 1.28 | 1.34E-23 |
| SMU_1200 | rs1 | putative ribosomal protein S1; sequence specific DNA-binding protein | 1.28 | 6.55E-34 |
| SMU_1844 | scrR | sucrose operon repressor | 1.28 | 1.22E-11 |
| SMU_1664c |  | putative acetoin utilization protein, acetoin dehydrogenase | 1.28 | 1.45E-14 |
| SMU_26 | plsX | putative fatty acid/phospholipid synthesis protein | 1.27 | 3.64E-20 |
| SMU_1918 | dedA | putative membrane-associated protein DedA | 1.27 | 1.05E-17 |
| SMU_278 |  | hypothetical protein | 1.27 | 2.11E-09 |
| SMU_1018 |  | hypothetical protein | 1.27 | 1.07E-11 |
| SMU_1417c |  | putative oleoyl-acyl carrier protein thioesterase | 1.27 | 1.19E-18 |
| SMU_1977c |  | putative transcriptional regulator | 1.27 | 8.26E-17 |
| SMU_1404c |  | conserved hypothetical protein | 1.27 | 3.17E-06 |
| SMU_428 |  | conserved hypothetical protein | 1.27 | 1.49E-23 |
| SMU_1416c |  | putative mutator protein MutT | 1.26 | 8.87E-26 |
| SMU_877 | agaL | alpha-galactosidase | 1.26 | 6.16E-11 |
| SMU_939 |  | putative dehydrogenase (FMN-dependent family protein) | 1.25 | 1.83E-24 |
| SMU_653c |  | putative ABC transporter, permease protein | 1.25 | 8.99E-11 |
| SMU_1760c |  | conserved hypothetical protein | 1.25 | 1.60E-07 |
| SMU_885 | galR | galactose operon repressor GalR | 1.24 | 8.77E-17 |
| SMU_1572 | murZ | putative UDP-N-acetylglucosamine-1-carboxyvinyl transferase | 1.23 | 1.26E-15 |
| SMU_1536 | glgA | putative starch (bacterial glycogen) synthase | 1.23 | 4.42E-05 |
| SMU_773c |  | lysyl-tRNA synthetase | 1.23 | 1.18E-21 |
| SMU_1832 |  | hypothetical protein | 1.22 | 1.22E-11 |
| SMU_1403c |  | conserved hypothetical protein | 1.22 | 2.07E-05 |
| SMU_782 |  | conserved hypothetical protein | 1.21 | 1.53E-22 |
| SMU_457 |  | hypothetical protein | 1.21 | 1.58E-03 |
| SMU_720 |  | conserved hypothetical protein; possible Na+/solute symporter | 1.20 | 6.49E-19 |
| SMU_1915 | comC | competence stimulating peptide, precursor | 1.20 | 9.74E-05 |
| SMU_1414c |  | conserved hypothetical protein | 1.19 | 1.38E-15 |
| SMU_784 | aroA | 5-enolpyruvylshikimate-3-phosphate synthase | 1.19 | 2.62E-12 |
| SMU_973 | potA | putative spermidine/putrescine ABC transporter, ATP-binding protein | 1.19 | 7.61E-26 |
| SMU_947 | dfrA | putative dihydrofolate reductase | 1.18 | 2.44E-11 |
| SMU_1761c |  | conserved hypothetical protein | 1.18 | 2.32E-05 |
| SMU_1150 |  | putative transporter, trans-membrane domain bacteriocin immunity protein | 1.18 | 3.48E-03 |
| SMU_1217c |  | putative ABC transporter, amino acid binding protein | 1.17 | 4.05E-11 |
| SMU_1961c |  | putative PTS system, sugar-specific enzyme IIA component | 1.16 | 6.67E-05 |
| SMU_652c |  | putative ABC transporter, ATP-binding protein; possible nitrate transport system | 1.16 | 2.65E-08 |
| SMU_1178c |  | putative amino acid ABC transporter, ATP-binding protein | 1.16 | 7.22E-20 |
| SMU_1678 |  | conserved hypothetical protein, possible acyl-CoA thioesterase | 1.16 | 3.09E-19 |
| SMU_318 |  | putative hippurate hydrolase | 1.15 | 2.73E-15 |
| SMU_257 | oppC | putative transmembrane protein, permease OppC | 1.15 | 4.07E-14 |
| SMU_456 | mraY | putative undecaprenyl-phosphate-UDP-MurNAc-pentapeptide transferase | 1.15 | 4.37E-24 |
| SMU_1535 | phsG | glycogen phosphorylase | 1.15 | 6.64E-07 |
| SMU_1177c |  | putative ABC transporter, glutamine binding protein | 1.14 | 8.34E-18 |
| SMU_339 |  | hypothetical protein | 1.14 | 2.27E-16 |
| SMU_1622 | pmsR | putative peptide methionine sulfoxide reductase | 1.14 | 1.70E-10 |
| SMU_1662 | holB | putative DNA polymerase III, delta subunit | 1.14 | 3.59E-21 |
| SMU_442 |  | conserved hypothetical protein | 1.13 | 1.99E-13 |
| SMU_627 |  | conserved hypothetical protein | 1.13 | 3.57E-12 |
| SMU_1604c |  | conserved hypothetical protein | 1.13 | 8.51E-06 |
| SMU_1767c |  | hypothetical protein | 1.13 | 3.66E-13 |
| SMU_996 |  | putative ABC transporter, permease protein; possible ferrichrome transport system | 1.12 | 2.34E-15 |
| SMU_1758c |  | conserved hypothetical protein | 1.12 | 1.76E-07 |
| SMU_1675 | metB | putative cystathionine gamma-synthase; possible bifunctional enzyme | 1.12 | 3.56E-12 |
| SMU_1641c |  | conserved hypothetical protein | 1.12 | 8.84E-05 |
| SMU_1406c |  | conserved hypothetical protein | 1.12 | 1.17E-10 |
| SMU_518 |  | conserved hypothetical protein | 1.11 | 1.58E-19 |
| SMU_1218 | nylA | putative amidase | 1.11 | 8.39E-16 |
| SMU_2147c |  | conserved hypothetical protein | 1.11 | 7.12E-13 |
| SMU_1812 |  | putative transposase, ISSmu2 | 1.10 | 1.28E-02 |
| SMU_241c |  | putative ABC transporter, ATP-binding protein; amino acid transport system | 1.09 | 1.85E-07 |
| SMU_849 |  | 50S ribosomal protein L27 | 1.09 | 1.43E-11 |
| SMU_781 |  | putative prephenate dehydrogenase | 1.09 | 1.15E-20 |
| SMU_1663 | kthY | putative thymidylate kinase | 1.09 | 6.12E-12 |
| SMU_1889c |  | hypothetical protein | 1.09 | 2.28E-07 |
| SMU_694c |  | putative ferredoxin (4Fe-4S) | 1.09 | 2.06E-03 |
| SMU_256 | oppB | putative oligopeptide transport system, permease protein OppB | 1.08 | 4.16E-09 |
| SMU_395 | pepX | X-prolyl dipeptidyl peptidase | 1.08 | 1.72E-20 |
| SMU_588 |  | conserved hypothetical protein | 1.08 | 5.18E-09 |
| SMU_1797c |  | conserved hypothetical protein | 1.07 | 3.01E-09 |
| SMU_225c |  | hypothetical protein | 1.07 | 4.89E-06 |
| SMU_502 |  | conserved hypothetical protein | 1.07 | 1.43E-11 |
| SMU_515 |  | putative 67 kDa myosin-crossreactive streptococcal antigen | 1.07 | 5.20E-10 |
| SMU_1051 |  | putative iron-sulfur cofactor synthesis protein; NifS family | 1.06 | 9.24E-15 |
| SMU_1748 | akh | putative aspartokinase | 1.06 | 1.09E-11 |
| SMU_1975c |  | conserved hypothetical protein; possible membrane protein | 1.06 | 2.42E-17 |
| SMU_879 | msmF | multiple sugar-binding ABC transporter, permease protein MsmF | 1.05 | 1.14E-06 |
| SMU_1017 | oadB | putative oxaloacetate decarboxylase, sodium ion pump subunit | 1.05 | 2.10E-14 |
| SMU_1571 |  | putative ABC transporter, ATP-binding protein, MsmK-like protein | 1.05 | 6.61E-11 |
| SMU_990 | dapA | putative dihydrodipicolinate synthase | 1.05 | 3.18E-20 |
| SMU_711 |  | conserved hypothetical protein | 1.05 | 3.02E-07 |
| SMU_1537 | glgD | putative glycogen biosynthesis protein GlgD | 1.04 | 1.34E-03 |
| SMU_71 |  | putative cation efflux pump (multidrug resistance protein) | 1.04 | 2.49E-09 |
| SMU_2042 | dexA | dextranase precursor | 1.04 | 4.45E-16 |
| SMU_33 |  | hypothetical protein | 1.04 | 2.35E-04 |
| SMU_275 |  | putative L-ribulose 5-phosphate 4-epimerase | 1.03 | 3.33E-05 |
| SMU_1221 | pyrE | putative orotate phosphoribosyltransferase PyrE | 1.03 | 3.72E-18 |
| SMU_1839 | manA | mannose-6-phosphate isomerase | 1.03 | 3.02E-20 |
| SMU_813 |  | hypothetical protein; putative transcriptional regulator | 1.03 | 2.45E-11 |
| SMU_970 | folA | putative dihydroneopterin aldolase | 1.02 | 6.65E-13 |
| SMU_1104c |  | conserved hypothetical protein; phosphoglycerate mutase-like protein | 1.02 | 3.43E-13 |
| SMU_1447c |  | conserved hypothetical protein | 1.02 | 2.87E-14 |
| SMU_483 |  | putative phosphoprotein phosphatase (pppL protein) | 1.02 | 6.62E-10 |
| SMU_591c |  | hypothetical protein | 1.02 | 1.71E-06 |
| SMU_1855 |  | hypothetical protein | 1.02 | 3.44E-06 |
| SMU_305 |  | hypothetical protein | 1.02 | 4.61E-09 |
| SMU_859 | pyrA | putative carbamoyl phosphate synthetase, small subunit | 1.02 | 8.56E-09 |
| SMU_1673 | upp | uracil phosphoribosyltransferase | 1.02 | 4.63E-10 |
| SMU_490 | pflC | putative pyruvate formate-lyase activating enzyme | 1.02 | 3.99E-10 |
| SMU_23 | prs | phosphoribosyl pyrophosphate synthetase (PRPP synthetase) | 1.02 | 1.45E-12 |
| SMU_1014 |  | hypothetical protein | 1.01 | 1.75E-11 |
| SMU_981 | bglB1 | putative BglB fragment | 1.01 | 4.16E-06 |
| SMU_779 | aroB | putative 3-dehydroquinate synthase | 1.01 | 6.10E-18 |
| SMU_1659c |  | conserved hypothetical protein | 1.01 | 2.85E-12 |
| SMU_1289c |  | putative permease, chloride channel | 1.01 | 1.97E-11 |
| SMU_1818c |  | hypothetical protein | 1.00 | 4.02E-07 |
| SMU_2007 | rl15 | 50S ribosomal protein L15 | -1.00 | 4.51E-21 |
| SMU_321 |  | conserved hypothetical protein; possible membrane protein | -1.00 | 1.66E-10 |
| SMU_756 |  | conserved hypothetical protein | -1.00 | 1.51E-12 |
| SMU_1590 | amyA | intracellular alpha-amylase | -1.01 | 3.41E-09 |
| SMU_656 | mutE2 | putative MutE | -1.01 | 1.79E-02 |
| SMU_735 |  | hypothetical protein | -1.01 | 1.32E-04 |
| SMU_130 | adhD | putative dihydrolipoamide dehydrogenase | -1.01 | 8.87E-03 |
| SMU_929c |  | conserved hypothetical protein | -1.01 | 9.37E-03 |
| SMU_806c |  | putative glutamine ABC transporter, permease protein | -1.01 | 3.48E-10 |
| SMU_1946 |  | hypothetical protein | -1.02 | 1.70E-07 |
| SMU_14 | hprT | putative hypoxanthine-guanine phosphoribosyltransferase | -1.04 | 3.90E-12 |
| SMU_682 |  | hypothetical protein | -1.04 | 1.93E-05 |
| SMU_1169c |  | putative thioredoxin family protein | -1.04 | 4.57E-12 |
| SMU_2072c |  | conserved hypothetical protein; possible acetyltransferase | -1.05 | 2.69E-05 |
| SMU_734 |  | conserved hypothetical protein; possible membrane protein | -1.05 | 2.72E-06 |
| SMU_390 |  | hypothetical protein | -1.05 | 3.75E-02 |
| SMU_643 |  | putative esterase | -1.05 | 2.62E-05 |
| SMU_127 | adhA | putative acetoin dehydrogenase (TPP-dependent), E1 component alpha subunit | -1.05 | 3.76E-03 |
| SMU_2009 | rs5 | 30S ribosomal protein S5 | -1.05 | 4.64E-19 |
| SMU_328 |  | putative carbonic anhydrase | -1.06 | 9.37E-12 |
| SMU_1682c |  | conserved hypothetical protein; possible intracellular protease | -1.06 | 4.26E-08 |
| SMU_533 | trpG | putative anthranilate synthase, beta subunit | -1.06 | 3.95E-08 |
| SMU_363 | glnR | transcriptional regulator; glutamine synthetase repressor | -1.06 | 6.71E-05 |
| SMU_480 | priA | primosomal replication factor Y (primosomal protein N') | -1.06 | 3.59E-13 |
| SMU_1069c |  | hypothetical protein | -1.07 | 4.76E-06 |
| SMU_149 |  | putative transposase | -1.07 | 1.26E-02 |
| SMU_1860 | rs6 | 30S ribosomal protein S6 | -1.07 | 1.61E-17 |
| SMU_1247 | eno | putative enolase | -1.07 | 5.08E-16 |
| SMU_2060 |  | putative transcriptional regulator (LysR family) | -1.08 | 2.71E-10 |
| SMU_364 | glnA | glutamine synthetase type 1; glutamate--ammonia ligase | -1.08 | 1.50E-05 |
| SMU_2104a |  | 50S ribosomal protein L32 | -1.08 | 3.06E-11 |
| SMU_613 |  | hypothetical protein | -1.08 | 8.53E-05 |
| SMU_1859 | ssb | putative single-stranded DNA-binding protein | -1.08 | 3.71E-18 |
| SMU_2116 | opuCa | putative osmoprotectant amino acid ABC transporter, ATP-binding protein | -1.09 | 2.45E-11 |
| SMU_1700c |  | conserved hypothetical protein; possible LrgB family protein | -1.09 | 6.50E-10 |
| SMU_1490 | lacG | 6-phospho-beta-galactosidase | -1.09 | 3.03E-08 |
| SMU_1530 | atpD | FoF1 membrane-bound proton-translocating ATPase, alpha subunit | -1.10 | 3.73E-18 |
| SMU_2008 | rl30 | 50S ribosomal protein L30 | -1.10 | 4.77E-21 |
| SMU_426 | copA | copper-transporting ATPase; P-type ATPase | -1.11 | 5.30E-05 |
| SMU_1055 | radC | putative DNA repair protein RadC | -1.12 | 1.38E-03 |
| SMU_1410 |  | putative reductase | -1.12 | 3.14E-07 |
| SMU_1827 |  | putative biotin biosynthesis protein | -1.12 | 3.21E-08 |
| SMU_649 |  | conserved hypothetical protein | -1.12 | 6.90E-16 |
| SMU_2016 | rl24 | 50S ribosomal protein L24 | -1.13 | 1.62E-22 |
| SMU_1637c |  | hypothetical protein | -1.13 | 5.14E-11 |
| SMU_2032 | rs2 | 30S ribosomal protein S2 | -1.13 | 6.54E-27 |
| SMU_1532 | atpF | FoF1 membrane-bound proton-translocating ATPase, b subunit | -1.14 | 1.36E-12 |
| SMU_1171c |  | conserved hypothetical protein | -1.14 | 2.38E-18 |
| SMU_1196c |  | conserved hypothetical protein | -1.14 | 1.40E-06 |
| SMU_1491 | lacE | PTS system, lactose-specific enzyme IIBC EIIBC-LAC) | -1.14 | 3.71E-08 |
| SMU_1060 | ffh | signal recognition particle protein subunit, Ffh | -1.14 | 8.77E-21 |
| SMU_563 |  | putative ornithine carbamoyltransferase | -1.15 | 1.89E-07 |
| SMU_630 |  | hypothetical protein | -1.15 | 1.04E-10 |
| SMU_1584c |  | putative 67 kDa myosin-crossreactive streptococcal antigen-like protein | -1.15 | 1.12E-14 |
| SMU_850 |  | conserved hypothetical protein | -1.15 | 2.42E-16 |
| SMU_365 | gltA | glutamate synthase (large subunit) | -1.15 | 7.11E-09 |
| SMU_2027 |  | putative transcriptional regulator | -1.15 | 2.58E-12 |
| SMU_1644c |  | hypothetical protein | -1.16 | 1.50E-06 |
| SMU_2118 | opuCc | putative ABC transporter; osmoprotectant-binding protein, glycine betaine/carnitine/choline ABC transporter | -1.16 | 9.63E-16 |
| SMU_139 |  | conserved hypothetical protein | -1.16 | 3.99E-03 |
| SMU_1296 |  | putative glutathione S-transferase | -1.16 | 2.74E-05 |
| SMU_827 | rgpC | putative polysaccharide ABC transporter, permease protein | -1.17 | 2.23E-19 |
| SMU_405c |  | putative transcriptional regulator | -1.17 | 9.09E-04 |
| SMU_1374 |  | hypothetical protein | -1.17 | 2.97E-03 |
| SMU_1817c |  | putative maturase-related protein | -1.17 | 4.70E-09 |
| SMU_1073 | fthS | putative formyl-tetrahydrofolate synthetase | -1.17 | 7.54E-07 |
| SMU_1945 |  | hypothetical protein | -1.17 | 1.34E-10 |
| SMU_104 |  | putative alpha-glucosidase; glycosyl hydrolase | -1.18 | 1.44E-06 |
| SMU_1837 | aroH | putative DAHP synthase; phospho-2-dehydro-3-deoxyphosphoheptonate aldolase | -1.18 | 5.65E-13 |
| SMU_329 |  | conserved hypothetical protein | -1.18 | 4.49E-17 |
| SMU_614 |  | hypothetical protein | -1.18 | 1.70E-10 |
| SMU_2117 | opuCb | putative osmoprotectant ABC transporter; permease protein | -1.19 | 6.42E-15 |
| SMU_1519 | glnQ | putative amino acid ABC transporter, ATP-binding protein | -1.19 | 5.81E-11 |
| SMU_2077c |  | conserved hypothetical protein | -1.19 | 6.76E-08 |
| SMU_419 |  | conserved hypothetical protein | -1.19 | 2.43E-08 |
| SMU_505 |  | putative adenine-specific DNA methylase | -1.19 | 1.69E-04 |
| SMU_1681c |  | conserved hypothetical protein | -1.19 | 8.03E-18 |
| SMU_129 | adhC | putative dihydrolipoamide acetyltransferase | -1.21 | 4.49E-03 |
| SMU_1626 | rl1 | 50S ribosomal protein L1 | -1.21 | 1.50E-31 |
| SMU_360 | gapC | extracellular glyceraldehyde-3-phosphate dehydrogenase | -1.21 | 6.62E-23 |
| SMU_421 |  | translation initiation factor 2 | -1.21 | 1.08E-18 |
| SMU_1003 | gid | putative glucose-inhibited division protein | -1.22 | 1.07E-13 |
| SMU_2119 | opuCd | putative osmoprotectant ABC transporter; permease protein | -1.22 | 3.81E-15 |
| SMU_1528 | atpB | FoF1 membrane-bound proton-translocating ATPase, beta subunit | -1.22 | 5.83E-32 |
| SMU_1237c |  | hypothetical protein | -1.23 | 5.34E-04 |
| SMU_943c |  | putative hydroxymethylglutaryl-CoA synthase | -1.23 | 3.06E-18 |
| SMU_602 |  | putative sodium-dependent transporter | -1.23 | 4.86E-12 |
| SMU_1390 |  | conserved hypothetical protein | -1.24 | 6.33E-18 |
| SMU_1703c |  | conserved hypothetical protein | -1.24 | 1.51E-09 |
| SMU_1007 |  | putative ABC transporter, permease protein | -1.24 | 4.12E-15 |
| SMU_1170 | ccdA | putative cytochrome C biogenesis protein | -1.24 | 7.49E-19 |
| SMU_506 |  | putative type II restriction endonuclease | -1.25 | 2.15E-08 |
| SMU_1531 | atpE | FoF1 membrane-bound proton-translocating ATPase, delta subunit | -1.26 | 1.27E-14 |
| SMU_1809 | scnG | putative bacteroiocin operon protein ScnG-like protein | -1.26 | 8.24E-13 |
| SMU_1805 |  | putative transcriptional regulator | -1.26 | 1.61E-08 |
| SMU_423 |  | hypothetical protein | -1.26 | 4.17E-02 |
| SMU_534 | trpD | putative phosphoribosyl anthranilate transferase | -1.27 | 1.00E-11 |
| SMU_137 | mleS | malolactic enzyme | -1.27 | 4.21E-03 |
| SMU_1203 | ilvE | putative branched-chain amino acid aminotransferase IlvE | -1.27 | 2.23E-19 |
| SMU_2155 |  | conserved hypothetical protein | -1.28 | 2.69E-10 |
| SMU_829 | rgpE | putative glycosyltransferase | -1.28 | 3.65E-30 |
| SMU_2018 | rs17 | 30S ribosomal protein S17 | -1.28 | 9.25E-21 |
| SMU_641 |  | putative oxidoreductase | -1.28 | 1.05E-05 |
| SMU_1117 | naoX | NADH oxidase (H2O-forming) | -1.28 | 3.01E-03 |
| SMU_1815 | scnR | putative response regulator; ScnR-like protein | -1.28 | 4.61E-09 |
| SMU_1680c |  | hypothetical protein | -1.28 | 1.36E-27 |
| SMU_1547c |  | putative response regulator | -1.29 | 1.49E-15 |
| SMU_2038 | pttB | putative PTS system, trehalose-specific IIABC component | -1.29 | 8.72E-09 |
| SMU_184 | sloC | putative ABC transporter, metal binding lipoprotein; surface adhesin precursor; saliva-binding protein; lipoprotein receptor LraI (LraI family) | -1.30 | 2.81E-08 |
| SMU_1648c |  | hypothetical protein | -1.30 | 4.34E-06 |
| SMU_1006 |  | putative ABC transporter, ATP-binding protein | -1.30 | 1.55E-14 |
| SMU_2071 |  | putative anaerobic ribonucleotide reductase activating protein | -1.30 | 2.69E-11 |
| SMU_668c |  | ribonucleotide reductase, large subunit | -1.30 | 3.81E-08 |
| SMU_723 |  | putative calcium-transporting ATPase; P-type ATPase | -1.32 | 2.36E-31 |
| SMU_1201c |  | conserved hypothetical protein | -1.32 | 6.12E-14 |
| SMU_1504c |  | hypothetical protein | -1.32 | 1.29E-06 |
| SMU_825 | rgpA | putative RgpAc; glycosyltransferase | -1.32 | 1.49E-20 |
| SMU_1638c |  | hypothetical protein | -1.33 | 1.59E-14 |
| SMU_1533 | atpG | FoF1 membrane-bound proton-translocating ATPase, a subunit | -1.33 | 1.53E-15 |
| SMU_435 |  | putative N-acetylglucosamine-6-phosphate deacetylase | -1.33 | 4.81E-07 |
| SMU_128 | adhB | putative acetoin dehydrogenase (TPP-dependent), E1 component beta subunit | -1.34 | 5.57E-04 |
| SMU_793 |  | conserved hypothetical protein | -1.34 | 1.79E-16 |
| SMU_1476c |  | putative GTP-binding protein | -1.34 | 2.49E-25 |
| SMU_472 |  | conserved hypothetical protein; possible N6-adenine-specific DNA methylase | -1.35 | 2.81E-18 |
| SMU_1810 | scnE | putative bacteriocin operon component, ScnE-like protein | -1.35 | 3.01E-07 |
| SMU_2115 |  | putative short-chain dehydrogenase | -1.36 | 1.49E-11 |
| SMU_1529 | atpC | FoF1 membrane-bound proton-translocating ATPase, gamma subunit | -1.37 | 4.74E-29 |
| SMU_960 |  | 50S ribosomal protein L7/L12 | -1.37 | 7.06E-19 |
| SMU_451 |  | hypothetical protein | -1.37 | 4.83E-14 |
| SMU_1002 | topA | putative DNA topoisomerase I | -1.38 | 1.89E-13 |
| SMU_532 | trpE | putative anthranilate synthase, alpha subunit | -1.38 | 7.55E-06 |
| SMU_1047c |  | hypothetical protein | -1.39 | 3.78E-13 |
| SMU_2010 | rl18 | 50S ribosomal protein L18 | -1.40 | 6.23E-28 |
| SMU_167 |  | hypothetical protein | -1.40 | 1.13E-08 |
| SMU_697 |  | putative translation initiation factor IF3 | -1.41 | 1.93E-25 |
| SMU_1399 |  | hypothetical protein | -1.42 | 4.27E-04 |
| SMU_102 |  | putative PTS system, IID component | -1.42 | 1.10E-07 |
| SMU_1340 | bacA2 | putative surfactin synthetase | -1.43 | 1.81E-05 |
| SMU_1293c |  | conserved hypothetical protein | -1.43 | 5.68E-25 |
| SMU_1745c |  | putative transcriptional regulator | -1.44 | 4.32E-19 |
| SMU_769 |  | conserved hypothetical protein | -1.44 | 4.88E-05 |
| SMU_2017 | rl14 | 50S ribosomal protein L14 | -1.45 | 6.64E-32 |
| SMU_1505c |  | hypothetical protein | -1.45 | 1.48E-07 |
| SMU_1650 | end3 | putative endonuclease III (DNA repair) | -1.46 | 5.11E-09 |
| SMU_1893c |  | putative transposase, ISSmu1 | -1.46 | 1.63E-04 |
| SMU_2162c |  | conserved hypothetical protein | -1.46 | 4.24E-15 |
| SMU_504 | dam | putative site-specific DNA-methyltransferase | -1.46 | 4.35E-12 |
| SMU_1127 | rs20 | putative 30S ribosomal protein S20 | -1.46 | 2.57E-10 |
| SMU_101 |  | putative sorbose PTS system, IIC component | -1.46 | 6.31E-06 |
| SMU_1701c |  | conserved hypothetical protein | -1.47 | 4.87E-12 |
| SMU_1297 |  | conserved hypothetical protein | -1.47 | 3.35E-06 |
| SMU_424 | copY | negative transcriptional regulator, CopY | -1.47 | 1.13E-07 |
| SMU_629 | sod | putative manganese-type superoxide dismutase, Fe/Mn-SOD | -1.47 | 3.84E-05 |
| SMU_1908c |  | hypothetical protein | -1.48 | 2.88E-02 |
| SMU_1902c |  | hypothetical protein | -1.48 | 1.09E-03 |
| SMU_2014 | rs14 | 30S ribosomal protein S14 | -1.48 | 1.35E-32 |
| SMU_1788c |  | putative bacterocin transport accessory protein, Bta | -1.49 | 5.53E-07 |
| SMU_747c |  | conserved hypothetical protein; putative permease | -1.50 | 1.51E-16 |
| SMU_1070c |  | conserved hypothetical protein | -1.50 | 5.80E-12 |
| SMU_1548c |  | putative histidine kinase | -1.51 | 6.12E-31 |
| SMU_738 |  | hypothetical protein | -1.51 | 1.67E-08 |
| SMU_2104 |  | conserved hypothetical protein; possible integral membrane protein | -1.51 | 1.75E-35 |
| SMU_1808c |  | putative integrase fragment | -1.51 | 5.30E-07 |
| SMU_1692 | pflA | pyruvate-formate lyase activating enzyme | -1.51 | 7.26E-10 |
| SMU_562 | clpE | ATP-dependent protease ClpE | -1.52 | 1.79E-25 |
| SMU_138 |  | putative malate permease | -1.52 | 4.95E-04 |
| SMU_944 | thyA | thymidylate synthase | -1.52 | 4.03E-15 |
| SMU_2078c |  | conserved hypothetical protein | -1.53 | 3.41E-15 |
| SMU_1061 | ylxM | putative DNA-binding protein | -1.53 | 5.26E-28 |
| SMU_2111c |  | hypothetical protein | -1.53 | 6.65E-12 |
| SMU_80 | hrcA | transcriptional regulator; repressor (HrcA) of class I heat shock genes | -1.54 | 9.15E-08 |
| SMU_1534 | atpH | FoF1 membrane-bound proton-translocating ATPase, c subunit | -1.54 | 3.98E-15 |
| SMU_1657c |  | putative nitrogen regulatory protein PII | -1.55 | 7.59E-03 |
| SMU_524 |  | putative ABC transporter, ATP-binding protein | -1.55 | 5.80E-05 |
| SMU_2003a |  | 50S ribosomal protein L36 | -1.56 | 1.57E-11 |
| SMU_754 |  | HPr(serine) kinase/phosphatase | -1.56 | 8.31E-36 |
| SMU_1679c |  | conserved hypothetical protein | -1.58 | 1.07E-29 |
| SMU_462 |  | conserved hypothetical protein | -1.58 | 1.35E-14 |
| SMU_260 |  | conserved hypothetical protein | -1.58 | 4.93E-08 |
| SMU_152 |  | hypothetical protein | -1.59 | 1.08E-02 |
| SMU_836 |  | hypothetical protein | -1.61 | 5.80E-04 |
| SMU_1861c |  | hypothetical protein | -1.61 | 1.34E-07 |
| SMU_204c |  | hypothetical protein | -1.61 | 2.93E-03 |
| SMU_525 |  | putative ABC transporter, ATP-binding protein | -1.62 | 1.79E-10 |
| SMU_1195 |  | conserved hypothetical protein; possible permease | -1.62 | 3.22E-25 |
| SMU_1495 | lacB | galactose-6-phosphate isomerase, subunit LacB | -1.62 | 3.03E-05 |
| SMU_794 |  | hypothetical protein | -1.63 | 1.74E-24 |
| SMU_1585c |  | putative transcriptional regulator | -1.63 | 6.22E-21 |
| SMU_2139c |  | 50S ribosomal protein L9 | -1.63 | 3.07E-30 |
| SMU_840c |  | hypothetical protein | -1.63 | 2.55E-13 |
| SMU_755 |  | putative prolipoprotein diacylglycerol transferase | -1.63 | 2.46E-32 |
| SMU_529 |  | hypothetical protein | -1.63 | 4.63E-09 |
| SMU_1595 | cah | putative carbonic anhydrase precursor | -1.64 | 2.38E-20 |
| SMU_636 |  | putative N-acetylglucosamine-6-phosphate isomerase | -1.64 | 9.98E-14 |
| SMU_796 |  | conserved hypothetical protein | -1.64 | 7.09E-30 |
| SMU_1284c |  | conserved hypothetical protein | -1.64 | 1.29E-10 |
| SMU_2079c |  | conserved hypothetical protein | -1.65 | 2.65E-17 |
| SMU_63c |  | conserved hypothetical protein | -1.66 | 2.28E-24 |
| SMU_890 |  | conserved hypothetical protein | -1.66 | 6.46E-31 |
| SMU_2015 | rl5 | 50S ribosomal protein L5 | -1.66 | 1.85E-37 |
| SMU_797 |  | conserved hypothetical protein | -1.67 | 6.35E-21 |
| SMU_1658 | nrgA | putative ammonium transporter, NrgA protein | -1.67 | 3.87E-03 |
| SMU_1999c |  | conserved hypothetical protein | -1.67 | 2.24E-27 |
| SMU_2166 |  | 50S Ribosomal Protein L23 | -1.69 | 8.35E-46 |
| SMU_2161c |  | conserved hypothetical protein | -1.69 | 1.98E-26 |
| SMU_565c |  | putative transposase, ISSmu1 | -1.69 | 6.32E-04 |
| SMU_166 |  | hypothetical protein | -1.71 | 1.26E-14 |
| SMU_335 |  | argininosuccinate lyase | -1.72 | 4.02E-19 |
| SMU_962 |  | putative dehydrogenase | -1.73 | 1.56E-21 |
| SMU_2020 | rl16 | 50S ribosomal protein L16 | -1.73 | 5.51E-41 |
| SMU_196c |  | putative transfer protein | -1.73 | 1.24E-03 |
| SMU_1627 | rl11 | 50S ribosomal L11 protein | -1.74 | 1.14E-43 |
| SMU_795 |  | conserved hypothetical protein; probable esterase | -1.74 | 1.91E-30 |
| SMU_109 |  | conserved hypothetical protein; possible permease (efflux protein) | -1.76 | 1.45E-18 |
| SMU_1869 | trxA | putative thioredoxin | -1.76 | 3.28E-09 |
| SMU_1185 | mtlA1 | PTS system, mannitol-specific enzyme IIBC component | -1.76 | 2.56E-06 |
| SMU_1341c |  | putative gramicidin S synthetase | -1.76 | 1.90E-07 |
| SMU_951 |  | putative amino acid permease | -1.76 | 5.70E-28 |
| SMU_1342 | bacA1 | putative bacitracin synthetase 1; BacA | -1.76 | 9.37E-07 |
| SMU_2021 | rs3 | 30S ribosomal protein S3 | -1.76 | 7.46E-48 |
| SMU_1910c |  | hypothetical protein | -1.77 | 1.17E-02 |
| SMU_531 |  | putative chorismate mutase | -1.77 | 6.64E-20 |
| SMU_144c |  | putative transcriptional regulator | -1.78 | 4.74E-18 |
| SMU_418 | nusA | putative transcription factor NusA | -1.79 | 7.40E-42 |
| SMU_1750c |  | hypothetical protein | -1.80 | 1.46E-17 |
| SMU_1912c |  | hypothetical protein | -1.81 | 6.71E-03 |
| SMU_1493 | lacD | tagatose-1,6-bisphosphate aldolase | -1.82 | 6.20E-10 |
| SMU_1419 |  | putative transcriptional regulator | -1.82 | 5.85E-08 |
| SMU_2022 | rl22 | 50S ribosomal protein L22 | -1.82 | 3.13E-39 |
| SMU_2084c |  | conserved hypothetical protein | -1.82 | 7.56E-22 |
| SMU_1904c |  | hypothetical protein | -1.83 | 2.82E-03 |
| SMU_133c |  | putative MDR permease; transmembrane efflux protein | -1.84 | 6.65E-17 |
| SMU_1193 |  | putative transcriptional regulator | -1.85 | 4.97E-24 |
| SMU_2023c |  | 30S ribosomal protein S19 | -1.85 | 1.61E-46 |
| SMU_644 |  | putative competence protein/transcription factor | -1.86 | 5.29E-03 |
| SMU_463 | trxB | putative thioredoxin reductase (NADPH) | -1.86 | 1.87E-12 |
| SMU_540 | dpr | peroxide resistance protein Dpr | -1.88 | 5.42E-18 |
| SMU_539c |  | signal peptidase type IV | -1.88 | 3.26E-04 |
| SMU_2138 | dnaC | putative replicative DNA helicase (DNA polymerase III delta prime subunit) | -1.89 | 4.22E-39 |
| SMU_673 |  | conserved hypothetical protein | -1.90 | 1.02E-03 |
| SMU_417 |  | conserved hypothetical protein | -1.90 | 6.49E-39 |
| SMU_1905c |  | putative bacteriocin secretion protein | -1.91 | 2.28E-03 |
| SMU_1420 |  | putative oxidoreductase | -1.91 | 3.10E-08 |
| SMU_2109 |  | putative MDR permease; possible multidrug efflux pump | -1.91 | 5.68E-25 |
| SMU_1194 |  | putative ABC transporter, ATP-binding protein | -1.91 | 1.42E-32 |
| SMU_611 |  | putative ATP-dependent RNA helicase, DEAD-box family | -1.92 | 5.76E-46 |
| SMU_510c |  | hypothetical protein | -1.93 | 5.87E-15 |
| SMU_1286c |  | putative permease; multidrug efflux protein | -1.93 | 6.46E-31 |
| SMU_208c |  | putative transposon protein; possible DNA segregation ATPase | -1.94 | 3.92E-05 |
| SMU_753 |  | conserved hypothetical protein | -1.95 | 8.75E-15 |
| SMU_1909c |  | hypothetical protein | -1.96 | 5.84E-03 |
| SMU_1345c |  | putative peptide synthetase | -1.98 | 6.27E-08 |
| SMU_2025 | rl3 | 50S ribosomal protein L3 | -1.99 | 1.38E-83 |
| SMU_202c |  | hypothetical protein | -1.99 | 1.98E-04 |
| SMU_2140c |  | conserved hypothetical protein | -1.99 | 1.45E-55 |
| SMU_2167 |  | 50S Ribosomal Protein L2 | -1.99 | 1.42E-80 |
| SMU_2141 | gidA | glucose inhibited division protein-like protein GidA | -2.00 | 2.38E-39 |
| SMU_737 |  | conserved hypothetical protein | -2.00 | 4.51E-22 |
| SMU_1913c |  | putative immunity protein, BLpL-like | -2.01 | 1.90E-03 |
| SMU_1397c |  | conserved hypothetical protein | -2.02 | 6.89E-23 |
| SMU_1346 | bacT | putative thioesterase BacT | -2.04 | 2.07E-10 |
| SMU_940c |  | putative hemolysin III | -2.05 | 7.06E-22 |
| SMU_1867c |  | putative alcohol dehydrogenase | -2.06 | 9.49E-16 |
| SMU_195c |  | hypothetical protein | -2.06 | 2.04E-05 |
| SMU_2003 | rs13 | 30S ribosomal protein S13 | -2.08 | 2.94E-71 |
| SMU_1903c |  | hypothetical protein | -2.10 | 6.20E-04 |
| SMU_1343c |  | putative polyketide synthase | -2.10 | 1.32E-08 |
| SMU_1488c |  | conserved hypothetical protein | -2.11 | 1.27E-13 |
| SMU_1236c |  | conserved hypothetical protein | -2.11 | 1.09E-20 |
| SMU_1494 | lacC | tagatose-6-phosphate kinase | -2.12 | 2.90E-13 |
| SMU_200c |  | hypothetical protein | -2.13 | 9.08E-04 |
| SMU_1344c |  | putative malonyl-CoA acyl-carrier-protein transacylase | -2.13 | 6.03E-09 |
| SMU_924 | tpx | thiol peroxidase | -2.14 | 5.10E-10 |
| SMU_2005 | adk | putative adenylate kinase | -2.14 | 3.24E-53 |
| SMU_961 |  | conserved hypothetical protein | -2.16 | 1.37E-24 |
| SMU_207c |  | putative transposon protein | -2.16 | 4.72E-06 |
| SMU_669c |  | putative glutaredoxin | -2.19 | 1.07E-14 |
| SMU_2057c |  | putative cadmium-transporting ATPase; P-type ATPase | -2.20 | 1.15E-14 |
| SMU_369c |  | conserved hypothetical protein | -2.21 | 5.98E-20 |
| SMU_2004 | if1 | putative translation initiation factor IF-1 | -2.21 | 3.10E-68 |
| SMU_1421 | pdhC | putative dihydrolipoamide acetyltransferase, E2 component | -2.23 | 7.36E-03 |
| SMU_2024c |  | 50S ribosomal protein L4 | -2.25 | 1.71E-79 |
| SMU_168 |  | putative transcriptional regulator | -2.25 | 8.81E-24 |
| SMU_1496 | lacA | galactose-6-phosphate isomerase, subunit LacA | -2.26 | 7.31E-08 |
| SMU_767 |  | putative transposase, ISSmu1 | -2.26 | 9.88E-08 |
| SMU_2083c |  | hypothetical protein | -2.26 | 1.98E-28 |
| SMU_527 |  | conserved hypothetical protein | -2.27 | 6.39E-21 |
| SMU_1489 | lacX | conserved hypothetical protein, LacX | -2.29 | 9.25E-21 |
| SMU_209c |  | hypothetical protein | -2.29 | 4.86E-06 |
| SMU_1367c |  | conserved hypothetical protein | -2.30 | 2.88E-28 |
| SMU_2001 | rpoA | DNA-directed RNA polymerase, alpha subunit | -2.32 | 1.19E-119 |
| SMU_1093 |  | putative ABC transporter, permease protein | -2.32 | 9.86E-18 |
| SMU_725c |  | conserved hypothetical protein | -2.33 | 1.19E-30 |
| SMU_1423 | pdhA | putative pyruvate dehydrogenase, TPP-dependent E1 component alpha-subunit | -2.36 | 3.69E-03 |
| SMU_1094 |  | putative ABC transporter, ATP-binding protein | -2.36 | 2.04E-09 |
| SMU_183 | sloB | putative Mn/Zn ABC transporter | -2.36 | 1.44E-13 |
| SMU_1577c |  | conserved hypothetical protein | -2.37 | 5.68E-25 |
| SMU_1034c |  | putative integrase/recombinase; XerC-like | -2.38 | 5.49E-18 |
| SMU_2002 | rs11 | 30S ribosomal protein S11 | -2.39 | 2.61E-73 |
| SMU_205c |  | hypothetical protein | -2.41 | 1.03E-05 |
| SMU_1550c |  | conserved hypothetical protein; possible integral membrane protein | -2.44 | 2.16E-31 |
| SMU_1422 | pdhB | putative pyruvate dehydrogenase E1 component beta subunit) | -2.44 | 2.72E-03 |
| SMU_100 |  | putative sorbose PTS system, IIB component | -2.45 | 6.01E-10 |
| SMU_941c |  | conserved hypothetical protein | -2.46 | 1.07E-32 |
| SMU_191c |  | putative integrase | -2.48 | 7.26E-06 |
| SMU_1424 | pdhD | putative dihydrolipoamide dehydrogenase | -2.49 | 1.09E-04 |
| SMU_197c |  | hypothetical protein | -2.50 | 6.04E-07 |
| SMU_2028 | sacB | levansucrase precursor; beta-D-fructosyltransferase | -2.52 | 1.27E-57 |
| SMU_16 |  | putative amino acid permease | -2.53 | 2.22E-52 |
| SMU_2026c |  | 30S ribosomal protein S10 | -2.53 | 9.49E-85 |
| SMU_1707c |  | putative rRNA methylase | -2.56 | 5.10E-27 |
| SMU_1552c |  | hypothetical protein | -2.57 | 3.07E-15 |
| SMU_1997 | comX1 | putative ComX1, transcriptional regulator of competence-specific genes | -2.59 | 4.44E-31 |
| SMU_1030 |  | putative polyribonucleotide nucleotidyltransferase; Tn916 ORF8-like | -2.62 | 3.11E-02 |
| SMU_2000 | rl17 | 50S ribosomal protein L17 | -2.63 | 9.64E-100 |
| SMU_210c |  | hypothetical protein | -2.68 | 2.11E-07 |
| SMU_198c |  | putative conjugative transposon protein | -2.68 | 1.31E-07 |
| SMU_1551c |  | putative ABC transporter, ATP-binding protein | -2.71 | 9.49E-27 |
| SMU_639 |  | putative acetyltransferase | -2.76 | 4.49E-21 |
| SMU_1554c |  | hypothetical protein | -2.78 | 1.66E-11 |
| SMU_1438c |  | putative Zn-dependent protease | -2.78 | 4.18E-51 |
| SMU_957 |  | 50S ribosomal protein L10 | -2.79 | 1.36E-68 |
| SMU_956 | clp | putative Clp-like ATP-dependent protease, ATP-binding subunit | -2.81 | 2.15E-18 |
| SMU_201c |  | putative transposon protein | -2.87 | 2.16E-08 |
| SMU_106c |  | putative transposase fragment | -2.91 | 9.55E-48 |
| SMU_1553c |  | hypothetical protein | -2.92 | 1.14E-20 |
| SMU_1856c |  | conserved hypothetical protein | -2.92 | 1.13E-82 |
| SMU_199c |  | hypothetical protein | -3.00 | 6.72E-09 |
| SMU_1131c |  | hypothetical protein | -3.03 | 1.10E-25 |
| SMU_182 | sloA | putative ABC transporter, ATP-binding protein; possible iron and/or manganese ABC transport system | -3.14 | 1.76E-12 |
| SMU_638 |  | putative 16S pseudouridylate synthase | -3.18 | 2.36E-30 |
| SMU_1894c |  | conserved hypothetical protein | -3.20 | 6.30E-08 |
| SMU_766 |  | conserved hypothetical protein | -3.21 | 8.69E-07 |
| SMU_40 |  | conserved hypothetical protein | -3.22 | 4.23E-10 |
| SMU_672 | idh | isocitrate dehydrogenase | -3.24 | 9.58E-07 |
| SMU_671 | citZ | citrate synthase | -3.53 | 1.36E-07 |
| SMU_1184c |  | putative transcriptional regulator, antiterminator | -3.69 | 1.98E-20 |
| SMU_670 | citB | aconitate hydratase; aconitase | -4.02 | 4.42E-10 |
| SMU_1408c |  | conserved hypothetical protein | -4.18 | 3.28E-09 |
| SMU_575c |  | putative membrane protein | -4.42 | 2.03E-05 |
| SMU_574c |  | putative membrane protein | -4.57 | 1.20E-04 |
| SMU_1365c |  | hypothetical protein; possible permease | -4.64 | 3.10E-52 |
| SMU_368c | rnjA | conserved hypothetical protein | -4.77 | 2.05E-165 |
| SMU_1348c |  | putative ABC transporter, ATP-binding protein | -5.25 | 3.96E-54 |
| SMU_1347c |  | conserved hypothetical protein; possible permease | -6.15 | 8.47E-42 |
| SMU_1366c |  | putative ABC transporter; ATP-binding protein | -8.42 | 3.15E-83 |

**Table S3 Differential gene expression (RNA-seq) analysis of *S. mutans* UA159 during CRISPRi-mediated knockdown of SMU_369c**

| **Locus tag** | **Gene** | **Description** | **log2 fold change** | **FDR** |
| --- | --- | --- | --- | --- |
| SMU_1405c | cas9 | conserved hypothetical protein | 7.85 | 2.37E-125 |
| SMU_2037 | treA | putative trehalose-6-phosphate hydrolase TreA | 1.40 | 1.79E-04 |
| SMU_1048 |  | conserved hypothetical protein | 1.09 | 4.44E-02 |
| SMU_1757c |  | conserved hypothetical protein | 1.04 | 5.27E-03 |
| SMU_1780 |  | conserved hypothetical protein | 1.02 | 4.44E-02 |
| SMU_1420 |  | putative oxidoreductase | -1.02 | 1.13E-02 |
| SMU_101 |  | putative sorbose PTS system, IIC component | -1.10 | 2.71E-03 |
| SMU_1928 | psaB | putative ABC transporter, permease protein | -1.26 | 1.83E-03 |
| SMU_1004 | gtfB | glucosyltransferase-I | -1.42 | 3.08E-06 |

**Table S4 Differential gene expression (RNA-seq) analysis of *S. mutans* UA159 during CRISPRi-mediated knockdown of SMU_393**

| **Locus tag** | **Gene** | **Description** | **log2 fold change** | **FDR** |
| --- | --- | --- | --- | --- |
| SMU_1405c | cas9 | conserved hypothetical protein | 7.48 | 4.62E-155 |
| SMU_1155 |  | hypothetical protein | 2.42 | 1.92E-04 |
| SMU_193c |  | conserved hypothetical protein | 2.32 | 4.88E-02 |
| SMU_215c |  | hypothetical protein | 2.19 | 5.22E-03 |
| SMU_687c |  | hypothetical protein | 1.87 | 2.47E-02 |
| SMU_196c |  | putative transfer protein | 1.86 | 5.22E-03 |
| SMU_213c |  | hypothetical protein | 1.73 | 2.47E-02 |
| SMU_211c |  | hypothetical protein | 1.70 | 1.37E-02 |
| SMU_207c |  | putative transposon protein | 1.66 | 4.12E-03 |
| SMU_202c |  | hypothetical protein | 1.66 | 1.20E-02 |
| SMU_200c |  | hypothetical protein | 1.65 | 4.15E-02 |
| SMU_191c |  | putative integrase | 1.62 | 1.89E-02 |
| SMU_198c |  | putative conjugative transposon protein | 1.55 | 1.21E-02 |
| SMU_208c |  | putative transposon protein; possible DNA segregation ATPase | 1.55 | 8.46E-03 |
| SMU_197c |  | hypothetical protein | 1.54 | 1.22E-02 |
| SMU_201c |  | putative transposon protein | 1.50 | 1.58E-02 |
| SMU_206c |  | hypothetical protein | 1.41 | 2.67E-02 |
| SMU_209c |  | hypothetical protein | 1.34 | 3.42E-02 |
| SMU_1150 |  | putative transporter, trans-membrane domain bacteriocin immunity protein | 1.30 | 4.73E-02 |
| SMU_210c |  | hypothetical protein | 1.27 | 4.93E-02 |
| SMU_641 |  | putative oxidoreductase | 1.24 | 3.70E-04 |
| SMU_1148 |  | putative transporter, ATP-binding protein; bacteriocin immunity protein | 1.19 | 1.21E-02 |
| SMU_1378 |  | hypothetical protein | 1.19 | 2.16E-02 |
| SMU_643 |  | putative esterase | 1.18 | 7.21E-05 |
| SMU_217c |  | hypothetical protein | 1.10 | 5.59E-02 |
| SMU_1747c |  | putative phosphatase | 1.07 | 3.89E-06 |
| SMU_642 |  | hypothetical protein | 1.06 | 5.52E-03 |
| SMU_772 | gbpD | glucan-binding protein D with lipase activity; BglB-like protein | 1.05 | 6.36E-07 |
| SMU_1862 |  | hypothetical protein | 1.03 | 1.77E-03 |
| SMU_1005 | gtfC | glucosyltransferase-SI | -1.01 | 3.41E-23 |
| SMU_812 |  | hypothetical protein | -1.01 | 2.06E-03 |
| SMU_555 | ylmG | conserved hypothetical protein | -1.02 | 5.56E-04 |
| SMU_405c |  | putative transcriptional regulator | -1.02 | 2.23E-02 |
| SMU_366 | gltB | NADPH-dependent glutamate synthase (small subunit) | -1.03 | 8.61E-06 |
| SMU_103 |  | putative PTS system, IIA component | -1.13 | 4.13E-04 |
| SMU_1861c |  | hypothetical protein | -1.13 | 2.60E-03 |
| SMU_1927 |  | putative ABC transporter, ATP-binding protein | -1.13 | 2.92E-02 |
| SMU_1803c |  | hypothetical protein | -1.14 | 7.31E-07 |
| SMU_239c |  | hypothetical protein | -1.15 | 1.73E-04 |
| SMU_223c |  | hypothetical protein | -1.22 | 4.81E-04 |
| SMU_984 |  | hypothetical protein | -1.25 | 2.86E-09 |
| SMU_1928 | psaB | putative ABC transporter, permease protein | -1.26 | 1.20E-02 |
| SMU_102 |  | putative PTS system, IID component | -1.26 | 7.21E-05 |
| SMU_1410 |  | putative reductase | -1.29 | 3.12E-07 |
| SMU_104 |  | putative alpha-glucosidase; glycosyl hydrolase | -1.34 | 1.69E-06 |
| SMU_1187 | glmS | glucosamine-fructose-6-phosphate aminotransferase | -1.37 | 4.78E-04 |
| SMU_101 |  | putative sorbose PTS system, IIC component | -1.38 | 3.70E-04 |
| SMU_457 |  | hypothetical protein | -1.38 | 4.00E-03 |
| SMU_1395c |  | hypothetical protein | -1.52 | 1.11E-04 |
| SMU_1575c |  | hypothetical protein | -1.55 | 4.65E-03 |
| SMU_510c |  | hypothetical protein | -1.71 | 2.60E-10 |
| SMU_1658 | nrgA | putative ammonium transporter, NrgA protein | -1.71 | 1.78E-02 |
| SMU_545 |  | hypothetical protein | -1.72 | 1.80E-03 |
| SMU_1804c |  | hypothetical protein | -1.76 | 9.39E-03 |
| SMU_1657c |  | putative nitrogen regulatory protein PII | -1.80 | 1.27E-02 |
| SMU_609 |  | putative 40K cell wall protein precursor | -1.82 | 2.28E-04 |
| SMU_100 |  | putative sorbose PTS system, IIB component | -1.85 | 5.12E-05 |
| SMU_1004 | gtfB | glucosyltransferase-I | -1.95 | 1.34E-13 |
| SMU_670 | citB | aconitate hydratase; aconitase | -2.11 | 4.86E-03 |
| SMU_750c |  | hypothetical protein | -2.16 | 3.95E-03 |
| SMU_503c |  | hypothetical protein | -2.32 | 2.05E-19 |
| SMU_671 | citZ | citrate synthase | -2.52 | 1.44E-03 |
| SMU_673 |  | conserved hypothetical protein | -2.59 | 2.03E-04 |
| SMU_672 | idh | isocitrate dehydrogenase | -2.92 | 1.86E-04 |
| SMU_393 |  | conserved hypothetical protein | -2.94 | 1.51E-37 |

**Table S5 Differential gene expression (RNA-seq) analysis of S. mutans UA159 during CRISPRi-mediated knockdown of SMU_415**

| **Locus tag** | **Gene** | **Description** | **log2 fold change** | **FDR** |
| --- | --- | --- | --- | --- |
| SMU_958 |  | hypothetical protein | 7.79 | 4.24E-42 |
| SMU_1405c | cas9 | conserved hypothetical protein | 7.18 | 9.25E-147 |
| SMU_2053c |  | hypothetical protein | 3.94 | 6.66E-06 |
| SMU_150 |  | hypothetical protein | 3.44 | 4.03E-06 |
| SMU_1914c |  | hypothetical protein | 3.15 | 5.27E-06 |
| SMU_574c |  | putative membrane protein | 2.84 | 2.77E-02 |
| SMU_193c |  | conserved hypothetical protein | 2.70 | 3.22E-02 |
| SMU_114 |  | putative PTS system, fructose-specific IIBC component | 2.65 | 2.11E-04 |
| SMU_1980c |  | conserved hypothetical protein | 2.59 | 3.70E-02 |
| SMU_206c |  | hypothetical protein | 2.56 | 1.27E-05 |
| SMU_1895c |  | hypothetical protein | 2.51 | 1.26E-13 |
| SMU_116 | lacD2 | tagatose 1,6-aldolase | 2.46 | 3.32E-04 |
| SMU_1906c |  | hypothetical protein | 2.44 | 6.24E-03 |
| SMU_153 |  | hypothetical protein | 2.35 | 4.24E-04 |
| SMU_215c |  | hypothetical protein | 2.31 | 1.27E-03 |
| SMU_1896c |  | hypothetical protein | 2.28 | 6.62E-13 |
| SMU_213c |  | hypothetical protein | 2.27 | 1.01E-03 |
| SMU_115 |  | putative PTS system, fructose-specific IIA component | 2.27 | 8.30E-03 |
| SMU_151 |  | hypothetical protein | 2.17 | 5.02E-03 |
| SMU_1423 | pdhA | putative pyruvate dehydrogenase, TPP-dependent E1 component alpha-subunit | 2.13 | 2.27E-02 |
| SMU_1597c |  | conserved hypothetical protein | 2.12 | 1.32E-02 |
| SMU_1359 |  | hypothetical protein | 2.06 | 3.53E-02 |
| SMU_1421 | pdhC | putative dihydrolipoamide acetyltransferase, E2 component | 2.02 | 3.60E-02 |
| SMU_212c |  | hypothetical protein | 1.99 | 3.32E-03 |
| SMU_1774c |  | conserved hypothetical protein | 1.94 | 4.12E-14 |
| SMU_113 |  | putative fructose-1-phosphate kinase | 1.90 | 4.13E-03 |
| SMU_1908c |  | hypothetical protein | 1.90 | 1.54E-02 |
| SMU_216c |  | hypothetical protein | 1.89 | 1.08E-02 |
| SMU_1600 | ptcB | putative PTS system, cellobiose-specific IIB component | 1.89 | 2.29E-03 |
| SMU_934 |  | putative amino acid ABC transporter, permease protein | 1.89 | 7.55E-04 |
| SMU_626 |  | putative competence protein | 1.87 | 1.16E-01 |
| SMU_1422 | pdhB | putative pyruvate dehydrogenase E1 component beta subunit) | 1.87 | 4.48E-02 |
| SMU_1912c |  | hypothetical protein | 1.87 | 1.50E-02 |
| SMU_217c |  | hypothetical protein | 1.85 | 3.39E-04 |
| SMU_1910c |  | hypothetical protein | 1.81 | 2.66E-02 |
| SMU_423 |  | hypothetical protein | 1.80 | 1.12E-02 |
| SMU_933 |  | putative amino acid ABC transporter, periplasmic amino acid-binding protein | 1.80 | 3.39E-04 |
| SMU_1904c |  | hypothetical protein | 1.80 | 1.03E-02 |
| SMU_1979c |  | conserved hypothetical protein | 1.78 | 1.97E-02 |
| SMU_1905c |  | putative bacteriocin secretion protein | 1.76 | 1.44E-02 |
| SMU_1913c |  | putative immunity protein, BLpL-like | 1.74 | 1.96E-02 |
| SMU_1752c |  | hypothetical protein | 1.69 | 9.89E-11 |
| SMU_935 |  | putative amino acid ABC transporter, permease protein | 1.69 | 4.50E-02 |
| SMU_262 | otcA | putative ornithine carbamoyltransferase | 1.69 | 1.64E-05 |
| SMU_11 |  | conserved hypothetical protein | 1.67 | 1.40E-02 |
| SMU_1260c |  | conserved hypothetical protein | 1.64 | 5.62E-11 |
| SMU_1902c |  | hypothetical protein | 1.63 | 1.33E-03 |
| SMU_202c |  | hypothetical protein | 1.63 | 6.61E-03 |
| SMU_194c |  | conserved hypothetical protein; Bacteriophage P2 associated | 1.62 | 1.91E-02 |
| SMU_27 | acpP | putative acyl carrier protein; AcpP; ACP | 1.61 | 1.33E-07 |
| SMU_108 |  | hypothetical protein | 1.57 | 1.35E-04 |
| SMU_124 |  | putative transcriptional regulator (MarR family) | 1.50 | 1.19E-06 |
| SMU_616 |  | hypothetical protein | 1.50 | 2.91E-04 |
| SMU_2019 | rl29 | 50S ribosomal protein L29 | 1.49 | 1.86E-14 |
| SMU_1641c |  | conserved hypothetical protein | 1.47 | 2.65E-06 |
| SMU_204c |  | hypothetical protein | 1.43 | 1.82E-02 |
| SMU_260 |  | conserved hypothetical protein | 1.42 | 6.53E-06 |
| SMU_2129c |  | conserved hypothetical protein | 1.36 | 6.63E-08 |
| SMU_125 |  | conserved hypothetical protein | 1.36 | 4.70E-06 |
| SMU_1604c |  | conserved hypothetical protein | 1.34 | 3.72E-06 |
| SMU_735 |  | hypothetical protein | 1.33 | 3.44E-06 |
| SMU_1299c |  | putative acetate kinase | 1.33 | 2.83E-10 |
| SMU_947 | dfrA | putative dihydrofolate reductase | 1.32 | 1.92E-12 |
| SMU_936 |  | putative amino acid ABC transporter, ATP-binding protein | 1.31 | 2.58E-02 |
| SMU_1882c |  | hypothetical protein | 1.31 | 1.47E-07 |
| SMU_369c |  | conserved hypothetical protein | 1.29 | 2.80E-07 |
| SMU_930c |  | putative transcriptional regulator | 1.28 | 2.23E-05 |
| SMU_1977c |  | putative transcriptional regulator | 1.28 | 8.10E-16 |
| SMU_195c |  | hypothetical protein | 1.28 | 1.79E-02 |
| SMU_299c |  | putative bacteriocin peptide precursor | 1.26 | 3.63E-05 |
| SMU_1370c |  | putative transposase, IS150-like | 1.22 | 1.49E-02 |
| SMU_993 |  | putative GTP-binding protein | 1.22 | 1.18E-16 |
| SMU_214c |  | hypothetical protein | 1.22 | 1.40E-02 |
| SMU_265 | arcC | putative carbamate kinase | 1.20 | 1.13E-03 |
| SMU_1298 | rl31 | 50S ribosomal protein L31 | 1.20 | 5.27E-06 |
| SMU_932 |  | hypothetical protein | 1.19 | 3.39E-02 |
| SMU_393 |  | conserved hypothetical protein | 1.19 | 8.91E-08 |
| SMU_277 |  | hypothetical protein | 1.18 | 6.63E-09 |
| SMU_218 |  | putative transcriptional regulator | 1.18 | 3.95E-06 |
| SMU_1479 |  | conserved hypothetical protein | 1.17 | 1.15E-05 |
| SMU_420 |  | putative ribosomal protein | 1.16 | 8.62E-05 |
| SMU_1064c |  | putative transcriptional regulator (GntR family) | 1.16 | 1.72E-06 |
| SMU_81 | grpE | heat shock protein GrpE (HSP-70 cofactor) | 1.16 | 5.58E-04 |
| SMU_1978 | ackA | putative acetate kinase | 1.15 | 1.84E-03 |
| SMU_711 |  | conserved hypothetical protein | 1.11 | 6.06E-07 |
| SMU_2137c |  | conserved hypothetical protein | 1.11 | 3.37E-08 |
| SMU_1972c |  | conserved hypothetical protein | 1.11 | 1.03E-06 |
| SMU_837 |  | putative reductase | 1.08 | 7.09E-03 |
| SMU_896 |  | conserved hypothetical protein | 1.08 | 3.27E-03 |
| SMU_849 |  | 50S ribosomal protein L27 | 1.08 | 4.31E-10 |
| SMU_730 |  | conserved hypothetical protein | 1.08 | 9.28E-03 |
| SMU_1300c |  | conserved hypothetical protein | 1.07 | 9.47E-10 |
| SMU_2136c |  | hypothetical protein | 1.07 | 7.20E-05 |
| SMU_1502c |  | conserved hypothetical protein | 1.06 | 1.38E-07 |
| SMU_427 | copZ | putative copper chaperone | 1.06 | 3.09E-03 |
| SMU_470 |  | conserved hypothetical protein | 1.06 | 1.51E-05 |
| SMU_999 |  | hypothetical protein | 1.03 | 1.74E-03 |
| SMU_68 |  | hypothetical protein | 1.03 | 4.41E-08 |
| SMU_772 | gbpD | glucan-binding protein D with lipase activity; BglB-like protein | 1.02 | 2.88E-07 |
| SMU_1256c |  | hypothetical protein | 1.02 | 6.83E-06 |
| SMU_1862 |  | hypothetical protein | 1.02 | 4.27E-04 |
| SMU_1065c |  | putative transcriptional regulator (GntR family) | 1.01 | 1.36E-07 |
| SMU_1775c |  | hypothetical protein | 1.00 | 8.92E-05 |
| SMU_89c |  | putative nitrite transporter | -1.00 | 1.49E-04 |
| SMU_40 |  | conserved hypothetical protein | -1.03 | 6.45E-02 |
| SMU_1396 | gbpC | glucan-binding protein C, GbpC | -1.09 | 2.94E-05 |
| SMU_510c |  | hypothetical protein | -1.09 | 5.91E-05 |
| SMU_1395c |  | hypothetical protein | -1.10 | 2.80E-03 |
| SMU_956 | clp | putative Clp-like ATP-dependent protease, ATP-binding subunit | -1.19 | 5.93E-04 |
| SMU_503c |  | hypothetical protein | -1.23 | 3.27E-06 |
| SMU_405c |  | putative transcriptional regulator | -1.29 | 1.17E-03 |
| SMU_404c |  | hypothetical protein | -1.35 | 6.38E-05 |
| SMU_1004 | gtfB | glucosyltransferase-I | -1.56 | 3.25E-09 |
| SMU_671 | citZ | citrate synthase | -1.56 | 3.47E-02 |
| SMU_1407c |  | putative transposase, ISSmu1 | -1.68 | 1.73E-03 |
| SMU_1894c |  | conserved hypothetical protein | -1.74 | 3.82E-03 |
| SMU_670 | citB | aconitate hydratase; aconitase | -1.78 | 9.86E-03 |
| SMU_940c |  | putative hemolysin III | -1.88 | 1.59E-17 |
| SMU_941c |  | conserved hypothetical protein | -1.91 | 8.46E-20 |
| SMU_100 |  | putative sorbose PTS system, IIB component | -1.96 | 4.03E-06 |
| SMU_416 |  | conserved hypothetical protein | -2.05 | 4.57E-46 |
| SMU_1184c |  | putative transcriptional regulator, antiterminator | -2.34 | 6.63E-09 |
| SMU_415 |  | conserved hypothetical protein | -2.43 | 3.06E-77 |
| SMU_1408c |  | conserved hypothetical protein | -2.93 | 1.17E-05 |
| SMU_1185 | mtlA1 | PTS system, mannitol-specific enzyme IIBC component | -3.01 | 2.89E-12 |
| SMU_1348c |  | putative ABC transporter, ATP-binding protein | -3.17 | 7.26E-25 |
| SMU_106c |  | putative transposase fragment | -3.56 | 9.34E-61 |
| SMU_1347c |  | conserved hypothetical protein; possible permease | -3.87 | 1.96E-20 |
| SMU_1030 |  | putative polyribonucleotide nucleotidyltransferase; Tn916 ORF8-like | -4.10 | 6.72E-02 |
| SMU_1365c |  | hypothetical protein; possible permease | -4.21 | 1.40E-44 |
| SMU_1366c |  | putative ABC transporter; ATP-binding protein | -7.01 | 1.68E-70 |

**Table S6 Differential gene expression (RNA-seq) analysis of *S. mutans* UA159 during CRISPRi-mediated knockdown of SMU_419**

| **Locus tag** | **Gene** | **Description** | **log2 fold change** | **FDR** |
| --- | --- | --- | --- | --- |
| SMU_1405c | cas9 | conserved hypothetical protein | 8.00 | 6.62E-170 |
| SMU_958 |  | hypothetical protein | 6.42 | 4.23E-43 |
| SMU_935 |  | putative amino acid ABC transporter, permease protein | 3.51 | 8.55E-06 |
| SMU_215c |  | hypothetical protein | 3.47 | 6.54E-06 |
| SMU_2053c |  | hypothetical protein | 3.17 | 1.50E-04 |
| SMU_217c |  | hypothetical protein | 2.96 | 2.90E-09 |
| SMU_2124 |  | hypothetical protein | 2.79 | 4.35E-02 |
| SMU_216c |  | hypothetical protein | 2.75 | 1.28E-04 |
| SMU_264 |  | conserved hypothetical protein | 2.46 | 1.50E-11 |
| SMU_265 | arcC | putative carbamate kinase | 2.45 | 7.55E-14 |
| SMU_214c |  | hypothetical protein | 2.41 | 3.62E-07 |
| SMU_1907 |  | hypothetical protein | 2.41 | 6.92E-03 |
| SMU_262 | otcA | putative ornithine carbamoyltransferase | 2.40 | 9.15E-11 |
| SMU_791c |  | hypothetical protein | 2.37 | 1.59E-09 |
| SMU_1747c |  | putative phosphatase | 2.30 | 2.50E-29 |
| SMU_11 |  | conserved hypothetical protein | 2.26 | 3.43E-04 |
| SMU_1896c |  | hypothetical protein | 2.24 | 3.42E-13 |
| SMU_208c |  | putative transposon protein; possible DNA segregation ATPase | 2.04 | 3.96E-05 |
| SMU_124 |  | putative transcriptional regulator (MarR family) | 1.92 | 5.53E-11 |
| SMU_206c |  | hypothetical protein | 1.91 | 1.55E-03 |
| SMU_150 |  | hypothetical protein | 1.86 | 7.90E-03 |
| SMU_1774c |  | conserved hypothetical protein | 1.86 | 1.79E-13 |
| SMU_263 |  | putative amino acid antiporter | 1.82 | 1.09E-07 |
| SMU_495 | gldA | glycerol dehydrogenase | 1.80 | 1.63E-37 |
| SMU_260 |  | conserved hypothetical protein | 1.80 | 2.04E-09 |
| SMU_982 | bglB2 | putative BglB fragment | 1.77 | 1.95E-05 |
| SMU_213c |  | hypothetical protein | 1.76 | 8.50E-03 |
| SMU_107 |  | hypothetical protein | 1.75 | 1.90E-02 |
| SMU_490 | pflC | putative pyruvate formate-lyase activating enzyme | 1.75 | 3.19E-26 |
| SMU_1196c |  | conserved hypothetical protein | 1.74 | 9.66E-13 |
| SMU_212c |  | hypothetical protein | 1.73 | 1.51E-02 |
| SMU_1895c |  | hypothetical protein | 1.69 | 2.80E-07 |
| SMU_125 |  | conserved hypothetical protein | 1.67 | 2.50E-09 |
| SMU_108 |  | hypothetical protein | 1.59 | 1.76E-04 |
| SMU_28 |  | putative ATP-binding protein | 1.56 | 3.35E-10 |
| SMU_1899 |  | putative ABC transporter, ATP-binding and permease protein (fragment) | 1.55 | 3.97E-01 |
| SMU_383c |  | conserved hypothetical protein; putative reductase | 1.54 | 1.61E-07 |
| SMU_2019 | rl29 | 50S ribosomal protein L29 | 1.54 | 5.95E-16 |
| SMU_1914c |  | hypothetical protein | 1.52 | 1.94E-02 |
| SMU_1250c |  | hypothetical protein | 1.51 | 2.38E-08 |
| SMU_494 |  | putative transaldolase | 1.47 | 2.15E-24 |
| SMU_722 |  | hypothetical protein | 1.45 | 8.73E-13 |
| SMU_382c |  | putative oxidoreductase | 1.44 | 1.40E-06 |
| SMU_1381 | leuD | putative 3-isopropylmalate dehydratase, small subunit | 1.42 | 7.95E-16 |
| SMU_642 |  | hypothetical protein | 1.41 | 9.29E-06 |
| SMU_633 |  | putative thioesterase | 1.35 | 1.35E-11 |
| SMU_655 | mutE1 | putative MutE | 1.35 | 1.38E-02 |
| SMU_211c |  | hypothetical protein | 1.34 | 2.37E-02 |
| SMU_977 | licT | putative transcriptional antiterminator LicT (fragment) | 1.33 | 2.86E-22 |
| SMU_1317c |  | hypothetical protein | 1.32 | 2.43E-10 |
| SMU_1148 |  | putative transporter, ATP-binding protein; bacteriocin immunity protein | 1.32 | 3.52E-04 |
| SMU_2129c |  | conserved hypothetical protein | 1.31 | 4.28E-08 |
| SMU_189 |  | hypothetical protein | 1.31 | 4.62E-02 |
| SMU_1299c |  | putative acetate kinase | 1.30 | 1.32E-10 |
| SMU_369c |  | conserved hypothetical protein | 1.29 | 6.66E-08 |
| SMU_93c |  | hypothetical protein; putative transposase fragment | 1.26 | 1.89E-02 |
| SMU_223c |  | hypothetical protein | 1.24 | 2.43E-06 |
| SMU_930c |  | putative transcriptional regulator | 1.22 | 2.03E-05 |
| SMU_998 |  | putative ABC transporter, periplasmic ferrichrome-binding protein | 1.22 | 1.59E-26 |
| SMU_94c |  | hypothetical protein; putative transposase fragment | 1.22 | 3.10E-02 |
| SMU_947 | dfrA | putative dihydrofolate reductase | 1.20 | 5.37E-11 |
| SMU_632 |  | putative transcriptional regulator | 1.19 | 3.35E-10 |
| SMU_657 | mutG | putative MutG | 1.19 | 4.10E-05 |
| SMU_1902c |  | hypothetical protein | 1.18 | 1.39E-02 |
| SMU_727 |  | putative transcriptional regulator | 1.17 | 2.84E-07 |
| SMU_605 |  | hypothetical protein | 1.16 | 2.90E-08 |
| SMU_1456c |  | hypothetical protein | 1.16 | 2.19E-03 |
| SMU_641 |  | putative oxidoreductase | 1.16 | 1.45E-04 |
| SMU_1604c |  | conserved hypothetical protein | 1.15 | 2.51E-05 |
| SMU_768c |  | conserved hypothetical protein | 1.14 | 1.09E-07 |
| SMU_285 |  | hypothetical protein | 1.14 | 6.67E-06 |
| SMU_55 |  | hypothetical protein | 1.14 | 3.46E-05 |
| SMU_1775c |  | hypothetical protein | 1.13 | 2.37E-06 |
| SMU_491 |  | putative DeoR-type transcriptional regulator | 1.12 | 5.23E-12 |
| SMU_687c |  | hypothetical protein | 1.11 | 9.38E-02 |
| SMU_1147c |  | hypothetical protein | 1.10 | 6.02E-05 |
| SMU_393 |  | conserved hypothetical protein | 1.10 | 2.13E-07 |
| SMU_1056 |  | hypothetical protein | 1.10 | 1.76E-04 |
| SMU_1475c |  | conserved hypothetical protein | 1.10 | 2.04E-08 |
| SMU_803c |  | putative ABC transporter, ATP-binding protein | 1.09 | 2.46E-08 |
| SMU_1832 |  | hypothetical protein | 1.09 | 9.45E-09 |
| SMU_283 |  | hypothetical protein | 1.08 | 1.26E-07 |
| SMU_1382 | leuC | putative 3-isopropylmalate dehydratase, large subunit | 1.07 | 6.28E-16 |
| SMU_750c |  | hypothetical protein | 1.07 | 2.63E-02 |
| SMU_1402c |  | conserved hypothetical protein | 1.06 | 6.69E-04 |
| SMU_1300c |  | conserved hypothetical protein | 1.05 | 4.18E-10 |
| SMU_1097c |  | putative transcriptional regulator protein | 1.05 | 1.10E-08 |
| SMU_1080c |  | conserved hypothetical protein; possible transposon-related protein | 1.05 | 2.53E-13 |
| SMU_1109c |  | putative integral membrane protein; possible permease | 1.04 | 1.58E-14 |
| SMU_711 |  | conserved hypothetical protein | 1.04 | 1.11E-06 |
| SMU_1999c |  | conserved hypothetical protein | 1.04 | 2.53E-11 |
| SMU_417 |  | conserved hypothetical protein | 1.04 | 1.81E-12 |
| SMU_2137c |  | conserved hypothetical protein | 1.03 | 8.30E-08 |
| SMU_1249c |  | hypothetical protein | 1.03 | 5.23E-06 |
| SMU_654 | mutF | putative ABC transporter, ATP-binding protein MutF | 1.02 | 7.76E-04 |
| SMU_798c |  | hypothetical protein | 1.02 | 2.92E-02 |
| SMU_653c |  | putative ABC transporter, permease protein | 1.02 | 1.06E-06 |
| SMU_1621c |  | conserved hypothetical protein | 1.01 | 1.93E-04 |
| SMU_2036 | pepO | putative peptidase | -1.01 | 1.20E-09 |
| SMU_1116c |  | hypothetical protein | -1.01 | 1.85E-02 |
| SMU_530c |  | conserved hypothetical protein | -1.02 | 1.06E-10 |
| SMU_1571 |  | putative ABC transporter, ATP-binding protein, MsmK-like protein | -1.02 | 7.86E-10 |
| SMU_1770 | syv | putative valyl-tRNA synthetase | -1.05 | 9.86E-12 |
| SMU_924 | tpx | thiol peroxidase | -1.05 | 3.65E-03 |
| SMU_2047 | ptsG | putative PTS system, glucose-specific IIABC component | -1.06 | 8.52E-08 |
| SMU_1537 | glgD | putative glycogen biosynthesis protein GlgD | -1.06 | 2.25E-03 |
| SMU_129 | adhC | putative dihydrolipoamide acetyltransferase | -1.06 | 1.95E-02 |
| SMU_2074 | nrdD | putative anaerobic ribonucleoside-triphosphate reductase | -1.06 | 2.79E-11 |
| SMU_2028 | sacB | levansucrase precursor; beta-D-fructosyltransferase | -1.06 | 1.41E-11 |
| SMU_1946 |  | hypothetical protein | -1.07 | 1.30E-07 |
| SMU_510c |  | hypothetical protein | -1.08 | 2.92E-05 |
| SMU_1928 | psaB | putative ABC transporter, permease protein | -1.08 | 1.15E-02 |
| SMU_1817c |  | putative maturase-related protein | -1.09 | 6.83E-07 |
| SMU_1821c |  | putative glutamyl-tRNA (Gln) amidotransferase subunit C | -1.10 | 1.06E-11 |
| SMU_667 | nrdG | putative ribonucleotide reductase, small subunit | -1.11 | 8.55E-06 |
| SMU_242c |  | putative amino acid ABC transporter, permease protein, glutamine transport system | -1.11 | 1.96E-06 |
| SMU_541 |  | conserved hypothetical protein | -1.12 | 7.22E-04 |
| SMU_1262c |  | hypothetical protein | -1.12 | 4.61E-05 |
| SMU_1945 |  | hypothetical protein | -1.12 | 3.38E-09 |
| SMU_99 | fbaA | fructose-1,6-biphosphate aldolase | -1.13 | 3.07E-14 |
| SMU_130 | adhD | putative dihydrolipoamide dehydrogenase | -1.16 | 4.82E-03 |
| SMU_1533 | atpG | FoF1 membrane-bound proton-translocating ATPase, a subunit | -1.19 | 4.43E-12 |
| SMU_1869 | trxA | putative thioredoxin | -1.19 | 1.37E-04 |
| SMU_404c |  | hypothetical protein | -1.20 | 2.16E-04 |
| SMU_166 |  | hypothetical protein | -1.20 | 1.55E-07 |
| SMU_128 | adhB | putative acetoin dehydrogenase (TPP-dependent), E1 component beta subunit | -1.20 | 3.49E-03 |
| SMU_650 |  | putative alanyl-tRNA synthetase (alanine--tRNA ligase) | -1.21 | 8.54E-19 |
| SMU_589 |  | putative DNA-binding protein | -1.22 | 7.28E-13 |
| SMU_1088 | apbE | putative thiamine biosynthesis lipoprotein | -1.24 | 2.38E-09 |
| SMU_674 | ptsH | phosphoenolpyruvate:sugar phosphotransferase system HPr | -1.25 | 4.76E-10 |
| SMU_1867c |  | putative alcohol dehydrogenase | -1.25 | 2.06E-06 |
| SMU_668c |  | ribonucleotide reductase, large subunit | -1.26 | 3.71E-07 |
| SMU_1265 | hisA | putative phosphoribosyl formimino-5-aminoimidazole carboxamide ribonucleotide isomerase | -1.26 | 2.45E-17 |
| SMU_496 | cysK | putative cysteine synthetase A; O-acetylserine lyase | -1.27 | 3.61E-05 |
| SMU_1879 |  | putative PTS system, mannose-specific component IID | -1.27 | 3.14E-12 |
| SMU_1398 |  | putative transcriptional regulator | -1.28 | 9.44E-10 |
| SMU_1805 |  | putative transcriptional regulator | -1.29 | 3.71E-08 |
| SMU_1077 | pgm | putative phosphoglucomutase | -1.30 | 1.69E-08 |
| SMU_402 | pfl | pyruvate formate-lyase | -1.31 | 3.33E-08 |
| SMU_1570 | malG | putative maltose/maltodextrin ABC transporter, MalG permease | -1.31 | 2.16E-12 |
| SMU_649 |  | conserved hypothetical protein | -1.36 | 9.86E-22 |
| SMU_1968c |  | conserved hypothetical protein | -1.36 | 5.78E-09 |
| SMU_675 |  | phosphoenolpyruvate:sugar phosphotransferase system enzyme I, PTS system EI component | -1.36 | 5.49E-20 |
| SMU_1263 | hisI | putative phosphoribosyl-ATP pyrophosphatase / phosphoribosyl-AMP cyclohydrolase | -1.36 | 7.51E-16 |
| SMU_1819 | gatB | putative glutamyl-tRNA (Gln) amidotransferase subunit B | -1.37 | 3.50E-33 |
| SMU_610 | spaP | cell surface antigen SpaP | -1.37 | 6.29E-12 |
| SMU_558 |  | isoleucine-tRNA synthetase | -1.38 | 1.35E-33 |
| SMU_1396 | gbpC | glucan-binding protein C, GbpC | -1.38 | 2.06E-08 |
| SMU_102 |  | putative PTS system, IID component | -1.38 | 7.60E-07 |
| SMU_672 | idh | isocitrate dehydrogenase | -1.39 | 4.30E-02 |
| SMU_360 | gapC | extracellular glyceraldehyde-3-phosphate dehydrogenase | -1.39 | 9.01E-29 |
| SMU_1185 | mtlA1 | PTS system, mannitol-specific enzyme IIBC component | -1.39 | 4.00E-04 |
| SMU_670 | citB | aconitate hydratase; aconitase | -1.40 | 2.95E-02 |
| SMU_1266 | hisH | putative glutamine amidotransferase HisH | -1.41 | 8.54E-19 |
| SMU_148 | adhE | putative alcohol-acetaldehyde dehydrogenase | -1.42 | 3.81E-03 |
| SMU_671 | citZ | citrate synthase | -1.44 | 3.52E-02 |
| SMU_405c |  | putative transcriptional regulator | -1.46 | 1.04E-04 |
| SMU_1270 | hisD | putative histidinol dehydrogenase | -1.48 | 1.51E-19 |
| SMU_1268 | hisB | putative imidazoleglycerol-phosphate dehydratase | -1.48 | 6.00E-17 |
| SMU_1269 | serB | putative phosphoserine phosphatase | -1.52 | 4.77E-19 |
| SMU_1599 | celR | putative transcriptional regulator; possible antiterminator | -1.53 | 7.59E-03 |
| SMU_956 | clp | putative Clp-like ATP-dependent protease, ATP-binding subunit | -1.54 | 2.12E-06 |
| SMU_1569 | malF | putative maltose/maltodextrin ABC transporter, permease protein MalF | -1.54 | 4.98E-15 |
| SMU_1820c |  | putative glutamyl-tRNA(Gln) amidotransferase A subunit | -1.55 | 4.77E-31 |
| SMU_420 |  | putative ribosomal protein | -1.57 | 6.66E-08 |
| SMU_1267c |  | hypothetical protein | -1.58 | 2.02E-18 |
| SMU_1539 | glgB | putative 1,4-alpha-glucan branching enzyme | -1.59 | 1.05E-06 |
| SMU_1271 | hisG | putative ATP phosphoribosyltransferase | -1.64 | 7.98E-19 |
| SMU_252 |  | hypothetical protein | -1.66 | 1.71E-14 |
| SMU_1272 | hisZ | putative histidyl-tRNA synthetase | -1.66 | 1.59E-18 |
| SMU_871 | pfkB | putative fructose-1-phosphate kinase | -1.71 | 1.80E-14 |
| SMU_1247 | eno | putative enolase | -1.71 | 2.50E-37 |
| SMU_1004 | gtfB | glucosyltransferase-I | -1.79 | 1.57E-12 |
| SMU_1587c |  | hypothetical protein | -1.83 | 4.02E-36 |
| SMU_1511c |  | putative acetyltransferase | -1.83 | 7.47E-22 |
| SMU_1273 | hisC | putative histidinol-phosphate aminotransferase | -1.84 | 1.70E-21 |
| SMU_1878 | ptnC | putative PTS system, mannose-specific component IIC | -1.85 | 5.49E-20 |
| SMU_1407c |  | putative transposase, ISSmu1 | -1.86 | 3.16E-04 |
| SMU_419 |  | conserved hypothetical protein | -1.88 | 8.54E-17 |
| SMU_101 |  | putative sorbose PTS system, IIC component | -1.90 | 1.75E-08 |
| SMU_1586 | syt1 | putative threonyl-tRNA synthetase | -1.90 | 6.76E-37 |
| SMU_2127 |  | putative succinate semialdehyde dehydrogenase | -1.92 | 1.39E-14 |
| SMU_1117 | naoX | NADH oxidase (H2O-forming) | -1.93 | 2.30E-05 |
| SMU_1184c |  | putative transcriptional regulator, antiterminator | -1.96 | 5.75E-07 |
| SMU_503c |  | hypothetical protein | -1.97 | 3.41E-15 |
| SMU_89c |  | putative nitrite transporter | -2.03 | 2.04E-16 |
| SMU_1822 | gatA | putative aspartyl-tRNA synthetase | -2.07 | 1.08E-40 |
| SMU_1568 | malX | putative maltose/maltodextrin ABC transporter, sugar-binding protein MalX | -2.08 | 2.27E-16 |
| SMU_1600 | ptcB | putative PTS system, cellobiose-specific IIB component | -2.12 | 5.70E-04 |
| SMU_183 | sloB | putative Mn/Zn ABC transporter | -2.14 | 4.02E-11 |
| SMU_422 |  | ribosome binding factor A | -2.16 | 1.02E-39 |
| SMU_1510 | syfB | putative phenylalanyl-tRNA synthetase (beta subunit) | -2.23 | 1.06E-44 |
| SMU_1877 | ptnA | putative PTS system, mannose-specific component IIAB | -2.24 | 5.81E-26 |
| SMU_609 |  | putative 40K cell wall protein precursor | -2.32 | 1.07E-07 |
| SMU_940c |  | putative hemolysin III | -2.33 | 1.20E-26 |
| SMU_1512 | syfA | putative phenylalanyl-tRNA synthetase (alpha subunit) | -2.34 | 4.10E-34 |
| SMU_626 |  | putative competence protein | -2.42 | 2.69E-02 |
| SMU_1980c |  | conserved hypothetical protein | -2.50 | 2.97E-02 |
| SMU_179 |  | conserved hypothetical protein | -2.52 | 6.66E-08 |
| SMU_1984 | comYC | putative competence protein ComYC | -2.52 | 2.94E-02 |
| SMU_1967 | ssb2 | putative single-stranded DNA-binding protein | -2.53 | 1.31E-03 |
| SMU_1983 | comYD | putative competence protein ComYD | -2.56 | 2.47E-02 |
| SMU_421 |  | translation initiation factor 2 | -2.59 | 3.26E-73 |
| SMU_1987 | comYA | putative ABC transporter, ATP-binding protein ComYA; late competence gene | -2.66 | 1.59E-02 |
| SMU_106c |  | putative transposase fragment | -2.74 | 1.70E-41 |
| SMU_1408c |  | conserved hypothetical protein | -2.80 | 1.33E-05 |
| SMU_500 |  | putative ribosome-associated protein | -2.82 | 1.02E-27 |
| SMU_1894c |  | conserved hypothetical protein | -2.84 | 2.72E-06 |
| SMU_1425 | clpB | putative Clp proteinase, ATP-binding subunit ClpB | -2.93 | 2.18E-10 |
| SMU_1985 | comYB | putative ABC transporter ComYB; probably part of the DNA transport machinery | -2.94 | 1.04E-02 |
| SMU_1982c |  | conserved hypothetical protein | -2.94 | 1.41E-02 |
| SMU_941c |  | conserved hypothetical protein | -2.94 | 2.34E-43 |
| SMU_870 |  | putative transcriptional regulator of sugar metabolism | -3.06 | 3.12E-27 |
| SMU_1424 | pdhD | putative dihydrolipoamide dehydrogenase | -3.10 | 6.20E-06 |
| SMU_100 |  | putative sorbose PTS system, IIB component | -3.15 | 3.03E-14 |
| SMU_575c |  | putative membrane protein | -3.17 | 2.42E-03 |
| SMU_1001 | smf | putative DNA processing Smf protein | -3.24 | 2.65E-03 |
| SMU_182 | sloA | putative ABC transporter, ATP-binding protein; possible iron and/or manganese ABC transport system | -3.26 | 9.71E-13 |
| SMU_1981c |  | conserved hypothetical protein | -3.29 | 5.68E-03 |
| SMU_1348c |  | putative ABC transporter, ATP-binding protein | -4.01 | 2.09E-36 |
| SMU_1347c |  | conserved hypothetical protein; possible permease | -4.11 | 4.95E-23 |
| SMU_1365c |  | hypothetical protein; possible permease | -4.47 | 1.67E-48 |
| SMU_1366c |  | putative ABC transporter; ATP-binding protein | -7.39 | 1.09E-72 |

**Table S7 Differential gene expression (RNA-seq) analysis of *S. mutans* UA159 during CRISPRi-mediated knockdown of SMU_1801c**

| **Locus tag** | **Gene** | **Description** | **log2 fold change** | **FDR** |
| --- | --- | --- | --- | --- |
| SMU_1405c | cas9 | conserved hypothetical protein | 7.86 | 3.24E-166 |
| SMU_958 |  | hypothetical protein | 6.12 | 5.71E-42 |
| SMU_1899 |  | putative ABC transporter, ATP-binding and permease protein (fragment) | 3.99 | 2.86E-02 |
| SMU_2053c |  | hypothetical protein | 3.44 | 5.16E-04 |
| SMU_206c |  | hypothetical protein | 3.26 | 3.53E-09 |
| SMU_212c |  | hypothetical protein | 3.17 | 6.17E-07 |
| SMU_217c |  | hypothetical protein | 3.11 | 1.22E-10 |
| SMU_213c |  | hypothetical protein | 3.02 | 2.21E-06 |
| SMU_193c |  | conserved hypothetical protein | 2.89 | 3.08E-03 |
| SMU_1854 |  | conserved hypothetical protein | 2.85 | 6.15E-36 |
| SMU_215c |  | hypothetical protein | 2.78 | 5.82E-05 |
| SMU_1855 |  | hypothetical protein | 2.76 | 2.12E-33 |
| SMU_204c |  | hypothetical protein | 2.66 | 9.67E-07 |
| SMU_214c |  | hypothetical protein | 2.62 | 2.55E-09 |
| SMU_194c |  | conserved hypothetical protein; Bacteriophage P2 associated | 2.58 | 2.27E-05 |
| SMU_150 |  | hypothetical protein | 2.58 | 2.08E-04 |
| SMU_1895c |  | hypothetical protein | 2.48 | 7.36E-14 |
| SMU_655 | mutE1 | putative MutE | 2.45 | 5.59E-06 |
| SMU_200c |  | hypothetical protein | 2.42 | 1.90E-04 |
| SMU_1896c |  | hypothetical protein | 2.38 | 4.42E-14 |
| SMU_202c |  | hypothetical protein | 2.34 | 1.79E-05 |
| SMU_1774c |  | conserved hypothetical protein | 2.33 | 4.90E-20 |
| SMU_1250c |  | hypothetical protein | 2.31 | 1.43E-17 |
| SMU_1914c |  | hypothetical protein | 2.30 | 2.97E-04 |
| SMU_495 | gldA | glycerol dehydrogenase | 2.26 | 8.99E-58 |
| SMU_191c |  | putative integrase | 2.20 | 8.97E-05 |
| SMU_216c |  | hypothetical protein | 2.15 | 9.91E-04 |
| SMU_2019 | rl29 | 50S ribosomal protein L29 | 2.13 | 3.23E-29 |
| SMU_896 |  | conserved hypothetical protein | 2.08 | 1.79E-09 |
| SMU_494 |  | putative transaldolase | 2.04 | 5.56E-46 |
| SMU_196c |  | putative transfer protein | 1.94 | 4.37E-04 |
| SMU_201c |  | putative transposon protein | 1.93 | 1.53E-04 |
| SMU_490 | pflC | putative pyruvate formate-lyase activating enzyme | 1.92 | 7.81E-32 |
| SMU_2129c |  | conserved hypothetical protein | 1.90 | 1.32E-15 |
| SMU_195c |  | hypothetical protein | 1.90 | 6.27E-05 |
| SMU_379 |  | hypothetical protein | 1.88 | 6.20E-02 |
| SMU_2037 | treA | putative trehalose-6-phosphate hydrolase TreA | 1.88 | 2.44E-15 |
| SMU_1761c |  | conserved hypothetical protein | 1.88 | 7.50E-11 |
| SMU_81 | grpE | heat shock protein GrpE (HSP-70 cofactor) | 1.87 | 1.44E-09 |
| SMU_382c |  | putative oxidoreductase | 1.85 | 4.97E-10 |
| SMU_1977c |  | putative transcriptional regulator | 1.84 | 1.34E-32 |
| SMU_1752c |  | hypothetical protein | 1.83 | 3.80E-13 |
| SMU_383c |  | conserved hypothetical protein; putative reductase | 1.79 | 6.93E-10 |
| SMU_791c |  | hypothetical protein | 1.76 | 3.67E-06 |
| SMU_211c |  | hypothetical protein | 1.75 | 1.74E-03 |
| SMU_1155 |  | hypothetical protein | 1.74 | 4.16E-05 |
| SMU_1862 |  | hypothetical protein | 1.72 | 5.29E-13 |
| SMU_1174 | pcrA | ATP-dependent DNA helicase | 1.69 | 1.15E-44 |
| SMU_803c |  | putative ABC transporter, ATP-binding protein | 1.68 | 1.39E-18 |
| SMU_205c |  | hypothetical protein | 1.67 | 1.89E-03 |
| SMU_153 |  | hypothetical protein | 1.67 | 6.73E-03 |
| SMU_735 |  | hypothetical protein | 1.65 | 9.91E-10 |
| SMU_1912c |  | hypothetical protein | 1.65 | 1.83E-02 |
| SMU_11 |  | conserved hypothetical protein | 1.65 | 8.91E-03 |
| SMU_1762c |  | conserved hypothetical protein | 1.63 | 4.05E-07 |
| SMU_491 |  | putative DeoR-type transcriptional regulator | 1.61 | 4.17E-24 |
| SMU_393 |  | conserved hypothetical protein | 1.60 | 3.32E-14 |
| SMU_2105 |  | hypothetical protein | 1.58 | 8.58E-17 |
| SMU_711 |  | conserved hypothetical protein | 1.57 | 1.17E-14 |
| SMU_1763c |  | conserved hypothetical protein | 1.57 | 2.15E-09 |
| SMU_665 | argB | putative acetylglutamate kinase | 1.57 | 2.22E-09 |
| SMU_208c |  | putative transposon protein; possible DNA segregation ATPase | 1.57 | 1.26E-03 |
| SMU_220c |  | hypothetical protein | 1.55 | 6.80E-16 |
| SMU_493 | pfl2 | formate acetyltransferase (pyruvate formate-lyase 2) | 1.55 | 4.86E-31 |
| SMU_768c |  | conserved hypothetical protein | 1.54 | 3.18E-13 |
| SMU_1237c |  | hypothetical protein | 1.54 | 1.63E-06 |
| SMU_223c |  | hypothetical protein | 1.54 | 3.41E-09 |
| SMU_1906c |  | hypothetical protein | 1.51 | 6.52E-02 |
| SMU_1354c |  | putative transposase fragment | 1.50 | 5.37E-12 |
| SMU_698 |  | 50S ribosomal protein L35 | 1.50 | 5.35E-15 |
| SMU_1249c |  | hypothetical protein | 1.48 | 2.51E-11 |
| SMU_94c |  | hypothetical protein; putative transposase fragment | 1.46 | 7.97E-03 |
| SMU_1161c |  | hypothetical protein | 1.46 | 4.75E-07 |
| SMU_753 |  | conserved hypothetical protein | 1.44 | 1.72E-08 |
| SMU_947 | dfrA | putative dihydrofolate reductase | 1.43 | 5.32E-15 |
| SMU_605 |  | hypothetical protein | 1.42 | 1.11E-11 |
| SMU_115 |  | putative PTS system, fructose-specific IIA component | 1.42 | 7.89E-02 |
| SMU_1604c |  | conserved hypothetical protein | 1.41 | 2.39E-07 |
| SMU_849 |  | 50S ribosomal protein L27 | 1.41 | 1.66E-17 |
| SMU_225c |  | hypothetical protein | 1.40 | 5.87E-09 |
| SMU_281 |  | hypothetical protein | 1.39 | 3.28E-06 |
| SMU_1159c |  | hypothetical protein | 1.37 | 1.25E-05 |
| SMU_1475c |  | conserved hypothetical protein | 1.36 | 2.59E-12 |
| SMU_2038 | pttB | putative PTS system, trehalose-specific IIABC component | 1.36 | 1.51E-09 |
| SMU_1172c |  | conserved hypothetical protein | 1.34 | 1.08E-28 |
| SMU_72 |  | conserved hypothetical protein | 1.34 | 9.30E-10 |
| SMU_1754c |  | conserved hypothetical protein | 1.33 | 5.62E-09 |
| SMU_591c |  | hypothetical protein | 1.33 | 6.00E-09 |
| SMU_1764c |  | conserved hypothetical protein | 1.33 | 6.69E-08 |
| SMU_722 |  | hypothetical protein | 1.32 | 6.77E-11 |
| SMU_642 |  | hypothetical protein | 1.32 | 3.21E-05 |
| SMU_209c |  | hypothetical protein | 1.31 | 1.06E-02 |
| SMU_1374 |  | hypothetical protein | 1.31 | 2.79E-05 |
| SMU_1419 |  | putative transcriptional regulator | 1.30 | 1.61E-04 |
| SMU_654 | mutF | putative ABC transporter, ATP-binding protein MutF | 1.29 | 6.97E-06 |
| SMU_659 |  | putative response regulator SpaR | 1.29 | 1.50E-14 |
| SMU_1757c |  | conserved hypothetical protein | 1.29 | 1.75E-07 |
| SMU_2104a |  | 50S ribosomal protein L32 | 1.28 | 1.13E-14 |
| SMU_657 | mutG | putative MutG | 1.28 | 2.87E-06 |
| SMU_1378 |  | hypothetical protein | 1.28 | 9.16E-04 |
| SMU_1502c |  | conserved hypothetical protein | 1.27 | 1.62E-11 |
| SMU_730 |  | conserved hypothetical protein | 1.25 | 4.94E-04 |
| SMU_1905c |  | putative bacteriocin secretion protein | 1.24 | 6.07E-02 |
| SMU_185 |  | hypothetical protein | 1.22 | 1.11E-04 |
| SMU_661 |  | putative transcriptional regulator | 1.22 | 1.42E-12 |
| SMU_1149 |  | putative transporter, trans-membrane domain bacteriocin immunity protein | 1.21 | 2.76E-03 |
| SMU_1892c |  | hypothetical protein | 1.21 | 1.55E-01 |
| SMU_283 |  | hypothetical protein | 1.21 | 2.94E-09 |
| SMU_1904c |  | hypothetical protein | 1.21 | 6.20E-02 |
| SMU_207c |  | putative transposon protein | 1.20 | 1.35E-02 |
| SMU_80 | hrcA | transcriptional regulator; repressor (HrcA) of class I heat shock genes | 1.20 | 4.90E-05 |
| SMU_1760c |  | conserved hypothetical protein | 1.18 | 1.41E-06 |
| SMU_844 |  | conserved hypothetical protein | 1.18 | 7.46E-13 |
| SMU_420 |  | putative ribosomal protein | 1.18 | 2.35E-05 |
| SMU_325 |  | putative dUTPase | 1.18 | 1.27E-13 |
| SMU_1173 | cysD | putative O-acetylhomoserine sulfhydrylase | 1.17 | 2.82E-18 |
| SMU_439 |  | putative transcriptional regulator | 1.17 | 7.49E-09 |
| SMU_56 |  | conserved hypothetical protein | 1.17 | 8.18E-06 |
| SMU_1913c |  | putative immunity protein, BLpL-like | 1.17 | 8.81E-02 |
| SMU_1027 |  | putative transcription regulator | 1.17 | 1.01E-05 |
| SMU_1363c |  | putative transposase | 1.16 | 1.00E-06 |
| SMU_1064c |  | putative transcriptional regulator (GntR family) | 1.16 | 3.97E-07 |
| SMU_68 |  | hypothetical protein | 1.16 | 6.54E-11 |
| SMU_1160c |  | hypothetical protein | 1.15 | 3.00E-05 |
| SMU_93c |  | hypothetical protein; putative transposase fragment | 1.15 | 2.17E-02 |
| SMU_198c |  | putative conjugative transposon protein | 1.15 | 2.70E-02 |
| SMU_1256c |  | hypothetical protein | 1.15 | 8.89E-08 |
| SMU_112c |  | putative transcriptional regulator | 1.15 | 1.44E-09 |
| SMU_434 |  | hypothetical protein | 1.15 | 7.64E-11 |
| SMU_197c |  | hypothetical protein | 1.14 | 2.66E-02 |
| SMU_1910c |  | hypothetical protein | 1.14 | 1.29E-01 |
| SMU_2061 |  | hypothetical protein | 1.14 | 9.18E-09 |
| SMU_1908c |  | hypothetical protein | 1.14 | 1.16E-01 |
| SMU_666 | argD | putative N-acetylornithine aminotransferase | 1.14 | 2.23E-08 |
| SMU_1547c |  | putative response regulator | 1.13 | 2.91E-12 |
| SMU_660 |  | putative histidine kinase SpaK | 1.13 | 1.18E-13 |
| SMU_2146c |  | hypothetical protein | 1.12 | 2.14E-13 |
| SMU_108 |  | hypothetical protein | 1.11 | 1.05E-02 |
| SMU_1417c |  | putative oleoyl-acyl carrier protein thioesterase | 1.11 | 9.88E-14 |
| SMU_2126c |  | putative purine-nucleoside phosphorylase | 1.11 | 1.34E-11 |
| SMU_1317c |  | hypothetical protein | 1.10 | 9.38E-08 |
| SMU_1775c |  | hypothetical protein | 1.10 | 5.04E-06 |
| SMU_887 | galT | galactose-1-P-uridyl transferase, GalT | 1.10 | 5.77E-07 |
| SMU_433 |  | putative transcriptional regulator | 1.10 | 3.60E-13 |
| SMU_149 |  | putative transposase | 1.09 | 1.44E-02 |
| SMU_1832 |  | hypothetical protein | 1.08 | 7.03E-09 |
| SMU_1148 |  | putative transporter, ATP-binding protein; bacteriocin immunity protein | 1.07 | 1.69E-03 |
| SMU_1219c |  | conserved hypothetical protein | 1.07 | 1.30E-10 |
| SMU_1178c |  | putative amino acid ABC transporter, ATP-binding protein | 1.06 | 2.59E-16 |
| SMU_2137c |  | conserved hypothetical protein | 1.06 | 2.81E-08 |
| SMU_1356c |  | putative transposase fragment | 1.05 | 3.67E-05 |
| SMU_685 |  | hypothetical protein | 1.05 | 3.55E-05 |
| SMU_52 |  | conserved hypothetical protein | 1.05 | 5.41E-06 |
| SMU_294 |  | conserved hypothetical protein | 1.04 | 9.76E-08 |
| SMU_662 |  | conserved hypothetical protein; possible membrane protein | 1.04 | 6.65E-11 |
| SMU_432 |  | putative ABC transporter, integral membrane protein | 1.04 | 2.10E-09 |
| SMU_606 |  | hypothetical protein | 1.02 | 9.50E-12 |
| SMU_739c |  | hypothetical protein | 1.02 | 7.25E-06 |
| SMU_789 |  | conserved hypothetical protein | 1.01 | 3.88E-13 |
| SMU_684 |  | hypothetical protein | 1.01 | 3.34E-04 |
| SMU_930c |  | putative transcriptional regulator | 1.01 | 2.91E-04 |
| SMU_1299c |  | putative acetate kinase | 1.01 | 1.35E-06 |
| SMU_1080c |  | conserved hypothetical protein; possible transposon-related protein | 1.01 | 2.18E-12 |
| SMU_695 |  | conserved hypothetical protein | 1.00 | 1.57E-15 |
| SMU_199c |  | hypothetical protein | 1.00 | 4.64E-02 |
| SMU_2120c |  | putative 3-methyladenine DNA glycosylase | 1.00 | 1.47E-08 |
| SMU_249 | nifS | putative NifS protein-like protein class-V aminotransferase | -1.00 | 3.53E-08 |
| SMU_1182 | mtlD | mannitol-1-phosphate dehydrogenase | -1.00 | 1.96E-01 |
| SMU_670 | citB | aconitate hydratase; aconitase | -1.00 | 1.20E-01 |
| SMU_1602 |  | putative NAD(P)H-flavin oxidoreductase | -1.01 | 3.50E-09 |
| SMU_1073 | fthS | putative formyl-tetrahydrofolate synthetase | -1.01 | 3.76E-05 |
| SMU_447 |  | conserved hypothetical protein | -1.01 | 3.08E-06 |
| SMU_618 |  | hypothetical protein | -1.01 | 9.91E-06 |
| SMU_2117 | opuCb | putative osmoprotectant ABC transporter; permease protein | -1.02 | 8.41E-11 |
| SMU_1571 |  | putative ABC transporter, ATP-binding protein, MsmK-like protein | -1.03 | 4.11E-10 |
| SMU_252 |  | hypothetical protein | -1.04 | 1.42E-06 |
| SMU_596 | pmgY | phosphoglyceromutase | -1.04 | 6.62E-16 |
| SMU_35 | purN | putative phosphoribosylglycinamide formyltransferase (GART) | -1.05 | 1.68E-04 |
| SMU_674 | ptsH | phosphoenolpyruvate:sugar phosphotransferase system HPr | -1.05 | 1.69E-07 |
| SMU_250 | nifU | putative nitrogen fixation-like protein, NifU | -1.05 | 4.29E-07 |
| SMU_1788c |  | putative bacterocin transport accessory protein, Bta | -1.05 | 6.60E-04 |
| SMU_1034c |  | putative integrase/recombinase; XerC-like | -1.05 | 1.84E-04 |
| SMU_251 |  | conserved hypothetical protein; possible ABC transporter, membrane component | -1.05 | 2.43E-10 |
| SMU_562 | clpE | ATP-dependent protease ClpE | -1.05 | 1.18E-12 |
| SMU_1357 |  | putative transposase fragment | -1.06 | 2.87E-01 |
| SMU_299c |  | putative bacteriocin peptide precursor | -1.06 | 4.30E-04 |
| SMU_248 |  | putative ABC transporter, membrane protein | -1.06 | 1.68E-07 |
| SMU_1452 | alsS | alpha-acetolactate synthase | -1.06 | 7.88E-06 |
| SMU_766 |  | conserved hypothetical protein | -1.06 | 1.11E-01 |
| SMU_446 | sygB | putative glycyl-tRNA synthetase (beta subunit) | -1.06 | 2.37E-10 |
| SMU_1042 |  | conserved hypothetical protein; inner membrane protein | -1.06 | 7.90E-09 |
| SMU_1531 | atpE | FoF1 membrane-bound proton-translocating ATPase, delta subunit | -1.06 | 1.73E-10 |
| SMU_32 | purF | phosphoribosylpyrophosphate amidotransferase | -1.07 | 4.12E-05 |
| SMU_131 | lplA | putative lipoate-protein ligase | -1.08 | 4.81E-03 |
| SMU_1822 | gatA | putative aspartyl-tRNA synthetase | -1.09 | 1.85E-12 |
| SMU_1269 | serB | putative phosphoserine phosphatase | -1.09 | 1.45E-10 |
| SMU_1265 | hisA | putative phosphoribosyl formimino-5-aminoimidazole carboxamide ribonucleotide isomerase | -1.09 | 2.81E-13 |
| SMU_1954 | groEL | putative chaperonin GroEL | -1.09 | 2.08E-08 |
| SMU_167 |  | hypothetical protein | -1.10 | 1.73E-05 |
| SMU_1722c |  | putative integral membrane protein | -1.10 | 1.21E-16 |
| SMU_676 | gapN | NADP-dependent glyceraldehyde-3-phosphate dehydrogenase | -1.10 | 3.53E-09 |
| SMU_629 | sod | putative manganese-type superoxide dismutase, Fe/Mn-SOD | -1.11 | 2.93E-03 |
| SMU_391c |  | conserved hypothetical protein | -1.11 | 5.88E-08 |
| SMU_1297 |  | conserved hypothetical protein | -1.11 | 6.46E-04 |
| SMU_36 |  | conserved hypothetical protein | -1.12 | 1.35E-07 |
| SMU_1841 | scrA | putative PTS system, sucrose-specific IIABC component | -1.13 | 1.27E-13 |
| SMU_1510 | syfB | putative phenylalanyl-tRNA synthetase (beta subunit) | -1.13 | 6.39E-13 |
| SMU_1246c |  | putative transcriptional regulator | -1.13 | 7.44E-12 |
| SMU_1266 | hisH | putative glutamine amidotransferase HisH | -1.15 | 7.46E-13 |
| SMU_2094c |  | conserved hypothetical protein | -1.15 | 3.55E-05 |
| SMU_2102 | hisS | histidyl-tRNA synthetase (histidine--tRNA ligase) | -1.16 | 3.53E-13 |
| SMU_818 |  | 30S ribosomal protein S21 | -1.17 | 5.03E-07 |
| SMU_865 |  | 30S ribosomal protein S16 | -1.17 | 1.05E-12 |
| SMU_445 | sygA | putative glycyl-tRNA synthetase (alpha subunit) | -1.17 | 2.08E-11 |
| SMU_120 |  | 50S ribosomal protein L28 | -1.18 | 7.43E-09 |
| SMU_985 | bglA | putative beta-glucosidase | -1.18 | 9.05E-10 |
| SMU_714 |  | translation elongation factor EF-Tu | -1.18 | 7.28E-18 |
| SMU_1029 |  | conserved hypothetical protein | -1.18 | 4.83E-01 |
| SMU_1968c |  | conserved hypothetical protein | -1.19 | 4.85E-07 |
| SMU_2036 | pepO | putative peptidase | -1.19 | 3.88E-13 |
| SMU_1878 | ptnC | putative PTS system, mannose-specific component IIC | -1.19 | 2.81E-09 |
| SMU_589 |  | putative DNA-binding protein | -1.20 | 1.33E-12 |
| SMU_924 | tpx | thiol peroxidase | -1.21 | 5.62E-04 |
| SMU_1820c |  | putative glutamyl-tRNA(Gln) amidotransferase A subunit | -1.22 | 1.03E-19 |
| SMU_166 |  | hypothetical protein | -1.22 | 8.07E-08 |
| SMU_132 |  | putative hippurate amidohydrolase | -1.22 | 2.49E-03 |
| SMU_102 |  | putative PTS system, IID component | -1.23 | 9.01E-06 |
| SMU_1268 | hisB | putative imidazoleglycerol-phosphate dehydratase | -1.24 | 2.44E-12 |
| SMU_1296 |  | putative glutathione S-transferase | -1.24 | 1.25E-05 |
| SMU_675 |  | phosphoenolpyruvate:sugar phosphotransferase system enzyme I, PTS system EI component | -1.24 | 8.42E-17 |
| SMU_669c |  | putative glutaredoxin | -1.25 | 1.42E-05 |
| SMU_1270 | hisD | putative histidinol dehydrogenase | -1.25 | 2.02E-14 |
| SMU_2118 | opuCc | putative ABC transporter; osmoprotectant-binding protein, glycine betaine/carnitine/choline ABC transporter | -1.25 | 1.89E-17 |
| SMU_1805 |  | putative transcriptional regulator | -1.26 | 8.04E-08 |
| SMU_765 |  | NADH oxidase/alkyl hydroperoxidase reductase peroxide-forming | -1.28 | 4.06E-07 |
| SMU_1273 | hisC | putative histidinol-phosphate aminotransferase | -1.28 | 2.08E-11 |
| SMU_2127 |  | putative succinate semialdehyde dehydrogenase | -1.30 | 1.50E-07 |
| SMU_1894c |  | conserved hypothetical protein | -1.30 | 3.46E-02 |
| SMU_404c |  | hypothetical protein | -1.30 | 4.65E-05 |
| SMU_474 | luxS | putative autoinducer-2 production protein LuxS | -1.31 | 8.43E-11 |
| SMU_1463c |  | conserved hypothetical protein | -1.31 | 8.58E-17 |
| SMU_1599 | celR | putative transcriptional regulator; possible antiterminator | -1.33 | 1.82E-02 |
| SMU_1533 | atpG | FoF1 membrane-bound proton-translocating ATPase, a subunit | -1.33 | 5.70E-15 |
| SMU_128 | adhB | putative acetoin dehydrogenase (TPP-dependent), E1 component beta subunit | -1.33 | 8.71E-04 |
| SMU_767 |  | putative transposase, ISSmu1 | -1.35 | 4.20E-04 |
| SMU_1091 | wapE | hypothetical protein; possible cell wall protein, WapE | -1.35 | 2.23E-12 |
| SMU_1271 | hisG | putative ATP phosphoribosyltransferase | -1.35 | 3.28E-13 |
| SMU_1127 | rs20 | putative 30S ribosomal protein S20 | -1.37 | 7.70E-09 |
| SMU_496 | cysK | putative cysteine synthetase A; O-acetylserine lyase | -1.37 | 5.27E-06 |
| SMU_1272 | hisZ | putative histidyl-tRNA synthetase | -1.37 | 3.28E-13 |
| SMU_1408c |  | conserved hypothetical protein | -1.40 | 2.69E-02 |
| SMU_99 | fbaA | fructose-1,6-biphosphate aldolase | -1.40 | 3.17E-21 |
| SMU_1569 | malF | putative maltose/maltodextrin ABC transporter, permease protein MalF | -1.42 | 5.11E-13 |
| SMU_89c |  | putative nitrite transporter | -1.42 | 5.07E-09 |
| SMU_2119 | opuCd | putative osmoprotectant ABC transporter; permease protein | -1.42 | 3.28E-19 |
| SMU_129 | adhC | putative dihydrolipoamide acetyltransferase | -1.42 | 1.21E-03 |
| SMU_987 | wapA | cell wall-associated protein precursor WapA | -1.42 | 4.55E-25 |
| SMU_1570 | malG | putative maltose/maltodextrin ABC transporter, MalG permease | -1.45 | 7.75E-15 |
| SMU_405c |  | putative transcriptional regulator | -1.45 | 8.94E-05 |
| SMU_21 | mreD | putative cell shape-determining protein MreD | -1.45 | 5.84E-05 |
| SMU_1322 | budC | putative acetoin dehydrogenase | -1.45 | 1.19E-15 |
| SMU_609 |  | putative 40K cell wall protein precursor | -1.47 | 5.86E-04 |
| SMU_1600 | ptcB | putative PTS system, cellobiose-specific IIB component | -1.48 | 1.76E-02 |
| SMU_667 | nrdG | putative ribonucleotide reductase, small subunit | -1.48 | 1.44E-09 |
| SMU_840c |  | hypothetical protein | -1.48 | 8.43E-11 |
| SMU_1861c |  | hypothetical protein | -1.50 | 1.70E-06 |
| SMU_1869 | trxA | putative thioredoxin | -1.50 | 7.56E-07 |
| SMU_101 |  | putative sorbose PTS system, IIC component | -1.51 | 7.13E-06 |
| SMU_1877 | ptnA | putative PTS system, mannose-specific component IIAB | -1.52 | 6.39E-13 |
| SMU_1185 | mtlA1 | PTS system, mannitol-specific enzyme IIBC component | -1.52 | 6.74E-05 |
| SMU_1262c |  | hypothetical protein | -1.52 | 2.25E-08 |
| SMU_130 | adhD | putative dihydrolipoamide dehydrogenase | -1.53 | 1.27E-04 |
| SMU_613 |  | hypothetical protein | -1.55 | 5.14E-08 |
| SMU_574c |  | putative membrane protein | -1.58 | 1.79E-01 |
| SMU_1267c |  | hypothetical protein | -1.59 | 1.54E-18 |
| SMU_37 | purH | putative phosphoribosylaminoimidazolecarboxamide formyltransferase/IMP cyclohydrolase | -1.59 | 6.42E-13 |
| SMU_668c |  | ribonucleotide reductase, large subunit | -1.60 | 6.05E-11 |
| SMU_360 | gapC | extracellular glyceraldehyde-3-phosphate dehydrogenase | -1.60 | 5.88E-38 |
| SMU_1247 | eno | putative enolase | -1.66 | 3.15E-35 |
| SMU_1568 | malX | putative maltose/maltodextrin ABC transporter, sugar-binding protein MalX | -1.72 | 7.24E-12 |
| SMU_500 |  | putative ribosome-associated protein | -1.72 | 9.50E-12 |
| SMU_402 | pfl | pyruvate formate-lyase | -1.74 | 1.45E-13 |
| SMU_540 | dpr | peroxide resistance protein Dpr | -1.77 | 1.57E-15 |
| SMU_1867c |  | putative alcohol dehydrogenase | -1.89 | 4.85E-13 |
| SMU_183 | sloB | putative Mn/Zn ABC transporter | -1.92 | 2.02E-09 |
| SMU_179 |  | conserved hypothetical protein | -1.98 | 1.36E-05 |
| SMU_870 |  | putative transcriptional regulator of sugar metabolism | -2.02 | 3.13E-13 |
| SMU_1184c |  | putative transcriptional regulator, antiterminator | -2.03 | 5.06E-08 |
| SMU_866 |  | conserved hypothetical protein | -2.03 | 2.89E-31 |
| SMU_106c |  | putative transposase fragment | -2.07 | 2.34E-25 |
| SMU_503c |  | hypothetical protein | -2.11 | 1.96E-16 |
| SMU_2028 | sacB | levansucrase precursor; beta-D-fructosyltransferase | -2.15 | 1.08E-42 |
| SMU_1407c |  | putative transposase, ISSmu1 | -2.28 | 3.38E-05 |
| SMU_1425 | clpB | putative Clp proteinase, ATP-binding subunit ClpB | -2.34 | 1.99E-07 |
| SMU_1424 | pdhD | putative dihydrolipoamide dehydrogenase | -2.59 | 1.04E-04 |
| SMU_1790c |  | putative transcriptional regulator | -2.61 | 2.04E-40 |
| SMU_1794c |  | hypothetical protein | -2.93 | 2.10E-35 |
| SMU_182 | sloA | putative ABC transporter, ATP-binding protein; possible iron and/or manganese ABC transport system | -2.93 | 7.50E-11 |
| SMU_100 |  | putative sorbose PTS system, IIB component | -3.00 | 3.81E-13 |
| SMU_1030 |  | putative polyribonucleotide nucleotidyltransferase; Tn916 ORF8-like | -3.10 | 1.92E-01 |
| SMU_1791c |  | conserved hypothetical protein | -3.67 | 5.71E-74 |
| SMU_1801c |  | putative GTP-binding protein | -3.68 | 1.24E-102 |
| SMU_575c |  | putative membrane protein | -3.73 | 3.25E-04 |
| SMU_1348c |  | putative ABC transporter, ATP-binding protein | -3.76 | 2.38E-31 |
| SMU_1347c |  | conserved hypothetical protein; possible permease | -3.85 | 1.36E-20 |
| SMU_1365c |  | hypothetical protein; possible permease | -4.16 | 4.35E-43 |
| SMU_1358 |  | putative transposase fragment | -4.22 | 4.18E-02 |
| SMU_1795c |  | conserved hypothetical protein | -4.36 | 5.40E-78 |
| SMU_1797c |  | conserved hypothetical protein | -4.42 | 4.92E-73 |
| SMU_1792c |  | hypothetical protein | -4.61 | 8.89E-72 |
| SMU_1798c |  | conserved hypothetical protein | -4.83 | 1.38E-151 |
| SMU_1799 | nadD | putative nicotinate mononucleotide adenylyltransferase | -5.35 | 3.00E-191 |
| SMU_1800c |  | conserved hypothetical protein | -5.58 | 4.32E-106 |
| SMU_1366c |  | putative ABC transporter; ATP-binding protein | -6.99 | 4.52E-62 |

**Table S8 Differential gene expression (RNA-seq) analysis of *S. mutans* UA159 during CRISPRi-mediated knockdown of SMU_1802c**

| **Locus tag** | **Gene** | **Description** | **log2 fold change** | **FDR** |
| --- | --- | --- | --- | --- |
| SMU_1405c | cas9 | conserved hypothetical protein | 7.40 | 5.21E-153 |
| SMU_431 |  | putative ABC transporter, ATP-binding protein | 4.28 | 5.47E-81 |
| SMU_432 |  | putative ABC transporter, integral membrane protein | 4.18 | 6.34E-118 |
| SMU_1854 |  | conserved hypothetical protein | 2.04 | 2.16E-19 |
| SMU_200c |  | hypothetical protein | 1.84 | 1.20E-02 |
| SMU_490 | pflC | putative pyruvate formate-lyase activating enzyme | 1.81 | 2.93E-28 |
| SMU_213c |  | hypothetical protein | 1.77 | 8.80E-03 |
| SMU_215c |  | hypothetical protein | 1.71 | 1.29E-02 |
| SMU_206c |  | hypothetical protein | 1.71 | 4.62E-03 |
| SMU_216c |  | hypothetical protein | 1.65 | 1.97E-02 |
| SMU_204c |  | hypothetical protein | 1.61 | 8.42E-03 |
| SMU_202c |  | hypothetical protein | 1.61 | 1.03E-02 |
| SMU_197c |  | hypothetical protein | 1.61 | 6.37E-03 |
| SMU_196c |  | putative transfer protein | 1.59 | 1.24E-02 |
| SMU_207c |  | putative transposon protein | 1.58 | 4.56E-03 |
| SMU_433 |  | putative transcriptional regulator | 1.56 | 2.38E-25 |
| SMU_211c |  | hypothetical protein | 1.54 | 1.74E-02 |
| SMU_198c |  | putative conjugative transposon protein | 1.50 | 1.20E-02 |
| SMU_493 | pfl2 | formate acetyltransferase (pyruvate formate-lyase 2) | 1.47 | 8.38E-28 |
| SMU_434 |  | hypothetical protein | 1.46 | 2.14E-16 |
| SMU_208c |  | putative transposon protein; possible DNA segregation ATPase | 1.45 | 9.66E-03 |
| SMU_191c |  | putative integrase | 1.44 | 2.95E-02 |
| SMU_189 |  | hypothetical protein | 1.43 | 4.99E-02 |
| SMU_201c |  | putative transposon protein | 1.42 | 1.72E-02 |
| SMU_209c |  | hypothetical protein | 1.42 | 1.63E-02 |
| SMU_199c |  | hypothetical protein | 1.38 | 1.59E-02 |
| SMU_210c |  | hypothetical protein | 1.37 | 2.05E-02 |
| SMU_205c |  | hypothetical protein | 1.35 | 3.46E-02 |
| SMU_494 |  | putative transaldolase | 1.34 | 1.04E-20 |
| SMU_194c |  | conserved hypothetical protein; Bacteriophage P2 associated | 1.28 | 7.55E-02 |
| SMU_495 | gldA | glycerol dehydrogenase | 1.27 | 1.84E-19 |
| SMU_2126c |  | putative purine-nucleoside phosphorylase | 1.23 | 1.00E-13 |
| SMU_1762c |  | conserved hypothetical protein | 1.23 | 7.37E-04 |
| SMU_294 |  | conserved hypothetical protein | 1.21 | 1.35E-09 |
| SMU_195c |  | hypothetical protein | 1.20 | 3.16E-02 |
| SMU_1855 |  | hypothetical protein | 1.20 | 6.04E-07 |
| SMU_2080 |  | conserved hypothetical protein | 1.03 | 1.41E-02 |
| SMU_1378 |  | hypothetical protein | 1.03 | 1.02E-02 |
| SMU_789 |  | conserved hypothetical protein | 1.00 | 2.67E-12 |
| SMU_667 | nrdG | putative ribonucleotide reductase, small subunit | -1.02 | 2.12E-04 |
| SMU_1803c |  | hypothetical protein | -1.04 | 2.54E-06 |
| SMU_1091 | wapE | hypothetical protein; possible cell wall protein, WapE | -1.06 | 1.99E-07 |
| SMU_1954 | groEL | putative chaperonin GroEL | -1.07 | 2.27E-07 |
| SMU_1554c |  | hypothetical protein | -1.12 | 1.14E-02 |
| SMU_566c |  | conserved hypothetical protein | -1.12 | 4.38E-01 |
| SMU_1955 | groES | putative co-chaperonin GroES | -1.15 | 4.35E-05 |
| SMU_104 |  | putative alpha-glucosidase; glycosyl hydrolase | -1.17 | 1.81E-05 |
| SMU_668c |  | ribonucleotide reductase, large subunit | -1.18 | 8.59E-06 |
| SMU_1575c |  | hypothetical protein | -1.22 | 1.97E-02 |
| SMU_103 |  | putative PTS system, IIA component | -1.26 | 1.91E-05 |
| SMU_102 |  | putative PTS system, IID component | -1.28 | 1.81E-05 |
| SMU_669c |  | putative glutaredoxin | -1.30 | 3.66E-05 |
| SMU_545 |  | hypothetical protein | -1.32 | 4.58E-03 |
| SMU_11 |  | conserved hypothetical protein | -1.37 | 7.12E-02 |
| SMU_750c |  | hypothetical protein | -1.40 | 1.16E-02 |
| SMU_101 |  | putative sorbose PTS system, IIC component | -1.40 | 1.40E-04 |
| SMU_541 |  | conserved hypothetical protein | -1.41 | 4.55E-05 |
| SMU_1116c |  | hypothetical protein | -1.49 | 1.67E-03 |
| SMU_1657c |  | putative nitrogen regulatory protein PII | -1.63 | 2.13E-02 |
| SMU_100 |  | putative sorbose PTS system, IIB component | -1.64 | 2.24E-04 |
| SMU_1117 | naoX | NADH oxidase (H2O-forming) | -1.77 | 4.09E-04 |
| SMU_1396 | gbpC | glucan-binding protein C, GbpC | -1.88 | 3.35E-14 |
| SMU_1658 | nrgA | putative ammonium transporter, NrgA protein | -1.88 | 6.42E-03 |
| SMU_673 |  | conserved hypothetical protein | -1.91 | 5.67E-03 |
| SMU_670 | citB | aconitate hydratase; aconitase | -2.04 | 4.72E-03 |
| SMU_1004 | gtfB | glucosyltransferase-I | -2.04 | 1.29E-15 |
| SMU_1804c |  | hypothetical protein | -2.11 | 9.26E-04 |
| SMU_672 | idh | isocitrate dehydrogenase | -2.26 | 3.18E-03 |
| SMU_671 | citZ | citrate synthase | -2.29 | 2.70E-03 |
| SMU_1395c |  | hypothetical protein | -2.50 | 3.48E-13 |
| SMU_1790c |  | putative transcriptional regulator | -2.57 | 7.60E-52 |
| SMU_1927 |  | putative ABC transporter, ATP-binding protein | -2.98 | 1.26E-11 |
| SMU_1791c |  | conserved hypothetical protein | -3.07 | 6.55E-68 |
| SMU_1794c |  | hypothetical protein | -3.11 | 1.30E-52 |
| SMU_1928 | psaB | putative ABC transporter, permease protein | -3.32 | 2.80E-14 |
| SMU_1792c |  | hypothetical protein | -3.35 | 1.61E-72 |
| SMU_1795c |  | conserved hypothetical protein | -3.55 | 5.04E-78 |
| SMU_1798c |  | conserved hypothetical protein | -3.64 | 1.06E-132 |
| SMU_1797c |  | conserved hypothetical protein | -3.69 | 1.71E-76 |
| SMU_1799 | nadD | putative nicotinate mononucleotide adenylyltransferase | -3.77 | 9.55E-144 |
| SMU_1800c |  | conserved hypothetical protein | -3.85 | 7.02E-90 |
| SMU_1801c |  | putative GTP-binding protein | -3.94 | 1.10E-122 |
| SMU_1802c |  | conserved hypothetical protein | -4.12 | 1.46E-97 |

**Table S9 Differential protein abundance (TMT-MS/MS) analysis of *S. mutans* UA159 following CRISPRi-mediated knockdown of SMU_393**

| **Protein** | **Description** | **log2 fold change** | **FDR** |
| --- | --- | --- | --- |
| SMU_393 | Regulator of chromosome segregation-like C-terminal domain-containing protein | -3.99 | 2.07E-08 |
| DinB | DNA polymerase IV | -1.31 | 3.28E-05 |
| Acn | Aconitate hydratase A | -1.26 | 7.15E-05 |
| Icd | Isocitrate dehydrogenase [NADP] | -1.24 | 2.76E-04 |
| AdhE | Aldehyde-alcohol dehydrogenase | -1.21 | 3.50E-07 |
| SMU_58 | Apea-like HEPN domain-containing protein | -1.17 | 1.86E-07 |
| PurF | Amidophosphoribosyltransferase | -1.14 | 2.87E-06 |
| AdhD | Dihydrolipoyl dehydrogenase | -1.07 | 7.23E-05 |
| GlnA | Glutamine synthetase I alpha | -1.05 | 3.56E-05 |
| GltA | Glutamate synthase (Large subunit) | -1.05 | 1.00E-05 |
| SMU_104 | Alpha-glucosidase glycosyl hydrolase | -1.03 | 1.15E-07 |
| WapE | Gram-positive cocci surface proteins LPxTG domain-containing protein | 1.37 | 2.87E-06 |
| SMU_933 | Amino acid ABC transporter, periplasmic amino acid-binding protein | 1.39 | 4.73E-03 |
| GbpD | triacylglycerol lipase | 1.5 | 9.40E-06 |
| SMU_1213c | 5-nucleotidase | 1.64 | 2.12E-05 |
| GtfD | Glucosyltransferase-S | 1.98 | 1.81E-05 |
| ComC | Competence stimulating peptide | 2.55 | 5.50E-05 |
| Cas9 | CRISPR-associated endonuclease Cas9 | 7.42 | 1.38E-07 |

**Table S10 Strains and plasmids used in this study**

| **Strain** | **Description** | **Source** |
| --- | --- | --- |
| *S. mutans* strains |  |  |
| UA159 | Wild-type | Shields Lab |
| CRISPRi SMU_368 |  | (1) |
| CRISPRi SMU_369c |  | (1) |
| CRISPRi SMU_393 |  | (1) |
| CRISPRi SMU_415 |  | (1) |
| CRISPRi SMU_419 |  | (1) |
| CRISPRi SMU_1801c |  | (1) |
| CRISPRi SMU_1802c |  | This study |
| Δ*cas9* P*_xyl_*-d*cas9_Smu_* | Backbone for CRISPRi growth/microscopy studies | (1) |
| *dnaA*^Q197E^ |  | This study |
| CRISPRi SMU_393 *dnaA*^Q197E^ |  | This study |
| *E. coli* strains |  |  |
| 10-beta | Cloning host, derivative of DH10B | New England Biolabs |
| Plasmids |  |  |
| pPM_sgRNA | *S. mutans* integration vector containing a single-guide RNA cassette; kanamycin resistance | (1) |
| pALH124 | *E. coli* vector containing the *aphA3* cassette | (2) |
| pLacG-Em | Suicide integration plasmid carrying erythromycin resistance; inactivates the *lacG* gene required for growth on lactose via a single crossover recombination | (3) |

**Table S11 Oligonucleotides used in this study**

| **Oligonucleotide name** | **Sequence (5’ - 3’)** | **Experiment** |
| --- | --- | --- |
| CRISPRi1802F | ATCCCGCAACGTTTTAGAGCTAGAAATAGC | Depletion of SMU_1802c with CRISPRi |
| CRISPRi1802R | TGGCTGGACAACATTTATTGTACAACACG | Depletion of SMU_1802c with CRISPRi |
| dnaA_Q197E_A | TCCACAAGAAAAGATGCTAT | Site-directed mutagenesis of *dnaA* |
| dnaA_Q197E_B | CCATATTTTCTAGTCTAATGTG | Site-directed mutagenesis of *dnaA* |
| dnaA_Q197E_C | CACATTAGACTAGAAAATATGG | Site-directed mutagenesis of *dnaA* |
| dnaA_Q197E_D | CCAAATTCCTTTCCGATTTT | Site-directed mutagenesis of *dnaA* |
